# A trade-off mechanism underpins the evolution of a young two-gene sex-determining system in plants

**DOI:** 10.1101/2025.07.26.666942

**Authors:** Yingxiao Mai, Jiakun Zheng, Weijian Sun, Zheng Qin, Chengjie Chen, Wanchun Wei, Chaoqiong Song, Xingxing Liu, Xingling Su, Fengqi Wu, Qiuping Wu, Baoxuan Yao, Yanwei Hao, Yuanlong Liu, Zaohai Zeng, Jing Xu, Deborah Charlesworth, Rui Xia

## Abstract

Sexual systems in animals and plants are remarkably diverse, with dioecy having evolved independently in numerous lineages. In plants, dioecy often evolved more recently than in the best-studied animal systems, making plants especially important for understanding how separate sexes evolved independently from functionally hermaphrodite ancestors. Despite long-standing theories of developmental trade-offs in sex allocation, the underlying genetic mechanisms remain elusive. Here, we show that the XY sex determination system in the dioecious plant species *Eurycorymbus cavaleriei* in Sapindaceae involves two Y-linked mutations that act jointly within the developmental male-female trade-off: *YUNΔ*, a truncated allele that lowers the dosage of the D-class MADS-box gene *YUN*, and *SUN^MAO^*, a novel sRNA locus that silences the X-linked *SUN* allele. In females, SUN stabilizes the HD-ZIP transcription factor KUN, which is a known sex determinant in another dioecious plant, thereby promoting femaleness by increasing *YUN* expression; loss of *SUN* expression, together with the effect of *YUNΔ*, shifts development toward males. Two interlocking regulatory loops in this “SKY” module (SUN-KUN-YUN) fine-tunes *YUN* dosage. This dioecious system in *E. cavaleriei* likely evolved by sequential mutations in genes acting in the predicted male-female trade-off system, with their close linkage reflecting a translocation, and later recombination-suppressing inversions.

---

Most angiosperm species are functionally hermaphroditic (with each flower producing male and female organs, or individuals producing both male and female flowers, termed monoecy). As only around 6-8% of species are dioecious (with separate male and female plants), dioecy has clearly evolved independently from hermaphrodite or monoecious ancestors in different flowering plant groups^1,2^, most commonly from monoecy^3^. Darwin^4^ first considered why separate sexes might have evolved, and proposed that the loss of male or female function might be “compensated” by an increase in the other sex function; today this is termed a “trade-off”. Westergaard^2^ noted that genetic results from dioecious plants often suggest involvement of two closely linked dominant genes promoting maleness, one actively promoting male function (*M*), and the other (*SuF*) suppressing female functions. A theoretical model^5^ showed that dioecy could evolve by suitable mutations in two closely linked genes with such effects, with *SuF* factor promoting maleness by suppressing female functions, in a trade-off; the *SuF* effect therefore shows sexual antagonism, which creates selection for closer linkage of these two factors, leading to the evolution of a sex-linked region, and subsequently to the properties of sex chromosomes. The model proposed that an initial mutation creating a *SuF* factor changed the ancestral cosexual phenotype to one with less female, and consequently more male function, and that complete maleness often requires further mutations with similar effects. Y-linked *SuF* factors have now been identified in several plants^6^, but *M* candidate genes have been inferred in only two systems, kiwifruit (*Actinidia deliciosa*)^7^, and asparagus (*Asparagus officinalis*)^8^, though the latter identification has been questioned^9^.

Systems that evolved by two mutations can also create one-gene systems, leaving only the male-determining factor segregating, and thus detectable^10^, as in the persimmon (*Diospyros* species, in the family Ebenaceae)^11^. The persimmon male-determining factor, *OGI*, is a partial duplicate of an autosomal gene encoding a HD-ZIP (homeodomain-leucine zipper) transcription factor, *MeGI*; *OGI* produces small RNAs that repress *MeGI*, preventing its female-promoting activity^11^. As *MeGI* is involved in controlling both maleness and femaleness, it can be inferred that it probably involves a trade-off between the two sex functions^10^. In poplar species in the Salicaceae, a partial duplicate of a gene with an essential female function also leads to production of small RNAs that repress the autosomal progenitor^12^. Thus, a further difficulty in identifying plant sex-determining factors is that they are not necessarily protein-coding genes. Below, we describe a sex-determining system that differs from those already known, but which is supported by a set of related developmental observations and mechanistic analyses, integrated with evolutionary inferences.

## Flower sex differentiation in Sapindaceae

The Sapindaceae family, in the Sapindales order within the Malvid clade, includes around 140 genera (25 growing in China) and 2,000 species, some of which are economically important trees, including maple (*Acer* spp.), ackee (*Blighia sapida*), litchi (*Litchi chinensis*), horse chestnut (*Aesculus hippocastanum*), and yellowhorn (*Xanthoceras sorbifolium*). The flowers of all Sapindaceae species are unisexual, but differ in size and color, with some having petals and some not (Fig. 1A and B). Female flowers have a well-developed carpel with stamen rudiments, while male flowers have fully developed stamens and rudimentary carpels (Fig. 1B). Almost all Sapindaceae species are monoecious, with male and female flowers on the same plant, usually many more male than female flowers.

**Fig. 1.**
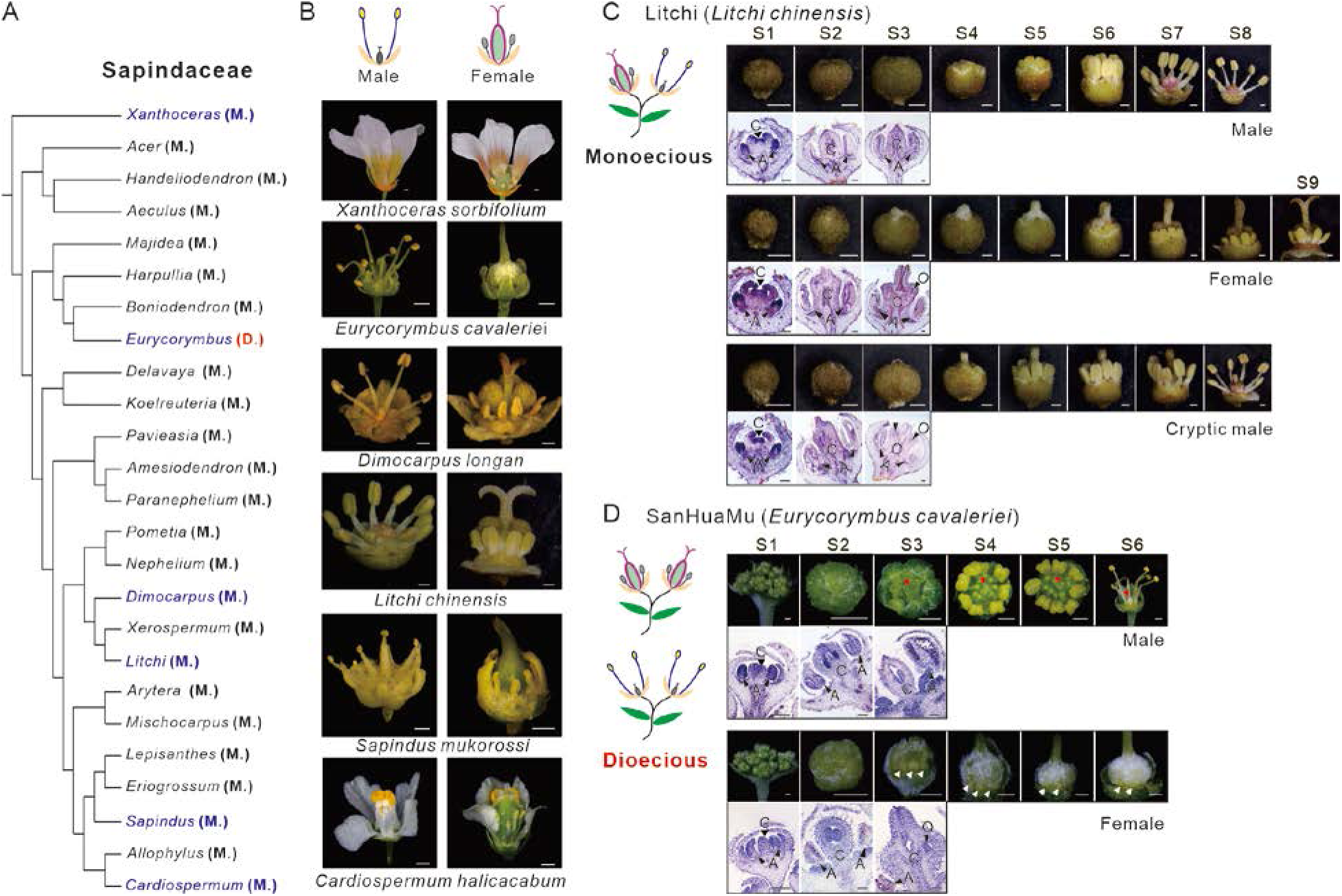
Similarity of flower sex differentiation in different Sapindaceae species. **(A)** The estimated phylogeny of the 25 genera of Sapindaceae species present in China. Monoecy is indicated by ‘M.’, and dioecy by ‘D.’. **(B)** Morphology of male and female flowers in six representative species. **(C)** Developmental stages of the three types of flowers described in the text (male, female, and cryptic male) in monoecious litchi. Developmental stages S1-S9 are described in detail in the Materials and Methods section. **(D)** The development of male and female flowers in dioecious *E. cavaleriei*, with the carpels in male flowers and anthers in female flower indicated by red and white arrows, respectively. Developmental stages S1-S6 are described in detail in the Materials and Methods section. Scale bars in the panels B-D of flower images denote 0.5 mm. Scale bars for images of paraffin sections represent 0.1 mm, and anthers, carpels and ovules are labeled “A”, “C”, and “O”, respectively.

In the monoecious litchi, each inflorescence produces both sexes of flowers, in an alternating sequence, with male flowers usually appearing first, followed by female ones (Fig. 1C). In some cultivars, such as ‘Feizixiao’, another flower type develops after each female phase. These have fully developed and functional stamens, but also an aborted and non-functional carpel that is larger than in normal male flowers (Fig. 1C); as they are functionally male, we termed them cryptic male flowers.

To identify genes involved in sex differentiation in the Sapindaceae, we first studied the process in litchi, which starts when the flower buds are about 1.00-1.50 mm in diameter (Fig.1C and S1A; stage 2). Comprehensive transcriptome profiling for all three types of flowers at multiple developmental stages (Fig. S2-S3) showed that several genes known to be essential for tapetum differentiation and function in plants, including orthologs of *GAMYB* (*MYB33*, *MYB65*) and several bHLH transcription factors, were expressed for longer periods than required in the abortive anthers of female flowers than in either type of male flower (Fig. S4). Similarly, the transcriptome of dissected carpels detected 424 genes preferentially expressed in well-developed carpels of female flowers, 22 of which are transcription factors (the Green module in Fig. S5). A few of these are known to function in flower organ development, including HD-ZIP, MYB, and ERF genes (Fig. S5), and one of them encodes a MADS-box protein, an ortholog of the Arabidopsis (*Arabidopsis thaliana*) *SEEDSTICK* (*STK*) gene, which is discussed below.

## The *Eurycorymbus* XY sex determination system

To investigate further these genes involved in controlling flower sexes in the Sapindaceae, we studied the strictly dioecious perennial tree, *Eurycorymbus cavaleriei* (Chinese name ‘SanHuaMu’, meaning tree with umbrella-like inflorescences; Fig. S6), the only species in the genus *Eurycorymbus.* As almost all Sapindaceae species are monoecious, we reasoned that its dioecy probably evolved from a monoecious ancestor, and thus sex-linkage might identify candidate genes for involvement in flower sex determination in both monoecious and dioecious species. Its flower sex phenotypes resemble those of litchi, with well-developed carpels in female flowers, and stamens in male flowers (Fig. 1D), suggesting a similar sex organ differentiation program in both species. However, flower sex differentiation in *E. cavaleriei* begins earlier, when the buds reach approximately 0.50-1.00 mm in diameter (Fig. 1D and S1B, after stage 1), and the carpel and anther rudiments in male and female flowers, respectively, are smaller than those of the corresponding sexes of litchi flowers.

We tested for a genetic sex-determining system in *E. cavaleriei* by searching for DNA sequence differences between male and female plants in the genomes of single male and female plants assembled from long-read PacBio sequences. Our haplotype-resolved (phased) reference assemblies for both sexes are around 280 Mb (Table S3-S7), with all contigs well anchored to the expected 13 chromosomes using Hi-C data (Fig. S7)^13^. Based on the phylogeny estimated by using single-copy orthologous genes, *E. cavaleriei* diverged from other Sapindaceae species around 40.6 million years ago (MYA, Fig. S8A and B). Neither *E. cavaleriei*, nor litchi or yellowhorn^14,15^, has undergone a recent whole-genome duplication (Fig. S8C). To identify the sex determining region (SDR), we sequenced a population of 78 male and 69 female plants from the natural population to discover single nucleotide polymorphisms (SNPs). A genome wide association study (GWAS) analysis, using flower sex as the phenotypic trait, detected a clear signal within the longer (left-hand) arm of the chromosome 2 assembly, in a region distant from the centromere (Fig. 2A and S9A). Analysis of linkage disequilibrium (LD) suggests that this sex-associated region on chromosome 2 has a lower recombination rate than the flanking regions (Fig. 2B and S9B). Sequence differentiation between the males and females in the natural population samples, estimated as *F*_ST_, is very high across most of this region (Fig. 2B and S9B).

**Fig. 2.**
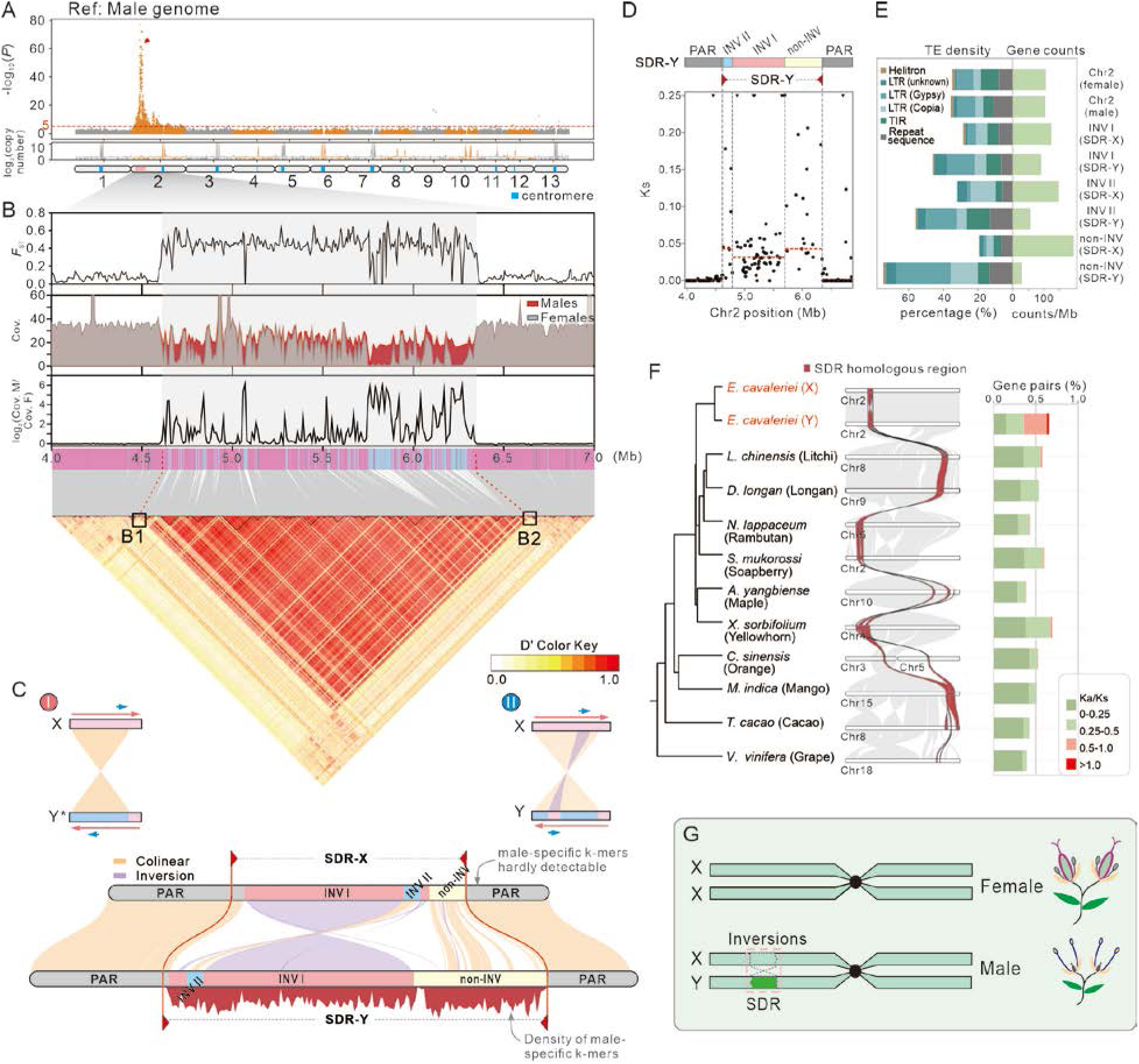
The sex-determining region (SDR) in the dioecious species *E. cavaleriei*. **(A)** Genome-wide association study (GWAS) analysis of plant sex using 147 accessions. The red dashed line indicates a p-value of 10^-5^. An association peak significantly associated with sex type is shown on chromosome 2. **(B)** Analyses of differentiation between males and females (quantified as *F*_ST_), coverage in females, males, and males versus females, and linkage disequilibrium (shown as D’), consistently located the SDR to the same chromosome 2 region, between 4.61 and 6.35 Mb in the male assembly. B1 and B2 represent the left and right boundaries of the SDR, respectively. Pink and blue lines indicate the presence or absence SNPs, respectively, and grey lines connect these SNPs to their corresponding positions in the heatmap. **(C)** Two inversions in the SDR-Y. The first, INV I, led to the formation of a proto-SDR-Y, and the second inversion (INV II) is within INV I. **(D)** Synonymous site divergence values (Ks) between alleles in the SDR-X and -Y, and in the flanking PARs (pseudoautosomal regions). **(E)** Transposable element (TE) densities (the percentages of assembly lengths of sequences attributed to TEs) and counts of gene numbers in 1 Mb windows in the fully sex-linked region (termed SDR), including its two inverted regions and its non-INV region, and in the rest of chromosome 2. **(F)** Synteny of genes in the SDR with the homologous regions of several eudicot species. The left panel shows the phylogenetic relationships of the species whose collinearity is shown in the middle panel, while the right panel shows Ka/Ks ratios for homologs of genes in reference to the SDR-X genes. **(G)** Model of chromosomal inversions resulting in the formation of the SDR.

The differentiated region coincides with a large paracentric inversion, INV I in Fig. 2C. As the arrangement in the female genome assembly is the same as in the outgroup species (Fig. S9C), we infer that the inversion occurred in the male-specific (Y) haplotype (Fig. 2C), and might be the cause of the lack of recombination in the region, as suggested for SDRs in other species with fully sex-linked regions^1,16^. Comparison of the X-linked haplotype (SDR-X, ∼1.05 Mb) with the corresponding region of the homologous chromosome (the ∼1.77 Mb SDR-Y) revealed a smaller inversion (INV II, 0.172 Mb) within INV I (Fig. 2C and S9D; details of the assembly of SDR regions are described in supplemental file 1). Within the SDR, the estimated synonymous site divergence values (Ks) between the X and Y haplotypes peak at around 3.6%, suggesting that X-Y recombination stopped about 4.3 MYA (substitution rate at 4.12 × 10^−9^, Fig. S9E and Table S8), consistent with recent evolution of the dioecious system. Higher X-Y Ks values were frequent in the non-INV region (Fig. 2D and S9F), suggesting that recombination was probably first suppressed in this region, initiating X-Y differentiation; the two inversions subsequently extended this non-recombining region, forming the present SDR. Gene densities are much lower in the Y-linked region than in the homologous X-linked one (Fig. 2E), consistent with the Y-linked region’s high transposon densities, which further supports complete sex-linkage of the entire region. Transposon densities are highest in the non-INV region (Fig. 2E), consistent with its higher Ks values. Genes in the SDR show collinearity with other eudicot genomes (Fig. 2F), suggesting that mutations in conserved genes within the region may have established sex-determining polymorphisms in *E. cavaleriei*. Taken together, the dioecious *E. cavaleriei* has an XY-linked genetic sex determination system, with male and female plants being the heterogametic “XY” and homogametic “XX” genotypes, respectively (Fig. 2G).

## *YUNΔ*, a Y-linked candidate *SuF* factor

The *E. cavaleriei* SDR region includes 147 annotated genes in total, with 128 located in the SDR-X, versus 132 in the SDR-Y, of which 113 are shared between the two (Table S9). The loss of only a few allelic genes is consistent with the small Ks values just described, as major genetic degeneration would be surprising in such a recently evolved and physically small Y-linked region. Expression analysis of six flower bud stages detected only a few genes in the SDR region with sex-biased expression, mostly with higher expression in females than males (Fig. 3A and Fig. S10). The gene with the largest sex difference in expression has a 114-fold higher total transcript abundance in females, and encodes a MADS-box transcription factor with strong homology to members of the AGAMOUS (AG) subfamily of MADS-box genes, especially to the AGL11 (SEEDSTICK, STK) protein in *A. thaliana* (Table S9 and Fig. S11).

**Fig. 3.**
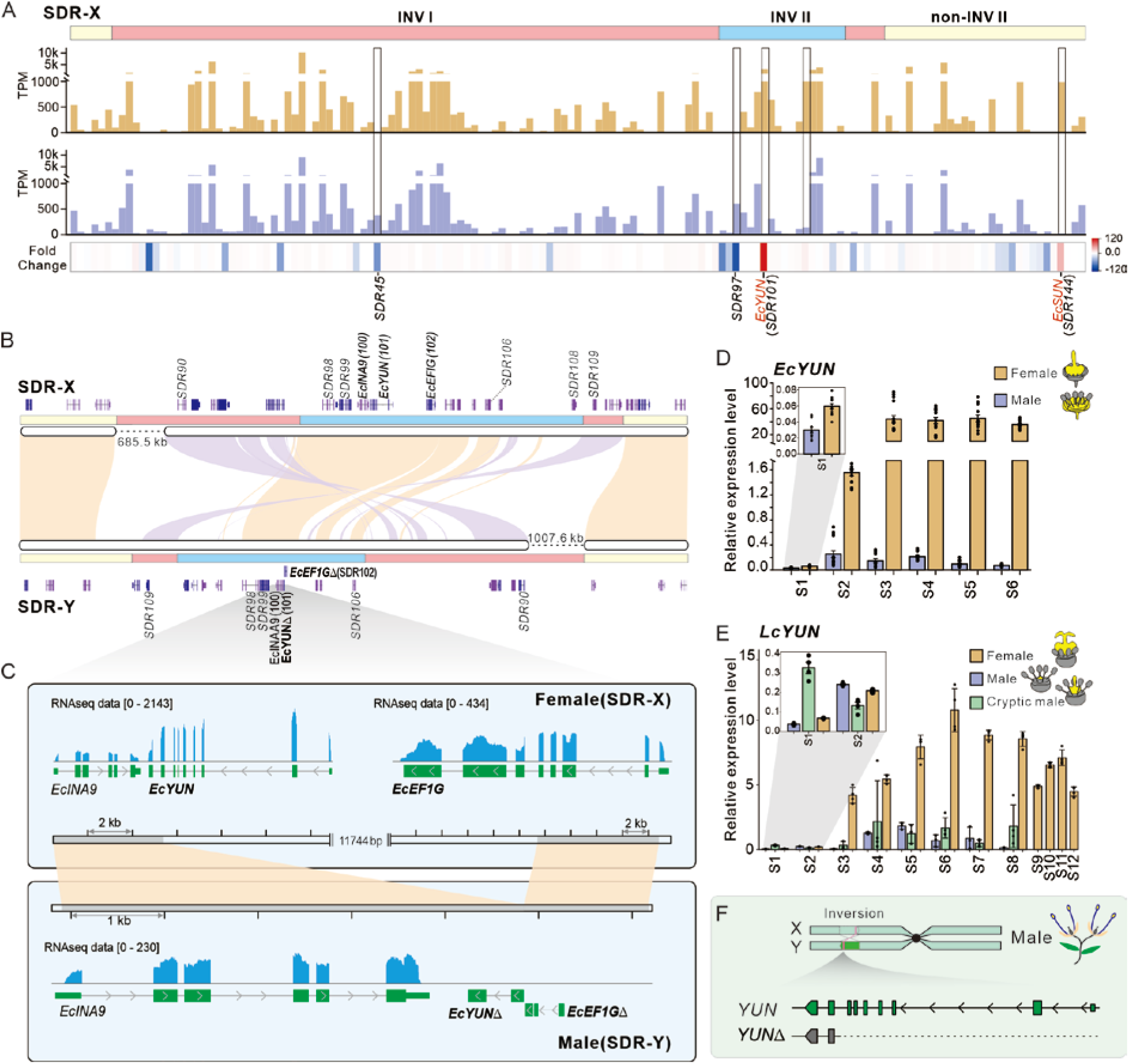
YUN involvement in sex determination in *E. cavaleriei*. **(A)** Total expression level (TPM) of genes in the SDR regions in female and male flower buds. Fold change was calculated as the ratio of the higher expression value to the lower one between female and male expression. Positive fold change values indicate higher expression in females, while negative values indicate higher expression in males. **(B)** The genes located within the INV II region of the SDR, including *EcYUN*. Orange shading denotes syntenic regions in forward orientation, while the purple shading indicates inversions (reverse orientation). **(C)** The large genomic deletion, around 29 kb long, in the SDR-Y that caused the *EcYUN* truncation. RNA-seq analysis of female flowers detected abundant reads from *EcYUN* and its neighboring SDR-X gene *EcEF1G*, but RNA-seq analysis of male flowers failed to detect any reads from the defective Y-linked *EcYUNΔ* and *EcEF1GΔ* genes. **(D) and (E)** Expression levels of *EcYUN* in flower buds of *E. cavaleriei* (D) and litchi (E). For S1-S2 stages, whole flower buds were used, and dissected carpels (highlighted in yellow) for later stages. Stages S1-S6 of *E. cavaleriei* correspond to those in Figure 1D, and the litchi stages S1-S9 are shown in Figure 1C. S10, S11, and S12 respectively represent the carpels of litchi female flowers at 3-, 6-, and 10-days post-pollination. Results shown are means ± standard error (SE, n ≥ 3). **(F)** The large truncation of the *YUN* gene in the SDR-Y, giving rise to the *YUNΔ* allele.

MADS-box genes are a class of transcription factors essential for plant growth and development. According to the “ABC model” initially proposed by Coen and Meyerowitz^17^ for flower development, and the later extended “ABCDE model”^18–21^, proteins encoded by a category of MADS-box genes termed MIKC-type specify the identity of different flower organs by forming tetrameric protein complexes^22,23^. Class A+E genes specify sepals, A+B+E petals, B+C+E stamens, C+E carpels and C+D+E specify ovules (reviewed in ^21^). *AGL11* encodes a type-II MADS-box protein of the D class, which are essential for ovule specification and seed development. Homozygotes for the *stk* mutant allele in *A. thaliana* show defects in ovule development, leading to reduced fruit size and seed number^20^. In the grapevine (*Vitis vinifera*), the *AGL11* ortholog, *VviAGL11*, controls fruit seedlessness^24,25^. In the tomato (*Solanum lycopersicum*), silencing of the orthologous gene *SlyAGL11* produces seedless fruits and the degree of seed development is proportional to transcript accumulation level^26^. This sex-linked MADS-box gene is therefore not an *M* factor candidate, but it is a candidate for a gene whose Y-linked allele in the dioecious species, *E. cavaleriei*, is mutated to promote maleness during the evolution of dioecy, such as an *SuF* factor. We named it ‘*YUN*’ (‘孕’ in Chinese, meaning “give birth to”), as it retains an ancestral female development function.

The X-linked *YUN* allele in *E. cavaleriei* is predicted to encode 225 amino acids in nine exons (Fig. S12A). Intriguingly, the Y-linked allele is an incomplete sequence (denoted as *YUNΔ*). It is within the small inversion, INV II, which has a ∼29 Kb deletion in the SDR-Y, relative to the X haplotype, causing the truncation of a large part of the *YUN* gene and half of the adjacent gene (Fig. 3B and 3C). The entire promoter region and the first seven exons are deleted, leaving only the last two exons; this SDR-Y *YUNΔ* allele remnant encodes only the final 37 amino acids, without a start codon or any complete functional domain (Fig. 3C, and Fig. S12A), making it unlikely to be transcribable and functional. Our allele-sensitive transcriptome profiling confirmed that almost no mRNA transcripts from the *YUNΔ* sequence is detectable (Table S10). The truncated Y-linked *YUNΔ* allele therefore seems unlikely to act by suppressing the complete, functional *YUN-X* copy in XY male plants.

As expected for a gene involved in ovule and carpel development, abundant *YUN* gene expression is detected only in the carpels of the flowers of female *E. cavaleriei* plants (Fig. 3D and S12B). *In situ* hybridization detected strong transcript signals in the two ovules of female flower buds in *E. cavaleriei* and litchi (Fig. S13). The early carpel abortion and the rudimentary carpel-like structures (Fig. 1D) are consistent with the low expression observed for the *YUN* gene in early male flower development (Fig. 3D). The expression pattern of the *YUN* ortholog in the monoecious litchi is also similar to that in the dioecious species, with much higher levels in the carpels of female flower buds, and specifically in ovules, than in male and cryptic male flowers (Fig. 3E, S12C and S13). The parallels in expression patterns between the monoecious litchi and dioecious *E. cavaleriei* support a conserved role of *YUN* in flower sex determination.

There is currently no transformation method for *E. cavaleriei* or litchi, and transformation experiments would be impracticable, as plants take several years to flower. We therefore tested the function of *YUN* in Arabidopsis, The *A. thaliana AGL11* (*SEEDSTICK*, *STK*) gene is a member of a MADS-box subfamily that also includes *AG* and two *SHATTERPROOF* genes, *SHP1* and *SHP2*. Although all four genes are required for normal ovule specification, *stk* single mutants produce shorter siliques with fewer and smaller seeds than wild-type plants^20^. Transformation of either *LcYUN* or *EcYUN* into Arabidopsis *stk* mutants rescued the defects in silique length, seed number, and seed length, consistent with an important role in ovule and seed development (Fig. S14 and S15). We therefore conclude that the *YUN* gene, a D-class MADS-box gene, has a conserved role in ovule specification and development and could be involved in flower sex differentiation in both the monoecious litchi and dioecious *E. cavaleriei*. As the truncated *YUNΔ* allele has the properties expected for a *SuF* factor in the dioecious species, we propose that this mutation could have contributed to *E. cavaleriei* SDR-Y heterozygous carriers being males (Fig. 3F).

## *YUN* dosage effects in the trade-off between male and female flower development

In *E. cavaleriei*, the phenotypic effects on flower sex of the Y-linked *YUNΔ* allele are dominant, which is surprising for a truncated loss-of-function mutation. Its effect cannot simply reflect Y degeneration involving loss of a functional copy from the Y-linked region, as the X-linked copy should still be expressed (in the absence of dosage compensation, its F/M expression should be about 2). Thus, a hypothetical additional Y-linked factor must be invoked to actively repress X-linked *YUN* expression in males (and possibly suppressed an ancestral Y-linked copy, making it dispensable). However, a search using a stringent pipeline identified no good candidate for such a factor (see below and refer to supplemental file 2 for details).

Alternatively, *YUN* could be a dosage-sensitive gene whose single wild-type *YUN* copy in males is insufficient to produce the wild-type female flower phenotype. Dosage sensitive effects also observed in some animal sex determination systems, including in *Drosophila* and *Caenorhabditis elegans* (as reviewed in ^27^), and two autosomal genes acting downstream of the mammalian Y-linked male-determining gene, *SRY*, *SF1* (Steroidogenic Factor 1) and *SOX9* (SRY-box 9)^28,29^. Consistent with this idea, the ortholog of *YUN* in oil palm (*Elaeis guineensis*), called the *SHELL* gene, is haploinsufficient as well, as heterozygotes with a single functional copy produce thinner fruit shells than the wild-type thick shell phenotype, while mutant homozygotes entirely lack the shell^30,31^. The known action of MADS-box proteins in quaternary complexes agrees with the observation that haploinsufficient genes often encode components of multimeric complexes or proteins that participate in highly connected nodes in regulatory or signalling pathways^32^. Such haploinsufficient properties are often conserved for homologous genes across different species ^33^. In Arabidopsis *STK* mutations also show haploinsufficiency in female organ development (Fig. S16). In the heterozygous genotypes, *SHP1^+/+^ SHP2^+/+^ STK^+/-^* and *shp1 shp2 STK^+/-^*, siliques were shorter, and contained less seeds, than the homozygotes, *SHP1 SHP2 STK^+/+^* (wild type) and *shp1 shp2 STK^+/+^*, respectively (Fig. S16C-E), but vegetative growth and stamen development were unaffected (S16A, B, and Fig. S17). The defects in female function were also greater in the *shp1 shp2 STK^+/-^* genotype, without wild-type copies of the other *SHP* genes, than in *SHP1 SHP2 STK^+/-^* individuals (Fig. S16C-E), indicating that dosage of other similar genes can ameliorate the mutant phenotype.

It is therefore plausible that *YUN-X* in heterozygous males is expressed normally, but at too low a level for sustainable ovule and carpel development after the earliest stages of ovule development. The vestigial carpels in male flowers of *E. cavaleriei* (Fig. 1D) and litchi (Fig. 1C) suggest that functional YUN proteins are present and lead to successful ovule and carpel initiation and perhaps early development. In flower buds approximately 0.50-1.00 mm in diameter (after stage S1), when flower sex differentiation begins (Fig. 1D and S1), *EcYUN* expression in heterozygous males was approximately half that in homozygous females (Fig. 3D) and, before the flower sex differentiation starts in litchi (at stage S2), *LcYUN* expression was similar in all three flower phenotypes, and the carpel sizes of male and cryptic male flowers are proportional to the *LcYUN* expression levels afterwards (Fig. 3E). These results suggest that dosage of the *YUN* gene is critical for the flower female organ differentiation in both species.

Haploinsufficient genes, for which low dosage in heterozygotes can have a dominant phenotypic effect, can sometimes also be toxic, having other phenotypic effects, when the gene product is overexpressed^34,35^. As *YUN* is highly expressed in females, we therefore asked whether this affects stamen development, which would suggest that it is part of a trade-off system between male and female structures. In *A. thaliana*, repressive effects of *AGL11* on stamens have indeed been observed in several studies in which *AGL11* orthologs were overexpressed^36–38^, and, in transgenic tomato plants overexpressing *SlAGL11*, the pollen was almost completely sterile when used to pollinate the pistils of WT plants^39^. However, these transformation experiments all used the CaMV 35S promoter, which is widely used for constitutive gene expression; thus, the observed repression might be due to detrimental ectopic expression of AGL11 protein in stamens, rather than reflecting its intrinsic function. We therefore re-tested the effect of an increased level of *STK* in *A. thaliana*, by introducing extra copy of the *STK* gene (including its whole genomic sequence and 3,534 bp of its native upstream promoter region, which should not lead to expression in stamens). These transgenic individuals again showed defective stamens, with short filaments and abnormal pollen grains. They produced shorter siliques with less seeds than control individuals (Fig. S18A-D, and J-K); these are effects of the defective pollen, as manual cross-pollination with WT pollen largely restored normal silique and seed development (Fig. S19). We also validated this anther-repressing effect of *NtYUN* in transgenic tobacco (*Nicotiana tabacum*) plants with additional *NtYUN* copies, which again had much shorter filaments and defective pollen grains, compared with control plants (Fig. S18E-G, and M-N). Thus, low dosage of *YUN* disrupts female organ development, and increased dosage has indirect pleiotropic effects on male ones, leading to the formation of large carpels and defective stamens, as in the female flowers in litchi. The *YUN* gene is therefore critical for the developmental trade-off between male and female organs in these plants.

## *SUN^MAO^*, a second Y-linked maleness-promoting factor

The non-recombining region in *E. cavaleriei* resulting from the genomic inversion is consistent with dioecy having evolved via the “two-gene” pathway, which predicts the presence of at least one (but probably often more than one) Y-linked suppressor of female function (*SuF*) closely linked to a gene whose X-linked allele differs from that in the cosexual ancestor, such as a maleness-promoting (*M*) gene^5^. As the Y-linked *YUNΔ* has the properties of a *SuF* factor, we searched for actively maleness-promoting candidates among the 132 *E. cavaleriei* SDR-Y genes mentioned above (Table S9). The stringent pipeline of gene expression analysis (detailed in Supplemental file 2) eliminated most genes; often their expression was negligible, or it did not differ significantly between the X- and Y-linked alleles. Of the 17 genes not definitively excluded by this criterion (Table S11), only two appear potentially related to reproduction: one encodes a protein homologous to gibberellin-regulated protein 9 (*GASA9*) and the other a zinc-finger homeodomain protein 2 (ZHD2). However, the protein sequence encoded by the SDR-Y allele of *GSAS9* lacks the critical functional domains necessary for its activity, and *ZHD2* is heterozygous in females rather than males and shows no significant male-specific expression. Therefore, although these protein-coding genes might contribute to sexual development in *E. cavaleriei*, none appears to be a strong maleness-promoting factor candidate. However, this does not exclude the possibility of a gene that promotes maleness by producing non-coding molecules, such as small RNAs (sRNAs).

We therefore investigated sRNAs in *E. cavaleriei* flower buds. We detected only a handful of sRNA-expressing loci within the SDR, mostly in intergenic regions and producing 24-nt sRNAs characteristic of heterochromatin (Table S12). Notably, however, a single locus within the non-INV region of the SDR-Y consistently produces abundant 21-nt sRNAs only in male flower buds and not female ones. These 21-nt sRNAs are generated from an inverted duplication in the SDR-Y, whose X-linked allele is a complete protein-coding sequence that we initially named *SDR144* (Figs. 3A and 4A). One arm of the Y-linked duplication is 2.26 kb long, and includes a 1.5 Kb promoter sequence and a ∼700 bp gene-body sequence of the *SDR144* gene; the profuse sRNAs are generated only from the latter region of the mutant SDR-Y allele, which is probably the only part transcribed (Fig. 4A and 4B). The X-linked *SDR144* allele is highly expressed in female flower buds, especially in well-developed carpels; however, like the *YUN* gene, its expression is extremely low in males (Fig. 4C), despite the presence of a complete allele in their SDR-X haplotype. This suggests silencing of the SDR-X full-length gene by sRNAs generated from the mutant SDR-Y allele, similar to the observation in the persimmon. This silencing effect was verified by co-infiltration into tobacco (*Nicotiana benthamiana*) leaves of the *SDR144* gene together with the sRNA-generating transcript (Fig. S20). To communicate the effect of the mutant allele, we renamed the *SDR144* gene ‘*SUN’* (SUN, “榫”, meaning tenon, a component of a mortise-tenon type of joint in carpentry) and the non-coding Y-linked mutant sRNA-generating allele that targets the *SUN* gene ‘*SUN^MAO^*’ (MAO, “卯”, means mortise).

**Fig. 4.**
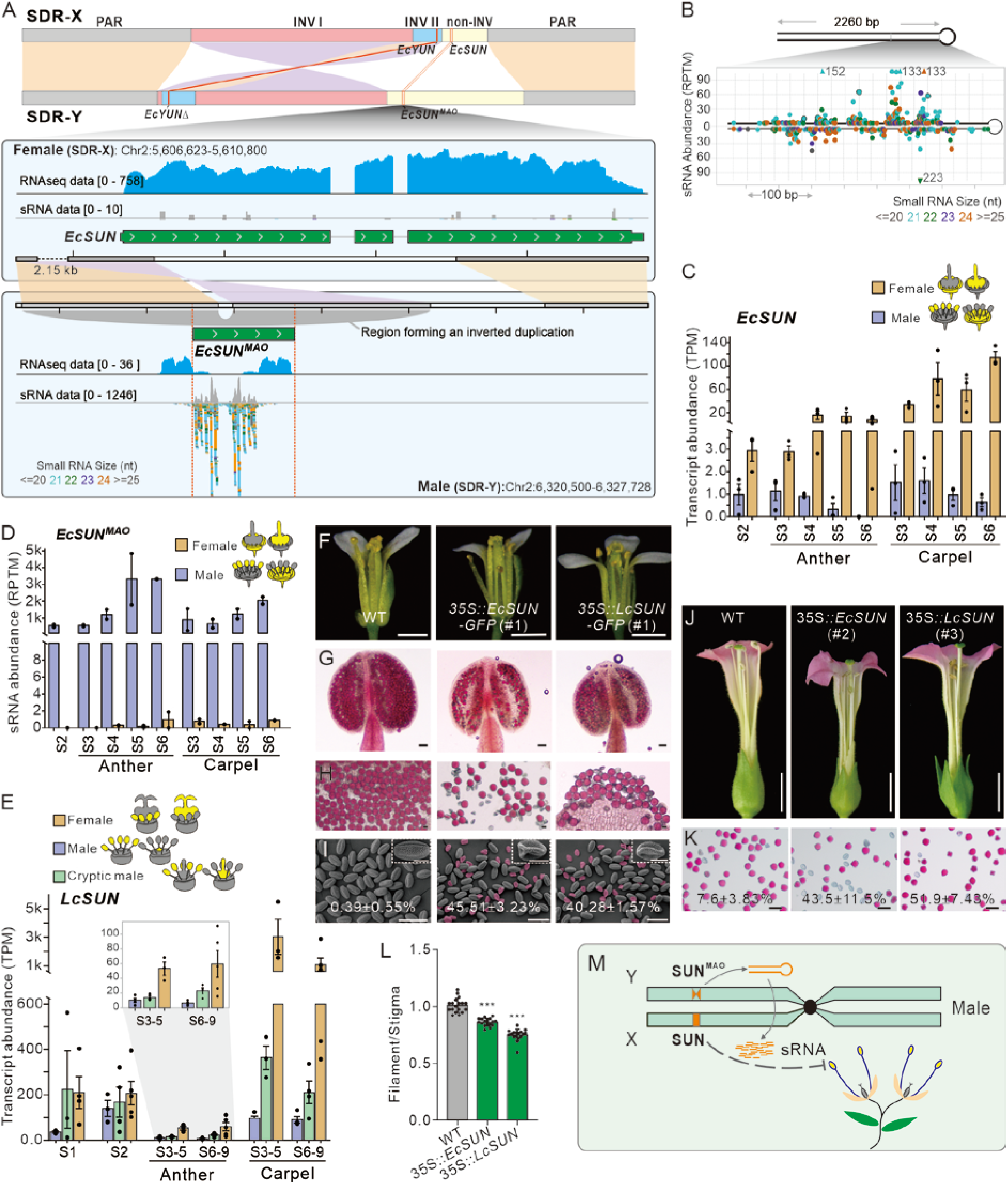
*SUN^MAO^* promotes male development by silencing the *SUN* allele. **(A)** An inverted duplication spanning the promoter and 5’-terminus of *EcSUN* (SDR144), located in the non-INV region of the SDR, created a novel sRNA-generating locus, *EcSUN^MAO^.* RNA-seq analysis detected profuse reads from *EcSUN* in female plants, but few in male plants for the *EcSUN^MAO^.* In contrast, abundant sRNAs were detected from the *EcSUN^MAO^* locus in male plants, while almost no sRNA reads were detected from the *EcSUN* locus in female plants. The invertedly duplicated region forming the stem-loop is indicated by a grey curve-bridge shape, and mapped sRNA reads are shown by color-coded short lines: 21-nt (light-blue, the predominant size), 22-nt (green), 23-nt (purple), 24-nt (orange), other sizes (grey). **(B)** The stem-loop structure into which *EcSUN^MAO^*locus can be folded to generate sRNAs of various sizes, as shown in the distribution; each dot shows a distinct sRNA detected in our experiment. sRNAs of different sizes are color-coded using the same scheme as in (A). **(C)** and **(D)** Transcript abundance of *EcSUN* (C) and the sRNA abundance of *EcSUN^MAO^* (D) in *E. cavaleriei* flower buds at different developmental stages (means ± standard errors, n ≥ 3). TPM, transcripts per million; RPTM, reads per ten million. **(E)** Transcript abundance of *LcSUN* in flower samples of litchi at different developmental stages (means ± standard errors, n = 3). In C-E, whole flower buds were used for stages S1-S2, and dissected organs for stages after S2. **(F)** Open flowers from the wild type (WT), the *35S::EcSUN-GFP* transgenic *A. thaliana* line 1 (#1), and the *35S::LcSUN-GFP* transgenic line 1 (#1). Bars = 1 mm. (**G)** and **(H)** Alexander staining of the anthers (G) and pollen grains (H) from WT, *35S::EcSUN-GFP* (#1) and *35S::LcSUN-GFP* (#1). Bars = 50 µm. Percentages of abnormal pollen grains are shown at the bottom. Results are means ± standard errors (n > 300). **(I)** SEM images of mature pollen grains from the WT, *35S::EcSUN-GFP* (#1) and *35S::LcSUN-GFP* (#1) lines. Abnormal pollen grains are highlighted in purple. Bars = 50 µm. The inset in the upper right corner shows an enlarged view of a pollen grain. Bars = 10 µm. (**J)** Open flowers from the WT, the *35S::EcSUN* transgenic tobacco line 2 (#2) and *35S::LcSUN* (#3). Bars = 1 cm. (**K)** Alexander staining of pollen grains from the WT, *35S::EcSUN* (#2) and *35S::LcSUN* (#3). Bars = 50 µm. Percentages of abnormal pollen grains was shown at the bottom. Results are means ± standard error (n > 200). (**L)** The ratio of filament length to stigma length in the WT, and in the transgenic tobacco plants, *35S::EcSUN* (#2), and *35S::LcSUN* (#3); means ± standard errors are shown. Significant differences (*p*) between the ratio of WT and those of *35S::EcSUN* (#2) and *35S::EcSUN* (#3) were assessed using Student’s *t*-tests and represented by asterisks (***, *P* < 0.001). **(M)** Diagram illustrating that the *SUN^MAO^* locus in SDR-Y can generate siRNAs that silence the *SUN* gene in the SDR-X.

The Y-linked sRNA-generating *SUN^MAO^*allele is therefore a good maleness-promoting gene candidate, while the X-linked *SUN* allele that can be silenced by *SUN^MAO^*-derived sRNAs must have the function of suppressing maleness in female plants. Interestingly, *SUN* encodes a homolog of the *A. thaliana SMAX1* gene (Suppressor of Max2 1)-like 7 (*SMXL7*), a transcriptional co-repressor in the strigolactone (SL) signaling pathway. SLs are a class of butanolide molecules produced by plant roots; they function as hormones that regulate various developmental processes and environmental responses, especially seed germination and shoot branching^40^. *SUN* is a member of the SMXL family that is known as *DWARF53* (*D53*) in rice, and includes the *SMXL6*, *SMXL7*, and *SMXL8* genes in Arabidopsis (Fig. S21). Upon perception of SLs by a receptor, the co-repressor SMXL protein is ubiquitinated and rapidly degraded by the 26S proteasome, and downstream transcriptional regulation is activated^40^. Expression of *EcSUN* in the rudimentary anthers of *E. cavaleriei* female flowers is much higher than in male flowers, consistent with a maleness-repressing role (Fig. 4C), while *SUN^MAO^*-derived sRNAs are produced exclusively in flower buds of male plants, aligning with their extremely low *EcSUN* expression (Fig. 4D). Similarly, expression of *LcSUN* in the monoecious litchi is high in carpels of female flowers, and is also higher in the abortive anthers of female than male flowers (Fig.4E).

To verify the maleness-suppressing function of *SUN* genes, we ectopically overexpressed them in Arabidopsis and tobacco (*N. tabacum*). As predicted, overexpression of either *EcSUN* or *LcSUN* resulted in defects in male reproductive organs (Fig.4F-4I). In the transgenic *A. thaliana* plants, defective pollen grains were observed alongside smaller anthers compared to control plants, with the overexpression of *EcSUN* and *LcSUN* specifically resulting in at least 40% pollen sterility (Fig 4G-I and S22). The defective effects in stamens were more pronounced in tobacco plants overexpressing the transgenes, with shortened filaments and higher proportions of abnormal pollen grains (Fig. 4J-4L and S23). We therefore conclude that the Y-linked *SUN^MAO^*could indeed be a factor promoting maleness by generating sRNAs that silence the expression of X-linked *SUN* allele in males, which otherwise acts to repress male functions in female flowers (Fig. 4M). The *SUN^MAO^* appears to have a *SuF* function, rather than being an M factor, since its X-linked allele (the *SUN* gene) is clearly not a loss-of-function mutation of a Y-linked gene.

## The SKY module of interlocking regulations

Although SUN functions as a maleness repressor, it is expressed more highly in carpels than anthers, especially in litchi (Fig. 4C and 4E). This unexpected expression pattern prompted us to investigate the underlying mechanism of *SUN’s* function. Therefore, we re-examined our expression data, to search for transcription factors whose expression correlates closely with the *SUN* expression (Fig. 5A and Fig. S24). Among the small number of such factors identified, a HD-ZIP transcription factor stands out, as it has high expression and tight correlations with *SUN* expression across all flower bud stages studied, in both *E. cavaleriei* (R=0.82) and litchi (R=0.89; Fig. 5A; Fig. S25). We named it ‘*KUN*’ (‘坤’ in Chinese, which refers to the nurturing and life-supporting role of mothers or the earth). Our single-nucleus RNAseq analysis revealed *SUN* is not only tightly co-expressed with *KUN*, but their mRNA transcripts are also co-localized at the cellular level in the ovule and ovary regions (Fig. 5B-C and 5E-F, Fig. S26 and S27). This co-localization was further confirmed by RNA in situ hybridization (Fig. 5D, 5G and S27). Strikingly, *KUN* is an autosomal gene that is orthologous to the *MeGI* gene in persimmon (in the Ebenaceae), which participates in the genetic control of dioecy^11^, and to the barley (*Hordeum vulgare*) *Vrs1* gene which affects inflorescence development, and the *CmHB40* in the monoecious plant melon (*Cucurbita melo*), whose expression inhibits stamen development^41,42^ (Fig. S28). Good binding motifs (“AAT(T/A)ATT”) for this HD-ZIP protein are present in the promoters of *EcSUN* and *LcSUN* (Fig. 5H), and we confirmed that KUN proteins bind to both promoters by yeast-one-hybrid and electrophoretic mobility shift assays, and that this activates *SUN* transcription (Fig. 5H-J and S29), explaining their tight coexpression.

**Fig. 5.**
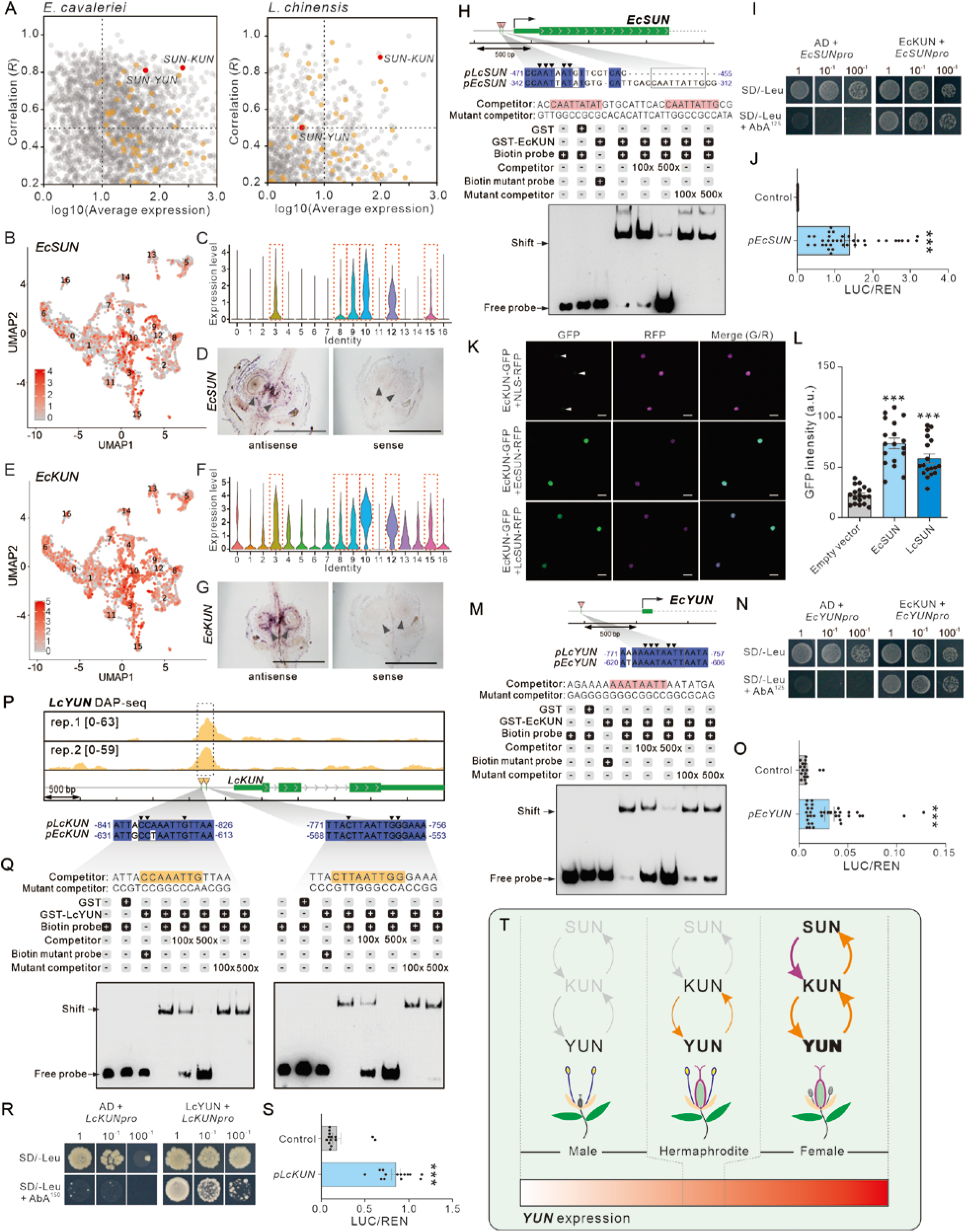
The SUN-KUN-YUN interlocking regulations. **(A)** Expression correlation analysis between candidate genes (see method for gene selection criteria) with *SUN* in *E. cavaleriei* and *L. chinensis*. The Y-axis represents Pearson correlation coefficients (*R*) of co-expression between candidate genes and *SUN*, while the X-axis shows the mean expression (log10(TPM)) of candidate genes. Orange dots denote transcription factors; correlation levels of *KUN*-*SUN* and *YUN*-*SUN* are highlighted by red dots. **(B-D, E-G)** Single nucleus RNAseq (B-C, E-F) and in-situ hybridization (D, G) indicates that *SUN* and *KUN* transcripts are co-localized at the cellular level. Feature plots (B, E) and violin plots (C, F) represent expression of *EcSUN* (B-C) and *EcKUN* (E-F) in 17 distinct groups identified by UMAP clustering of 6,115 nuclei. Both *EcSUN* and *EcKUN* are highly expressed in Clusters 3, 8, 9, 10, 12, and 15 (dotted red rectangles). Expression of *EcSUN* (D) and *EcKUN* (G) detected by RNA in situ hybridization assays on S4 female flower buds. The ovule positions are indicated by arrows. Bars = 0.5 mm. **(H, M)** Binding of EcKUN to the identified sites in the promoter of *EcSUN* (H) or *EcYUN* (M) in electrophoretic mobility shift assays (EMSA). Truncated EcKUN protein (containing the HD-ZIP I DNA-binding domain) was used in the assay. **(I, N)** Binding of EcKUN to the promoter of *EcSUN* (I) or *EcYUN* (N) in yeast one-hybrid (Y1H) assays. **(J, O)** Activation of the *EcSUN* (F) or *EcYUN* (L) expression by EcKUN in vivo in transient dual-luciferase reporter assays. Results shown are means ± standard error (SE, n >10). Asterisks represent significant differences between control and *EcSUNpro* treatments in Student’s *t* tests (***, *P* < 0.001). **(K)** Representative images form confocal laser scanning microscopy showing EcSUN or LcSUN enhances the protein stability of EcKUN. Tobacco cells were examined by green GFP fluorescence of EcKUN protein and red fluorescence of EcSUN and LcSUN protein. White arrows indicate attenuated nuclear signals of EcKUN-GFP in the absence of EcSUN or LcSUN protein. Scale bars = 20 μm. **(L)** Quantification of nuclear GFP intensity corresponding to (K). Results are means ± SD (n = 20). Asterisks represent significant differences between the empty vector control and EcSUN/LcSUN treatments in Student’s *t* tests (***, *P* < 0.001). **(P)** DAP-seq analysis of LcYUN identified a significant peak in the promoter region of *LcKUN* in two replicates. Two binding sites for LcYUN were detected in the promoter of *LcKUN*. **(Q)** Binding of LcYUN on the promoter of *LcKUN* in the EMSA assay. Truncated LcYUN protein (containing the MADS-box DNA-binding domain) was used. **(R)** Binding of LcYUN to the promoter of *LcKUN* in the Y1H assay. **(S)** Activation of the *LcKUN* expression by LcYUN in vivo in transient dual-luciferase reporter assays. Results are collected and shown similar to (H) and (K). **(T)** The SKY (SUN-KUN-YUN) model for how *YUN* expression is controlled in a developmental trade-off between male and female organs. In Sapindaceae, when the SUN-KUN-YUN circuit is not active, *YUN* expression is below the threshold required to support female development, leading to unisexual male flowers; when the *SUN* is active in carpels, the reinforcing SUN-KUN and KUN-YUN loops increase *YUN* expression, suppressing male development and resulting in unisexual female flowers. When the carpel-specific *SUN* expression is lacking (in species other than the Sapindaceae), the reciprocal KUN-YUN regulation maintains *YUN* expression at an intermediate level, yielding perfect (bisexual) flowers. Orange and purple arrows indicate transcriptional regulation and protein interaction, respectively.

We mentioned above that *SUN* copies in the Sapindaceae have high sequence similarity to SMXL/D53 genes. However, they have a mutation in a critical functional motif (discussed below), suggests a function different from that of other SMXL/D53 proteins. SMXL proteins have domain structures and key motifs resembling members of the caseinolytic peptidase (ClpB) ATPase family, which can function as chaperones to stabilize or correctly fold proteins^43^. Instead of responding to SL signals, SUN might therefore function as a chaperone protein. Given the tight co-expression and co-localization of *SUN* with *KUN* transcripts, we next investigated whether SUN and KUN protein could interact. Indeed, SUN interacts with KUN and, consistent with its proposed role as a chaperone, it enhances the stability of KUN proteins (Fig. 5K-L and Fig. S30). The maleness-suppressing function of *KUN* was confirmed by exogenous transformation in both Arabidopsis (Fig.S31) and tobacco (Fig. S32). In both litchi and *E. cavalerei*, we observed higher expression of the *KUN* genes in the developing stamens of female flowers (which later abort) than in those male flowers (Fig. S25), consistent with *KUN* acting as a maleness repressor, like the inferred effect of *MeGI* expression in the persimmon. Therefore, it is plausible that *SUN* represses maleness in female flowers, probably involving the *KUN* function.

In addition to the mutual co-expression observed between *KUN* and *SUN*, the *YUN* gene is also co-expressed with the *KUN* and *SUN* genes in both *E. cavaleriei* and litchi (Fig. 5A and Fig. S24), albeit with a lower correlation. This suggests that *YUN* may also be regulated by KUN. We identified binding motifs for KUN protein in the promoter of the *YUN* alleles of both species, and experimentally confirmed both its binding and activation of *YUN* expression (Fig. 5M-O and S32). To learn more about the trade-off system, including how high *YUN* dosage affects stamen development, we searched for potential targets of the YUN protein in litchi, using DNA affinity purification sequencing^44^. We identified two binding sites for the YUN protein in the promoter of the *LcKUN* (Fig. 5P). YUN protein could bind effectively to both sites to activate expression of the *KUN* gene (Fig. 5P-S and Fig. S33). Therefore, we conclude that *KUN* and *YUN* forms a mutual regulatory transcriptional loop and *YUN* affects anther development, probably by activating the *KUN* gene.

The three genes, *SUN*, *KUN* and *YUN*, are found in the genomes of all eight Sapindaceae species analysed, and their interactions described above are probably conserved across these species (Fig. S34 and S35). Therefore, flower sex determination in Sapindaceae is likely controlled by a regulatory network collectively termed the “SKY” module (*<u>S</u>UN*-*<u>K</u>UN*-*<u>Y</u>UN*). This consists of two interlocking regulatory loops: *SUN-KUN* and *KUN*-*YUN* (Fig. 5T). SUN enhances KUN activity by stabilizing the KUN protein; KUN, in turn, regulates *SUN* transcription; and *KUN* and *YUN* reciprocally modulate each other’s expression. These coupled loops control the expression of *YUN*, which is central to the male-female trade-off. In the SKY module, the *SUN*-*KUN* loop serves as an amplifier that stimulates the expression of the MADS-box *YUN* gene in females. When the amplifier is active, *YUN* expression is enhanced, then leading to the development of female flowers; when this amplification is absent, *YUN* levels remain low, resulting in the production of male flowers (Fig. 5T). This amplification is absent in species other than the Sapindaceae (for instance, the Rutaceae in the Sapindales order), in which *SUN* and *KUN* are not co-expressed (though orthologs of both genes are present); consequently, *KUN* and *YUN* expression permit development of hermaphrodite flowers.

## Evolution of dioecy in *E. cavaleriei*

The X-linked *YUN* allele in *E. cavaleriei* is orthologous to the *A. thaliana AGL11* gene (*STK*), whose mutants have defects in female functions^20^. Phylogenetic analysis of the AG-subfamily of MADS-box genes, which include the *STK* gene and two *SHP* genes in *A. thaliana*, shows that AGL11 clade genes are duplicated in several angiosperm lineages (Fig. S11); Interestingly, the *SHP* ortholog is completely absent in *E. cavaleriei*, and the monoecious litchi has only one *SHP* gene (Fig. S11). Therefore, *EcYUN* is probably an indispensable gene to fulfil the function of ovule specification, making it of central importance in developing female structures in *E. cavaleriei*. In *A. thaliana shp1 shp2 stk* triple mutants, normal ovule and seed development is completely disrupted, but male organ morphology and functions remain normal, transforming bisexual flowers into male (female-sterile) ones (Fig. S16 and S17).

Since most Sapindaceae species are monoecious, the dioecious state of *E. cavaleriei* probably evolved from a monoecious ancestor, and the two-gene model predicts the involvement of genes that were already closely linked^5^. In *E. cavaleriei*, *EcSUN* and *EcYUN* are indeed both within a single syntenic block of the chromosome homologous to litchi Chr8, the chromosome that carries the *LcYUN* gene, but in litchi, the *LcSUN* gene is on Chr 13 (Fig. 6A). The litchi state (with the *SUN* and *YUN* orthologs on different chromosomes) is shared by most Eudicotyledon species (Fig. 6A and S36). Close linkage before dioecy evolved would require that the *SUN* gene translocated to near the *YUN* gene on chromosome 2 in the lineage leading to *E. cavaleriei*. To test this hypothesis, we sequenced the genome of another Sapindaceae species in the subfamily to which *E. cavaleriei* belongs, Dodonaeoideae. This outgroup species, *Boniodendron minus*, has two *SUN* paralogs, suggesting that a duplicative transposition event occurred in a Dodonaeoideae ancestor, leaving one copy corresponding to the *LcSUN* on Chr 13 and the other in a location near the ancestral *YUN* gene on Chr2, so that *YUN* and *SUN* were already closely linked before *E. cavaleriei* evolved dioecy. Although *SUN* in these Sapindaceae species is the homolog of the Arabidopsis *SMXL* and rice *D53* genes, it is unlikely to function in the SL signalling pathway, because all *SUN* copies in the Sapindaceae species studied here have mutations in the “RGKT” motif (Fig. 6A and S37), which is reported to be crucial for the D53 protein in rice^45^.

**Fig. 6.**
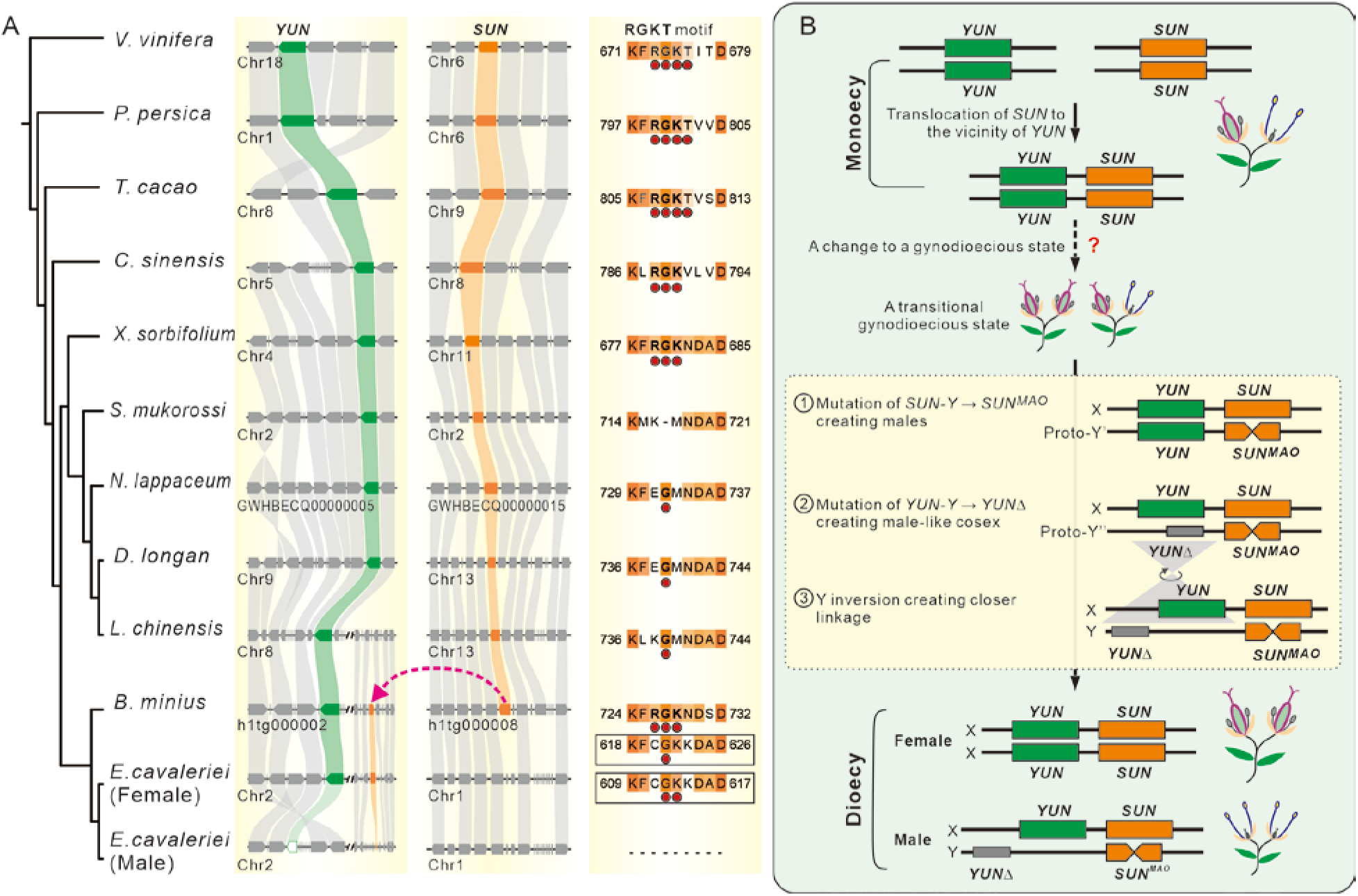
Evolution from monoecy to dioecy in Sapindaceae. **(A)** Comparison of syntenic regions carrying the *YUN* and *SUN* genes in eudicot species shown in the left-hand panel. The two middle panels show the results of collinearity analyses of the regions carrying the *YUN* and *SUN* genes, indicated in green and orange, respectively. The pink dashed arrow indicates the duplicative transposition of the *SUN* gene to the proximity of *YUN* in *B. minus*. The right-hand panel shows the RGKT motif amino acid sequences, with red dots indicating the essential residues. **(B)** Diagram illustrating the changes from monoecy to dioecy evolution in *E. cavaleriei*. The red question mark is to indicate the unknown changes leading to the appearance of a transient gynodioecious state.

Dioecy in *E. cavaleriei* probably evolved from monoecy, the state in other Sapindaceae. As explained above, both Y-linked factors, *YUNΔ* and *SUN^MAO^*, regulate flower sex via fine-tuning *YUN* expression, exerting dual roles in suppressing femaleness and promoting maleness via the trade-off mechanism, they both function as *SuF* factors, and neither has properties expected for an *M* factor (a gene essential for male function, whose X-linked allele is a non-functional mutant copy, creating females). As the X-linked *YUN* and *SUN* alleles are both functional wild-type copies of the gene, homozygotes would be expected to be monoecious, as in litchi. Our evolutionary hypothesis therefore proposes that one or both of these two genes likely evolved from their original state in the monoecious ancestor to acquire a female-promoting function. If females that arose in this way increased in frequency in the ancestral monoecious population, this would create a gynodioecious population with female and monoecious (cosexual) individuals, the predicted first step in the evolution of dioecy^5^. The presence of females favours enhanced maleness of the cosexual individuals, leading to the emergence of the Y-linked maleness-promoting alleles. The *SUN^MAO^* was probably the first maleness-promoting mutation, as it is in the non-INV region where recombination was probably first suppressed in the SDR-Y (Fig. 2D). *SUN^MAO^*silences the *EcSUN-X* allele, eliminating its enhancement of *YUN* expression via the SKY module. However, this silencing alone was probably insufficient to reduce the *YUN* dosage to the low levels required for a fully male phenotype, as both *YUN* alleles remained functional, so it probably created male-like individuals with enhanced male function while retaining some potential to occasionally express female function, an intermediate step in the transition to dioecy^5^. Selection would then favour the evolution of a fully male phenotype. A subsequent mutation in the Y-linked *YUN* copy to the non-functional *YUNΔ* allele would complete this transition, by reducing the genomic *YUN* dosage, transforming the male-like individuals into completely male individuals (Fig. 6B). Finally, the large inversion (INV I) created complete linkage between *YUNΔ* and *SUN^MAO^*, stabilizing the XY genetic sex determining system in *E. cavaleriei* (Fig. 6B).

The essential role of MADS-box genes in flower organ specification and development in plants is well established, and a critical role in plant sex determination has frequently been suggested for this gene family. However, our discovery that the *YUN* gene, a D-class MADS-box gene, is involved in sex determination in *E. cavaleriei*, is the first direct evidence for such involvement, though a role for *AGAMOUS* homologs is implied in *Ficus*^46^.

An important implication of this study is that Y-linked factors could act in sex determination in ways other than those previously proposed^5,10^. In *E. cavaleriei*, both Y-linked factors are mutant alleles of their X-linked counterparts (*YUNΔ* is a loss-of-function mutation, and *SUN^MAO^*, an allele producing non-coding sRNAs that silence *SUN*, can also be considered a functional knockout mutation). Sex determination systems are often categorized as “one-gene” or “two-gene” based on the number of actively male-determining Y-linked factors^6^. As *SUN^MAO^* is the only active Y-linked factor, *E. cavaleriei* might therefore be considered a “one-gene” system. Yet complete maleness also depends on *YUNΔ* mutation which reduces the genomic dosage of *YUN* in males, and affects the male-female trade-off. Both Y-linked mutations promote maleness, but, unlike the original two-gene model in which one X-linked factor is a mutant allele, their X-linked alleles, *YUN* and *SUN*, both are functional allele and actively promote femaleness. This architecture effectively behaves like a “two-gene” model, favouring close linkage between the two Y-linked loci and can explain the inversion observed in the Y-linked region.

Finally, we note that, although a few sex-determination factors have been identified in dioecious plants, and they are very varied, our discovery that, in the SKY regulatory module in Sapindaceae, the *KUN* (*MeGI* ortholog) gene is involved in the actions of *SUN* and *YUN*, demonstrates the involvement of conserved genetic elements in the evolution of dioecy in plants. The broad conservation of *KUN* and *YUN* homologs in angiosperms suggest these factors may control female-male trade-offs and contribute to the evolution of diverse sexual systems in plants.

The evolution of the XY sex determination system in the dioecious *E. cavaleriei* involves loss-of-function mutations involving male-female trade-offs, gene duplication and sRNA mediated gene silencing, as well as large genomic inversions. The closest dioecious species with a well-studied sex-determining system is papaya (*Carica papaya*) in the Caricaceae family within the Brassicales, which diverged from their common ancestor with Sapindales around 100 MYA^14^. Therefore, *E. cavaleriei* represents a valuable system for studying the evolution of sexual system in plants. Furthermore, many Sapindaceae species, including *E. cavaleriei*, are perennial trees, a type of plant once considered intractable for studies of genetic and developmental questions, despite their ecological and evolutionary importance. Our study demonstrates that, with advances in sequencing technologies and comparative and population genomics, biological questions related to perennial trees, which were once considered intractable, can now be studied, ushering in new opportunities for plant biology research.

## Acknowledgments

We are grateful to Drs. Chengcai Chu, Yaowu Yuan, Abdelhafid Bendahmane, and Blake Meyers for critical reading or discussion of the manuscript. We thank Dr. Ming Kang, Dr. Baohuan Wu, Mr. Bin Zhou, Mr. Conglin Xie and Dr. Mingxuan Zheng for their help in locating the E. cavaleriei trees, Dr. Linchuan Liu and Mr. Wenjie Liu for their help in protein co-immunoprecipitation assays, and Dr. Fanjiang Kong and Dr. Huan Liu for their assistance in in-situ hybridization. We also thank members of the XIALAB for kind help, comments and discussion on the projects.

## Funding

National Natural Science Foundation of China #32372665

National Natural Science Foundation of China #32402528

National Natural Science Foundation of China #32072547

Key Area Research and Development Program of Guangdong Province 2022B0202070003

## Author contributions

R.X., Y.X.M., J.K.Z., and C.J.C. designed the experiments; Y.X.M., C.J.C., and Q.P.W. analyzed the data. J.K.Z., W.J.S., Z.Q., W.C.W., X.X.L., and Y.W.H. conducted the experiments. C.Q.S., C.J.C., J.K.Z., and J.X. observed and conducted the study on litchi flowers. Y.X.M., F.Q.W., W.J.S., Y.L.L., and Z.H.Z. collected SanHuaMu samples from the wild. R.X., Y.X.M., and J.K.Z. drafted the manuscript. D.C. made essential suggestions and critically revised the manuscript. All authors participated in data interpretation, edited the manuscript, and approved the final manuscript.

