## Supplementary material for "A trade-off mechanism underpins the evolution of a young two-gene sex-determining system in plants": Supplemetal materials (M&M and Figures)

**The PDF file includes:**

Materials and Methods
Figs. S1 to S37
Tables S1 to S16
References 1 to 55

### Materials and Methods

#### Plant Materials and Growth Conditions

Flower samples of litchi were collected from litchi trees, cultivar “Feizixiao”, grown in an orchard located at South China Agricultural University (Guangzhou, China; N: 23.16195290°, E: 113.36100955°). Flower buds and opened flowers were pooled from three trees to constitute one biological replicate, yielding a total of three biological replicates. The flowers were dissected into various tissues, including carpel and anther, prior to freezing. Following harvest, samples were rapidly frozen in liquid nitrogen and stored at -80°C for RNA sequencing and RT-PCR analysis.

Flower and leaves samples of *E. cavaleriei* were collected from trees grown in a hilly area at Wengyuan, China (N: 24.360879°, E: 114.191531°). A total of 147 tender leaf samples of *E. cavaleriei* were collected from the field, including 69 samples from female individuals and 78 from male individuals. All tender leaves were subjected to DNA extraction using the CTAB method, library construction, and sequencing by Illumina HiSeq2000 and PacBio HiFi (CCS) platforms and high-throughput chromosome conformation capture (Hi-C) technique (Table S1). For the flower samples, flower buds and opened flowers from one tree were pooled to create one biological replicate, resulting in a total of three biological replicates. The female flowers were dissected into various tissues, including carpels and other tissues, while the male flowers were dissected into tissues such as anthers and others, prior to freezing. After harvest, the samples were rapidly frozen in liquid nitrogen and stored at -80°C for RNA sequencing.

*Arabidopsis thaliana* (Columbia-0 (Col-0) ecotype) plants were grown in growth chambers (day/night: 16 h/8 h, 21°C) and transformed using the floral dipping method as described[1]. Tobacco plants used for stable transformation (*Nicotiana tabacum* L. cv. Petit Havana SR1) and transient expression assays (*Nicotiana benthamiana*) were grown under a 16-h light/8-h dark photoperiod at 25°C. Genetic transformation was conducted using the Agrobacterium-mediated transformation method, following the leaf disk procedure described by Horsch[2]. The T-DNA insertion line SALK_206790 (*stk*) were obtained from Arashare (https://www.arashare.cn), while the CS3844 line (*shp1 shp2*) was kindly provided by Professor Ruibo Hu (University of Chinese Academy of Science, China). The CS3844 line (*shp1 shp2*) and SALK_206790 (*stk*) were used as the maternal and paternal lines, respectively. Self-crossing following the initial cross generated the following genotypes: *SHP1 SHP2* $STK^{+/+}$ (*XXYYZZ*), *SHP1 SHP2* $STK^{+/-}$ (*XXYYZz*), *SHP1 SHP2* $STK^{-/-}$ (*XXYYzz*), *shp1 shp2* $STK^{+/+}$ (*xxyyZZ*), *shp1 shp2* $STK^{+/-}$ (*xxyyZz*), *shp1 shp2* $STK^{-/-}$ (*xxyyzz*). Columbia-0 was used as the wild type. Transgenic *Arabidopsis* and tobacco plants were selected on half-strength Murashige and Skoog medium supplemented with either 50 µg/mL Kanamycin or 20 µg/mL Basta salt.

**Genome Survey and Assembly**

Quality assessment of raw sequencing reads and removal of adapters and low-quality sequences were performed by fastp (0.19.5)[3] with default settings. We applied the processed data to estimate genome size, heterozygosity, and repeat content using GCE (1.0.0)[4]. The k-mer_freq_hash pipeline generated 17 bp k-mers and calculates their frequency distribution to create a hash list. GCE adjusted and estimated the genomic information using the k-mer frequency hash list. The male and female genomes of *E. cavaleriei* were estimated to be 292.96 and 296.54 Mb in size, respectively (Table S2), with low heterozygosity of 0.70% and 0.71%, respectively.

The Pacbio HiFi long reads were de novo assembled into contigs using Hifiasm[5] (0.20.0-r639). The completeness of the draft assemble was estimated based on contig length and conserved genes. N50, N90 and the average length of the draft assembly were calculated using a self-designed Python pipeline. Additionally, we assessed genome completeness by evaluating the presence of conserved embryophyte genes using BUSCO with the embryophyte_odb10 dataset[6] (v5.8.0). Hi-C raw data were processed for quality control, mapping, and construction of contact maps using HiC-Pro[7] (v3.1.0). The contact matrix (100 kb) was then used to anchor the genome into 13 pseudochromosomes using EndHiC[8] (1.0.0). To obtain an accurate reference genome (female), we performed manual correction using Juicebox[9] (v2.20.00). The draft assembly of the male genome was anchored into chromosome level using the female reference genome, with the scaffold program Ragtag[10] (v2.1.0). The Hi-C contact matrices of both female and male genome were calculated, corrected, and plotted using HiCExplorer[11] (3.7.2).

**Genome Annotation**

Repeat sequences were masked and annotated by EDTA[12] (v2.1.0) with the parameter "--species others, anno 1". RNA-seq from floral tissues were mapped to the repeat-masked genomes using STAR[13] (v2.5.3a) with parameters "-alignIntronMax 20000 --alignMatesGapMax 25000 --outFilterMultimapNmax 50". The protein sets of *Arabidopsis* (TAIR10), rice (IRGSP), litchi and yellowhorn were used as the homologous protein set. Transcriptome and homologous protein evidence were integrated to annotate the gene structure of the genomes using BRAKER3 (v3.0.8). The coding sequences and protein sequences were extracted using GffRead[14] (0.12.7). The functions of the entire protein set of *E. cavaleriei* were annotated using the eggNOG-mapper[15] (v2) database. Additionally, the proteins were assessed for completeness using BUSCO (v5.8.0; embryophyte_odb10) in protein mode.

**Transcriptomic Analysis**

The raw FASTQ data of RNAseq were pre-preprocessed by fastp[3] (0.19.5) with default parameters. Then, STAR[13] (2.5.3a) was used to build the genome index and align the FASTQ files to reference genomes. The estimated expression levels (Transcript per Kilobase per Million mapped reads; TPM) of the samples were calculated by StringTie[16] (v2.2.1). Weighted gene correlation network analysis (WGCNA)[17] was performed on the expression matrix of floral tissues from both litchi and *E. cavaleriei*. The expression of candidate genes were visualized as heatmaps using TBtools-II[18].

**GO Enrichment**

The Gene Ontology (GO) annotations of total coding-sequence set was conducted using eggNOG-mapper (v2.1.12)[15]. Then, genes in candidate modules from WGCNA were used as the target list, and GO enrichment analysis was performed using GO Enrichment function of TBtools-II[18] to obtain multiple-testing corrected p-values (FDR/q-value) and target gene IDs for each term. Visualization of GO terms is implemented using the R package ggplot2[19].

**Small RNA Analysis**

The adaptor and ribosome RNA sequences in raw small RNA (sRNA) data were identified and removed using sRNAminer[20] (v1.1.2). And genome index was built by bowtie-build[21] (2.2.5). Clean sRNA sequence reads were mapped into corresponding genomes using bowtie with the parameter '-a --best --strata -m 20 -v 1 -p 1'. The sRNA distribution of the $SUN^{MAO}$ locus was visualized using IGV-sRNA[22] (0.6.73).

**Single Nucleus RNA-seq Analysis**

Nuclei isolation from the ovaries of *E. cavaleriei* at stage 5 and *L. chinensis* at stage 8 was followed by 10x Genomics snRNA-seq library preparation and sequencing. Raw reads were mapped to the genome of *E. cavaleriei* (female genome) and *L. chinensis* by CellRanger (8.0.1) using the default parameters. For quality control purpose, genes expressed in less than five nuclei were discarded, and cells with gene counts fewer than 100 were removed by the CreateSeuratObject function in Seurat (5.1.0)[23]. The final matrix was subsequently analyzed by

Seurat (5.1.0)[23] with recommended parameters for normalization (normalization.method = "LogNormalize", scale.factor = 10000), scaling, and finding variable features. Subsequently, we performed principal component analysis (PCA) on the top 2,000 variable features (RunPCA function) and constructed a nearest-neighbor graph using the first 30 principal components (FindNeighbors function), followed by cell population clustering through modularity optimization (FindClusters function). Nonlinear dimensionality reduction was performed on the single-cell matrix using Uniform Manifold Approximation and Projection (UMAP) based on the top 30 principal components (RunUMAP function), achieving segregation of cell populations through two-dimensional visualization. Gene expression patterns of target genes (*YUN*, *SUN* and *KUN*) were visualized using FeaturePlot and VlnPlot.

**Prediction of Binding Site of YUN and KUN**

The position weight matrices (PWMs) of the YUN (MA0585.1 and MA0585.2; MA0005.1 and MA0005.2) and KUN (MA1214.1 and MA1214.2; MA1213.1; MA1215.1 and MA1215.2) homologous genes were retrieved from the JASPAR (2024) database[24]. The upstream 2 kb sequences of the target genes were analyzed for potential binding sites using MEME Suite (v5.5.2) [25] with the MAST tool in default parameters. The predicted binding sites were visualized using TBtools-II[18].

**Conserved Non-Coding Sequence (CNS) Analysis**

Extract the 3-kb upstream regions of target genes from Sapindaceae species and their closely related species (*Citrus sinensis*) using the SeqKit (0.7.2)[26] toolkit. Subsequently, mVISTA[27] was used to predict conserved non-coding sequences (CNS) in promoter regions and visualize genomic similarity.

**Evolutionary Analysis**

Genomes and annotations from Sapindaceae species and other eudicots species were collected from different databases (detail in Table S13), and the longest transcripts were extracted using TBtools-II[18]. Orthogroup, single-copy ortholog sequences and species trees for 14 species were inferred using Orthofinder2[28] (v2.5.4) with parameters "-M msa -S diamond". Single-copy ortholog genes were aligned in phylip format using MUSCLE[29] (v3.8.31). Species divergence times were estimated using the MCMCTree algorithm from PAML (v4.8), incorporating two calibration points: the divergence between Sapindaceae species and *Vitis*, which occurred approximately 107 to 120 million years ago (MYA), and the divergence between Sapindaceae species and *Citrus*, which occurred around 65 MYA. The divergence time tree was visualized using FigTree (v 1.4.3).

**Synteny Analysis**

Homologous gene pairs were identified by JCVI[30] (v1.2.7), while $K_a$ and $K_s$ values were calculated using Simple Ka/Ks Calculator in TBtools-II. The collinear relationships between species were visualized by the graphics module in JCVI[30] (v1.2.7).

**Haplotypic genome alignment analysis**

Pairwise alignments between haplotypic genomes were performed using minimap2 (v2.24-r1122)[31] with a sequence divergence setting of ~5% (-ax asm5). Subsequently, SyRI (v1.6.3)[32] was used to detect structural variants and genomic rearrangements from the alignment results. Finally, these variations between haplotypic genomes were visualized using plotsr (v0.5.4)[33].

**Identification of Centromere Regions**

The CentromereLocation plugin in TBtools-II[18] was used to predict centromeres in the genome sequence. This tool relies on Tandem Repeats Finder (TRF)[34] to perform sliding-window statistical analysis (window size: 5 kb; step size: 1 kb) on tandem repeat elements (100–200 bp) and calculate their copy numbers. Regions exhibiting high tandem repeat copy numbers were identified as candidate centromeres. Finally, python scripts were used to visualize the distribution of tandem repeat copy numbers and annotate the predicted centromeric regions.

**Identification of Sex Determination Region**

A total of 147 *E. cavaleriei* individuals were collected from the field, including 78 males and 69 females. DNA extraction, library construction, sequencing, and quality control were performed according to the methods described above. The sequencing data were mapped to the reference genome using bwa-mem algorithm of BWA[35] (Burrow Wheeler Aligner; 0.7.17-r1188), with default parameters. SAM files were converted to BAM files using SAMtools[36] (1.6), and the BAM files were sorted using sambamba[37] (0.8.2) . The coverage of the BAM files for all samples was calculated using the bedcov program of SAMtools[36] (1.6) with a sliding window size of 10 kb.

Total SNP information was extracted from these BAM files using the HaplotypeCaller program of GATK[38] (4.4.0.0). Plink2[39] (2.0) was used to filter SNPs with high missingness and high linkage, with the parameters: --indep-pairwise 50 10 0.1. Fixation index (*F*st) and nucleic acid diversity (Pi) for SNPs in male and female populations were calculated by VCFtools[40] (0.1.16). Genome wide association study (GWAS) analysis was conducted using the linear mixed model (LMM) by the GEMMA[41] (v0.98.5). The R package *qqman*[42] (0.1.9) was used to visualize the *p* values representing the statistical significance of the associations. Finally, PopLDdecay[43] (3.43) was used to calculate and visualize the linkage disequilibrium (LD) within the sex determination region (SDR), further refining the localization of the SDR.

**Male Specific K-mers Analysis**

All 147 Illumina sequencing data were processed using KMC (v3.2.2)[44] to generate 27-bp k-mer sets. Low-frequency k-mers (occurring fewer than 5 times; parameter: -ci5) were removed to reduce noise. The male k-mer sets were merged using kmc_tools complex to create a unified male k-mer union: male_union_set=male_set1+ male_set2+…+ male_set78. To isolate male-specific k-mers, the intersection with all female k-mer sets was sequentially removed: male_specific_kmer_set = male_union_set - female_set1 - female_set2 - ... - female_set69. The male_specific_kmer_set were converted into nucleotide sequences using kmc_dump. These k-mer sequences were strictly aligned to the male and female reference genomes of *E. cavaleriei* using Bowtie (v10.4.0)[45] with the following parameters: -v 0 --best --strata -m 1. The k-mer alignment depth was calculated in 10,000 bp sliding windows using BEDTools (v2.30.0)[46] coverage. The distribution of depth across genomic regions was plotted using matplotlib.pyplot (Python).

**Correlation Analysis of Genes Expression with *SUN***

Genes with an average expression level >1 in ovary development-related modules (brown module for *E. cavaleriei*; green and black modules for *L. chinensis*) were selected as candidate gene sets. Pearson correlation coefficients (*R*) were calculated between *SUN* and candidate gene expression. And the correlations between *SUN* and candidate genes were visualized as scatter plots using matplotlib.pyplot (Python).

**Oligonucleotides**

The sequences of all oligonucleotides used for plasmid construction, genotyping PCRs, qRT-PCR, and *in situ* probe labelling are listed in Table S15.

***In situ* Hybridization**

*In situ* hybridization in *E. cavaleriei* and litchi was performed as described previously (*93*), with minor modifications. Female flower carpels at stage 5 of *E. cavaleriei* and stage 8 of litchi were excised and fixed in FAA solution [50% (v/v) ethanol, 5% (v/v) acetic acid, and 3.7% (v/v) formalin]. The samples were then embedded in paraffin, sectioned into 8 µm thick slices using a microtome (HistoCore MULTICUT, Leica), and hybridized with specific probes. Digoxigenin-labeled sense and antisense RNA probes were generated via PCR amplification, followed by transcription using SP6 and T7 RNA polymerase (11175025910, Roche, Mannheim, Germany). The primer sequences used are listed in Table S15.

**Plasmid Construction**

All constructs were generated using In-Fusion cloning (ClonExpress II One Step Cloning kit, Vazyme) or Gateway system.

Both In-Fusion cloning and Gateway system were used to construct expression vectors for *Arabidopsis* transformation, aiming to functionally complement the *stk* mutant by overexpressing *LcYUN* and *EcYUN*, each driven by the *AtSTK* promoter (*pAtSTK::LcYUN* and *pAtSTK::EcYUN*). The plasmid constructs were generated as previously described [47]. Firstly, to generate entry vectors, DNA fragments containing the 3,534 bp promoter region of *AtSTK* and the coding sequences (CDS) of *LcYUN* (675 bp) and *EcYUN* (675 bp), excluding the stop codon, were amplified from wild-type *Arabidopsis* genomic DNA and cDNA of litchi and *E. cavaleriei* female flowers, respectively, using the following primer pairs: TOPO-*pAtSTK*-F/R for *AtSTK* promoter, TOPO-*LcYUN*-F/R for *LcYUN* CDS, and TOPO-*EcYUN*-F/R for *EcYUN* CDS. The In-Fusion cloning system (ClonExpress II One Step Cloning Kit) was used to fuse both *AtSTK* promoter and *LcYUN* or *EcYUN* CDS into the pENTR/D/TOPO vector (Invitrogen). Subsequently, destination vectors for *pAtSTK::LcYUN* and *pAtSTK::EcYUN* were generated by LR reactions using LR Clonase II (Invitrogen) as described by Zhou *et al* [47].

To verify the dosage sensitivity of *YUN* (*AGL11*) orthologs, transgenic plants were generated in both *Arabidopsis* and tobacco (*N. tabacum*). For *Arabidopsis*, a 7,921 bp DNA fragment, comprising the 3,534 bp *AtSTK* promoter region and the 4,387 bp complete *STK* genomic region, was amplified from wild-type genomic DNA and cloned into the pCAMBIA1300-eGK vector, generously provided by Professor Fei Zhang from Huazhong Agricultural University [48]. The amplification was performed using the primers 1300eGK-*AtSTK*-F1/R1 and 1300eGK-*AtSTK*-F2/R2 for segmented amplification. The amplified fragments were then assembled into the pCAMBIA1300-eGK vector using the In-Fusion system. For tobacco, the large size of the *NtYUN* genomic sequence (over 12 kb) pose challenges for cloning and could potentially affect expression efficiency. To address this, the CDS sequence of *NtYUN* was selected for vector construction. To generate the entry vector, A 3,534 bp *NtYUN* promoter region and the 696 bp *NtYUN* CDS (excluding the stop codon) were amplified from wild-type tobacco genomic DNA and ovary cDNA, respectively, using the primer pairs TOPO-*pNtYUN*-F/R for the promoter and TOPO-*NtYUN*-F/R for the CDS sequence. The amplified fragments were cloned into an entry vector via the In-Fusion system. Subsequently, LR reactions were performed to generate the destination vector, following the procedure previously described.

To investigate the function of *SUN*, transgenic plants overexpressing *SUN* were generated in both *Arabidopsis* and tobacco (*N. tabacum*). In *Arabidopsis*, the CDS without the stop codon of *EcSUN* (2697 bp) and *LcSUN* (3099 bp), were amplified using the primer pairs pC2300-*EcSUN*-F/R and pC2300-*LcSUN*-F/R, respectively. The amplified fragments were then inserted into the SalI/SpeI-digested pC2300-35S-eGFP vector using the ClonExpress II One Step Cloning Kit. For tobacco, the CDS of *EcSUN* and *LcSUN* were amplified with the primer pairs pE201-*EcSUN*-F/R and pE201-*LcSUN*-F/R, respectively, and subsequently inserted into the XhoI/SpeI -digested pEarly-Gate-201 vetor using the ClonExpress II One Step Cloning Kit.

To verify the silencing effect of $SUN^{MAO}$ on *SUN*, tobacco (*N. benthamiana*) was used for transient transfection. The full length of $SUN^{MAO}$ (1509 bp) was divided into two segments, A1

(832 bp) and S2 (677 bp), which were synthesized by Tsingke Biotechnology Co., Ltd. (China). The A1 and S2 segments were amplified using the primer pairs A1-F/R and S2-F/R, respectively, and subsequently spliced together before being inserted into the XhoI/SpeI -digested pEarly-Gate-201 vetor using the ClonExpress II One Step Cloning Kit. The pC2300-*EcSUN*-eGFP and pC2300-*LcSUN*-eGFP vectors, previously used for functional identification, were also employed in this transient transfection experiment.

For fluorescence-based protein stability assays, we generated expression constructs using the Gateway cloning system. The CDS without the stop codon of *EcSUN* (2697 bp), *LcSUN* (3099 bp), *EcKUN* (876 bp), and *LcKUN* (849 bp, with the stop codon) were PCR-amplified with primer pairs *EcSUN*-TOPO-F/R, *LcSUN*-TOPO-F/R, *EcKUN*-TOPO-F/R, and *LcKUN*-TOPO-F/R, respectively, and cloned into the pENTR/D-TOPO entry vector (Invitrogen). Through LR recombination using LR Clonase II (Invitrogen), we created the following destination vectors: (1) proUBQ10:*EcKUN*-GFP and proUBQ10:*GFP-LcKUN* for GFP-tagged constructs, and (2) 35S:*EcSUN*-RFP and 35S:*LcSUN*-RFP for RFP-tagged constructs[49,50].

To construct plasmids for yeast two-hybrid (Y2H) assays, we prepared both bait and prey constructs using In-Fusion cloning. For bait constructs, the coding sequences (CDS) of *EcSUN* (2697 bp) and *LcSUN* (3099 bp) were PCR-amplified using primer pairs BD-*EcSUN*-F/R and BD-*LcSUN*-F/R, respectively. The products were digested with NdeI/NotI and cloned into the pGBKT7 vector. For prey constructs, the CDS of *EcKUN* (879 bp) and *LcKUN* (849 bp) were amplified with AD-*EcKUN*-F/R and AD-*LcKUN*-F/R primers, digested with EcoRI/BamHI, and ligated into pGADT7.

For co-immunoprecipitation (Co-IP) assays, we generated FLAG-tagged (EcSUN/LcSUN) and GFP-tagged (EcKUN/LcKUN) constructs via In-Fusion cloning. The coding sequences (excluding stop codons) of *EcSUN* (2,697 bp) and *LcSUN* (3,099 bp) were PCR-amplified using gene-specific primers (*EcSUN*-FLAG-F/R and *LcSUN*-FLAG-F/R) and recombined into KpnI/BamHI-linearized pCAMBIA1300-C-FLAG vector, while *EcKUN* (876 bp) and *LcKUN* (846 bp) were amplified with *EcSUN*-eGFP-F/R and *LcSUN*-eGFP-F/R primers and cloned into the corresponding KpnI/BamHI-linearized pCAMBIA1300-C-eGFP vector[51].

For luciferase complementation imaging (LCI) assays, we generated fusion constructs by cloning full-length coding sequences into split-luciferase vectors. The CDS without the stop codon of *EcSUN* (2697 bp) and *LcSUN* (3099 bp) were PCR-amplified using primer pairs *EcSUN*- nLUC-F/R and *LcSUN*- nLUC-F/R, respectively, while *EcKUN* (879 bp) and *LcKUN* (849 bp) CDS were amplified with *EcKUN*-cLUC-F/R and *LcKUN*-cLUC-F/R. Amplified fragments were digested with KpnI/SalI (for nLUC fusions) or KpnI/BamHI (for cLUC fusions), then ligated into the corresponding pCAMBIA1300-nLUC or pCAMBIA1300-cLUC vectors.

To generate protein expression constructs for DNA affinity purification sequencing (DAP-seq) and electrophoretic mobility shift assays (EMSA), we amplified the full-length *LcYUN* CDS (678 bp) from litchi female flower cDNA for DAP-seq analysis, which was subsequently cloned into BamHI-linearized pIX-HALO vector using the ClonExpress II One Step Cloning Kit (Vazyme). For EMSA studies, we amplified the DNA-binding domains of *EcKUN* (354 bp), *EcYUN* (243 bp), *LcKUN* (354 bp), and *LcYUN* (243 bp) using gene-specific primer pairs (*EcKUN*-GST-F/R, *EcYUN*-GST-F/R, *LcKUN*-GST-F/R, and *LcYUN*-GST-F/R, respectively) and cloned these fragments into EcoRI-digested pGEX4T-GST vector using the same cloning system. All recombinant constructs were verified by Sanger sequencing prior to protein expression.

To verify of transcriptional regulation in the SUN-KUN-YUN module, yeast one-hybrid (Y1H) assays were conducted. The CDS of *EcKUN* (879 bp), *EcYUN* (678 bp), *LcKUN* (849 bp), and *LcYUN* (678 bp) were amplified using primer pairs AD-*EcKUN*-F/R, AD-*EcYUN*-F/R, AD-*LcKUN*-F/R, and AD-*LcYUN*-F/R, respectively, and cloned into EcoRI/BamHI-digested pGADT7 via the ClonExpress II One Step Cloning Kit (Vazyme). Promoter fragments containing cognate binding sites—*EcSUN* (279 bp), *EcYUN* (339 bp), *LcSUN* (279 bp), and *LcYUN* (507 bp) for KUN proteins, and *EcKUN* (211 bp) and *LcKUN* (134 bp) for YUN proteins—were amplified with pAbAi-*pEcSUN*-F/R, pAbAi-p*EcYUN*-F/R, pAbAi-*pLcSUN*-F/R, pAbAi-*pLcYUN*-F/R, pAbAi-*pEcKUN*-F/R, and pAbAi-*pLcKUN*-F/R, then inserted into KpnI/XhoI- or SacI/XhoI-digested pAbAi vectors. Constructs were validated by sequencing and transformed into yeast strain Y1HGold for interaction assays on SD/-Leu + Aureobasidin A (AbA) plates.

For reporter constructs of the dual-luciferase reporter assays, promoter fragments of *EcSUN* (1877 bp), *EcKUN* (2070 bp), *EcYUN* (1694 bp), *LcSUN* (2000 bp), *LcYUN* (1936 bp), and *LcKUN* (2010 bp) were cloned into KpnI/BamHI-digested pGreenII 0800-LUC vector using the ClonExpress II One Step Cloning Kit (Vazyme) with gene-specific primer pairs (0800-p[Gene]-F/R). For effector constructs, coding sequences of *EcKUN* (879 bp), *EcYUN* (678 bp), *LcYUN* (678 bp), and *LcKUN* (849 bp) were inserted into EcoRI/BamHI-digested pGreenII 62-SK vector using the same cloning kit with corresponding 62SK-[Gene]-F/R primers.

**DNA and RNA Preparation**

Genomic DNA was extracted from leaves using the 2 x CTAB extraction buffer (Coolaber, China). Briefly, samples were homogenized in liquid nitrogen, lysed in 800 μL preheated CTAB buffer (65°C, 30 min), and extracted with chloroform:isoamyl alcohol (24:1). DNA was precipitated with 2/3 volume isopropanol, washed twice with 70% ethanol, and air-dried. The pellet was resuspended in 50 μL sterile $ddH_2O$ and stored at –20°C. DNA quality was assessed spectrophotometrically (1 μL aliquot).

High-quality total RNA was isolated from various tissues using RNAiso Plus reagent (Takara) following the manufacturer's instructions. In litchi, whole flower buds at stages 1 and 2, as well as flowers at stages 3 to 8 (female flowers up to stage 12), which were dissected into carpel and anther tissues, were collected for RNA extraction. Similarly, in *E. cavaleriei*, whole flower buds at stage 1 and 2, as well as flowers at stages 3 to 6, dissected into carpel and other tissues for female flowers or anther and other tissues for male flowers, were collected for RNA extraction. Additionally, developing *Arabidopsis* leaves/siliques and tobacco leaves/ovaries were collected for RNA extraction. For the litchi samples described above, total RNA was extracted using RNAiso Plus reagent combined with Fruit-mate (Takara) to remove polysaccharides and polyphenols. The extracted RNA was stored at -80°C for sequencing and qRT-PCR analysis. For small RNA (sRNA) sequencing in *E. cavaleriei*, libraries were constructed from 10 μg of pooled total RNA (RIN ≥ 7.5; Agilent Bioanalyzer 2100) using the NEBNext® Multiplex Small RNA Library Prep Kit (New England Biolabs) following the manufacturer's protocol.

**Genotyping PCRs and qRT-PCR**

Mutants and transgenic *Arabidopsis* and tobacco plants were screened by amplifying specific DNA fragments, with the primer pairs listed in Table S15. For *E. cavaleriei* sex identification, male plants yielded a 722-bp product (F1/R1 primers) and a 453-bp product (F2'/R1' primers), while female plants produced a 961-bp product (F1/R2 primers) and an 807-bp product (F1'/R1' primers). To identify the *stk* allele in *Arabidopsis*, amplification with primers LBb1.3 (T-DNA-specific) and *AtSTK*-RP yields a ~500-bp product in plants containing the *stk* allele, while amplification with *AtSTK*-LP/*AtSTK*-RP yields a 975-bp product in plants with a wild-type *STK* allele. *SHP1* and *SHP2* genotyping followed the method by Pinyopich *et al*[52]. For *SHP1*, amplification with primers *shp1*-gtyF and T-DNA-LB yields a 597-bp product for plants containing the *shp1* allele, while amplification with *SHP1*-gtyF/*SHP1*-gtyR yields an 871-bp product in wild-type *SHP1* plants, with no product in plants homozygous for the *shp1* allele. For *SHP2*, amplification with primers *SHP2*-gtyF/*SHP2*-gtyR yields a 1.5-kb product in wild-type plants and a 2.8-kb product in plants homozygous for the *shp2* allele. To identify *pAtSTK::LcYUN* and *pAtSTK::EcYUN* transgenic *stk* mutants, amplification with primers *LcYUN*-F/R or *EcYUN*-F/R produces a 678-bp product in plants containing *pAtSTK::LcYUN* or *pAtSTK::EcYUN*, respectively. For identifying *pAtSTK::AtSTK* transgenic *Arabidopsis*, amplification with primer *AtSTK*gty-F/R yields a 1502-bp product. Similarly, for identifying *pNtYUN::NtYUN* transgenic tobacco, amplification with primer *NtYUN*gty-F/R produces an 887-bp product.

To evaluate target gene expression in plants, reverse transcription was performed with 1 μg of total RNA using TransScript® One-Step gDNA Removal and cDNA Synthesis SuperMix (AT311-02, TransGen Biotech, China). RT-qPCR was conducted in 10-μL reactions containing 1 μL of 1:5 diluted cDNA, 0.5 μL of specific primers, 3 μL distilled deionized water, and 5 μL GoTaq® qPCR Master Mix (Promega) in a BioRad CFX96 Real-Time PCR Detection System. The elongation factor *EF1α* from litchi, *E. cavaleriei*, *Arabidopsis,* and tobacco served as the reference gene for their respective samples. The relative transcript abundance was calculated as the ratio of target gene expression to *EF1α* expression using the $2^{-\Delta\Delta CT}$ method. The primers for target gene detection are listed in Supplemental Table S15, and the melt curve and amplification curve data (provided in Table S15) demonstrate the primer efficiency in qRT-PCR.

**Phenotypic Analysis**

Pollen development analysis, including Alexander staining and SEM, was conducted as described by Zhou et al[47]. To assess pollen viability in *Arabidopsis* and tobacco, mature anthers were soaked overnight in Alexander stain and observed under an CX31 biological microscope

(OLYMPUS). For SEM analysis, mature pollen was coated with gold-palladium and observed using an EVO-MA-15 scanning electron microscope (ZEISS).

**Verification of *SUN$^{MAO}$*-Mediated *SUN* Gene Silencing**

The pE201-SUN$^{MAO}$, pC2300-*EcSUN*-eGFP and pC2300-*LcSUN*-eGFP constructs were individually transferred into *Agrobacterium tumefaciens* strain GV3101 and infiltrated into *N. benthamiana* leaves for transient expression assays. The co-injection combinations include: pC2300-*EcSUN*-eGFP, pC2300-*EcSUN*-eGFP + pE201-*SUN$^{MAO}$*, pC2300-*LcSUN*-eGFP and pC2300-*LcSUN*-eGFP + pE201-*SUN$^{MAO}$*. For co-injection, *Agrobacterium* carrying the target plasmids was resuspended in infiltration buffer (10 mM MES, 10 mM $MgCl_2$, 150 μM acetosyringone, pH = 5.6) at $OD_{600}$ = 0.8 and incubated at room temperature for 3 hours. Infiltrated leaf samples were harvested 3 days post-injection for fluorescence observation and expression level detection.

**Protein Interaction Assays**

Co-IP assays were performed using transient co-expression in *N. benthamiana* leaves by infiltrating *Agrobacterium tumefaciens* GV3101 strains carrying the following construct combinations: FLAG-*EcSUN* + *EcKUN*-GFP and FLAG-*LcSUN* + GFP-*LcKUN*, along with appropriate negative controls. The infiltrated leaves were flash-frozen in liquid nitrogen and ground into a fine powder. Total proteins were extracted in ice-cold buffer [50 mM Tris-HCl (pH 8.0), 150 mM NaCl, 5 mM EDTA, 0.1% (v/v) Triton X-100, 0.2% (v/v) Nonidet P-40 (Roche), and protease inhibitor cocktail (2 tablets/100 mL, Roche)]. Lysates were clarified by centrifugation (14,000 × g, 15 min, 4°C), and supernatants were incubated with anti-EBS7 antibody (4°C, 4 h), followed by protein A/G-agarose beads (Abmart). Beads were washed four times with extraction buffer (600 × g, 2 min, 4°C), resuspended in 2× SDS loading buffer, and denatured (95°C, 10 min).

Immunoprecipitated proteins were separated by SDS-PAGE and analyzed by immunoblotting with anti-FLAG (Sigma-Aldrich, #A8592) and anti-GFP (Abcam, #ab290) antibodies.

Y2H assays were performed by co-transforming bait (pGBKT7-*EcSUN* or pGBKT7-*LcSUN*) and prey (pGADT7-*EcKUN* or pGADT7-*LcKUN*) constructs into yeast strain Y2HGold (Clontech) using the lithium acetate/PEG method. Transformants were selected on synthetic defined minimal medium lacking tryptophan and leucine (SD/-Trp/-Leu) at 30°C for 2-3 days. Protein-protein interactions were evaluated by colony growth on stringent selection medium (SD/-Trp/-Leu/-His/-Ade) supplemented with 300 ng/mL aureobasidin A (AbA) following 3 days of incubation at 30°C.

For luciferase complementation imaging (LCI) assays, we tested protein-protein interactions using nLUC-*EcSUN* + cLUC-*EcKUN* and nLUC-*LcSUN* + cLUC-*LcKUN* combinations with corresponding negative controls. *Agrobacterium tumefaciens* GV3101 strains harboring these constructs were cultured overnight at 28°C, pelleted by centrifugation (5,000 rpm, 8 min), and resuspended in infiltration buffer [50 mM $MgCl_2$, 50 mM MES (pH 5.7), 200 μM acetosyringone] to $OD_{600}$ = 1.5. After 2-3 h dark incubation at room temperature, bacterial suspensions were co-infiltrated (1:1 ratio of nLUC- and cLUC-fusion constructs) into *N. benthamiana* leaves using needleless 1-mL syringes. Infiltrated plants were maintained under standard growth conditions for 24 h followed by 24 h dark adaptation before luminescence detection using a Night SHADE LB 985 imaging system (Berthold Technologies).

**DAP-seq and data analysis**

The DNA affinity purification sequencing (DAP-seq) experiment in litchi was performed as described by Zhang *et al*[53]. Briefly, total genomic DNA was extracted from young litchi leaves to construct a DNA library, and the *LcYUN* coding sequence was cloned into pIX-HALO vector and translated in vitro. After incubating the litchi genome DNA library with LcYUN, the DNA-protein complex was eluted, amplified using indexed primers, and subsequently sequenced.

The raw sequencing data were pre-processed by fastp[3] (0.19.5) with default parameters. And these filtered data were mapped to litchi reference genome by bowtie[21] (2.2.5). And peak calling was performed by MACS[54](v3.0.2).

The binding sites of *AGL1* (MA0585), *AGL5* (MA0548), and *AG* (MA0005),which is a homologous gene of *YUN* (*STK*), were download from JASPAR[24] database. MAST program of the MEME Suite[55] (5.5.7) was used to identify homologous binding sites in the region with the enrichment peak of DAP-seq of *LcYUN*.

**Electrophoretic mobility shift assay**

The DNA-binding domains of *EcKUN*, *EcYUN*, *LcKUN*, and *LcYUN* were cloned into the pGEX4T-GST vector to generate GST-fusion proteins, which were expressed in *E. coli Rosetta* (DE3) cells through induction with 0.2 mM IPTG at 37°C for 5 hours. For EMSA analysis, biotin-labeled probes were prepared using the EMSA Probe Biotin Labeling Kit (Beyotime, China), including: (1) *EcSUN*, *EcYUN*, *LcSUN*, and *LcYUN* promoter fragments containing KUN-binding sites, and (2) *EcKUN* and *LcKUN* promoter fragments containing YUN-binding sites. Binding reactions were performed with the LightShift® Chemiluminescent EMSA Kit (Thermo Fisher Scientific, 20148), including competition assays with unlabeled identical probes (100×/500× molar excess). All probe sequences are provided in Supplemental Table 15.

**Dual-luciferase Reporter Assays**

Dual-luciferase reporter assays were conducted following previously described methods (*99*) to investigate KUN-*SUN*/*YUN* and YUN-*KUN* regulatory interactions. Two experimental systems were employed: (i) for KUN-*SUN*/*YUN* analysis, pGreenII 0800-LUC reporter constructs containing either *EcSUN*/*EcYUN* promoters (paired with EcKUN effector) or *LcSUN*/*LcYUN* promoters (paired with LcKUN effector) in the pGreenII 62-SK vector; and (ii) for YUN-*KUN* regulation, *EcKUN* or *LcKUN* promoter-driven reporters with their cognate YUN effectors. All constructs were introduced into *Agrobacterium tumefaciens* GV3101 (pSoup) and adjusted to $OD_{600}$ = 0.8 in infiltration buffer (10 mM MES, 10 mM MgCl2, 150 μM acetosyringone, pH 5.6). Following 3 h incubation at room temperature, bacterial suspensions were co-infiltrated into *N. benthamiana* leaves. Luciferase activity was quantified 72 h post-infiltration using the Dual-Luciferase® Reporter Assay System (Yeasen), with all primer sequences detailed in Supplementary Table S15.

**Yeast One-Hybrid (Y1H) Assays**

For yeast one-hybrid (Y1H) assays, all recombinant pAbAi reporter constructs were linearized with BstBI restriction enzyme (NEB) and transformed into Y1HGold competent yeast cells (WEIDI) using the lithium acetate (LiAc)/PEG method. Transformants were initially selected on SD/-Ura medium to determine the minimal inhibitory concentration of Aureobasidin A (AbA) for each bait-reporter pair through gradient screening (0-400 ng/mL). For interaction analysis, yeast strains co-transformed with AD-effector (pGADT7 constructs) and pAbAi-reporter constructs were plated on SD/-Leu selection medium supplemented with optimized concentrations of Aureobasidin A (AbA): 125 ng/mL for EcKUN+*EcSUNpro*, EcKUN+*EcYUNpro*, EcYUN+*EcKUNpro*, and LcKUN+*LcYUNpro* interactions; and 150 ng/mL for LcKUN+*LcSUNpro* and LcYUN+*LcKUNpro* interactions. All plates were incubated at 30°C for 3-5 days. Negative controls consisted of yeast transformed with empty pGADT7 vector paired with each reporter construct.

**Data Analysis**

All experiments were carried out in at least three replicates. Data were statistically analyzed using Microsoft Excel 2010 to calculate means and standard errors (SE). Comparisons between groups were performed using Student's *t* tests in Microsoft Excel 2010 or a two-way ANOVA test in SPSS (version 17.0).

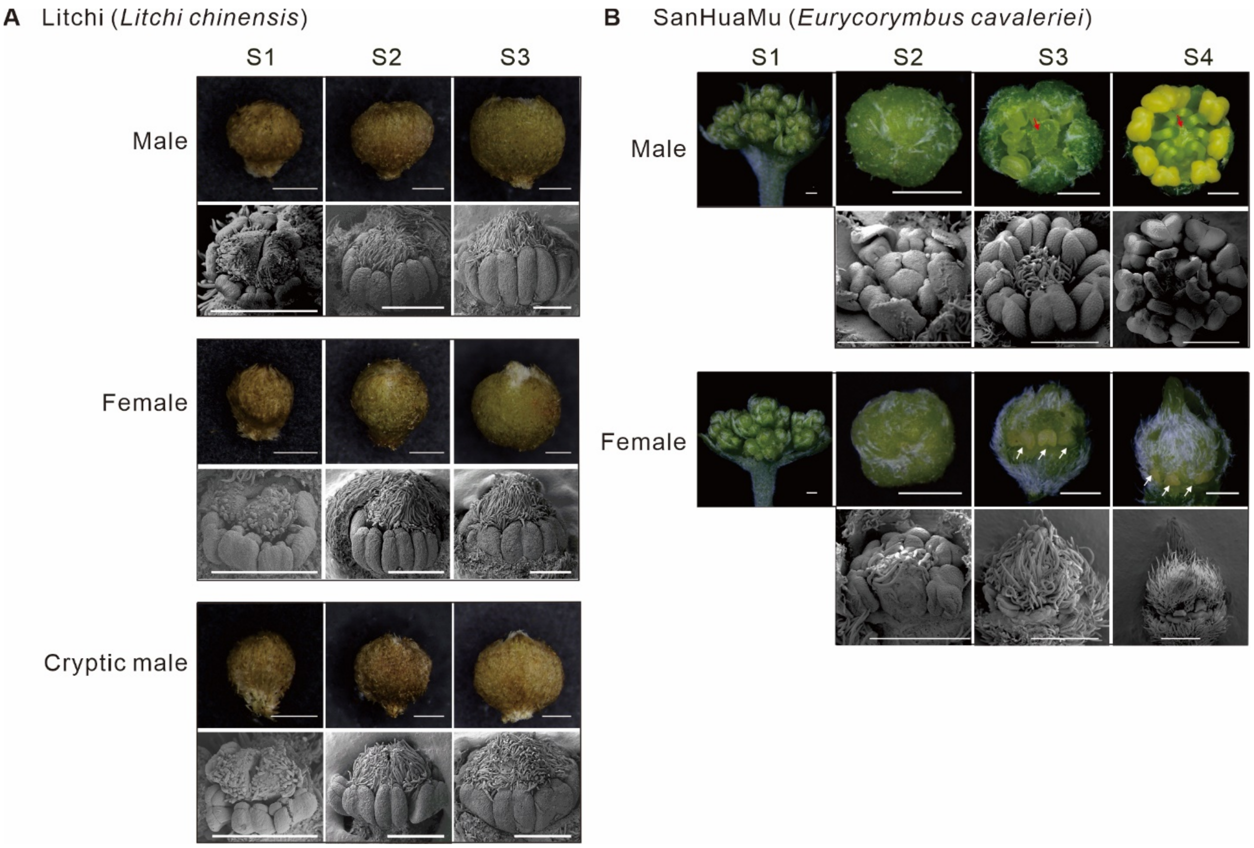


**Fig. S1. SEM (Scanning Electron Microscope) observation of early flower buds in litchi and *E. cavaleriei*.**

**(A)** Early flower buds at stages 1 (0.5-1.0 mm), 2 (1.0-1.5 mm), and 3 (1.5-2.0 mm) of male, female, and cryptic male flowers in litchi. **(B)** Early flower buds at stages 1 (< 0.5 mm), 2 (0.5-1.0 mm), 3 (1.0-1.5 mm) and 4 (1.5-2.0 mm) of male and female flower in *E. cavaleriei*. The red arrows in male flowers indicate abortive carpels, while the white arrows in female flowers denote abortive anthers. The scale bars in the flower photos represent 0.5 mm, while the scale bars in the SEM images represent 0.1 mm.

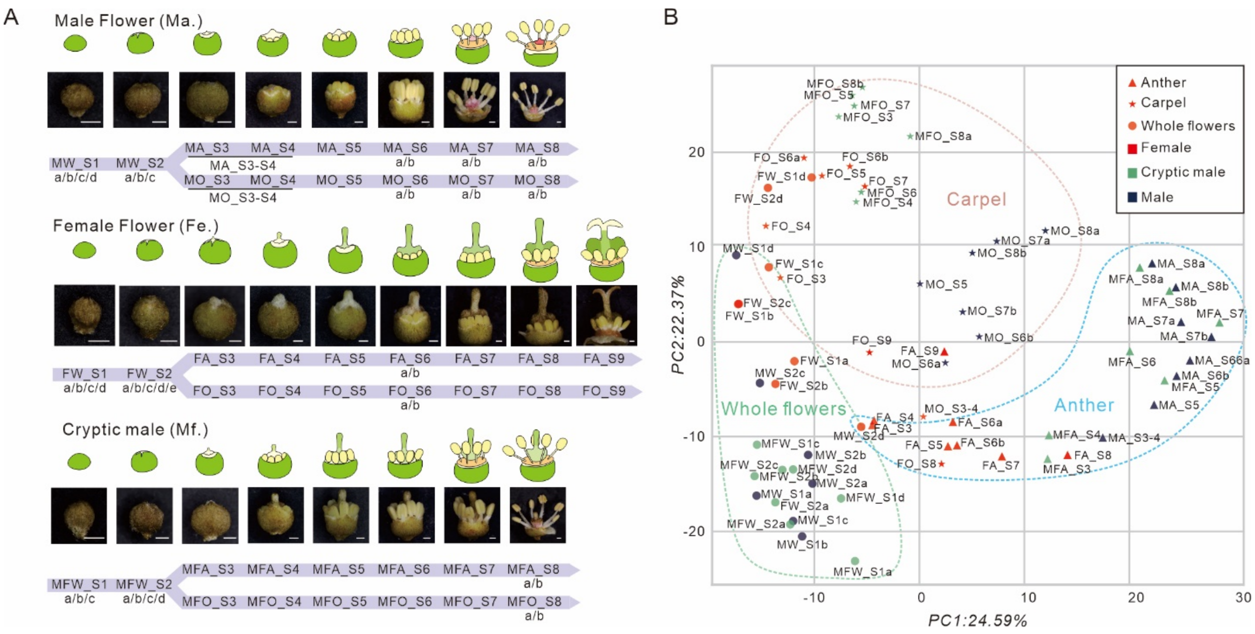


**Fig. S2. Transcriptome analysis of flowers in litchi.**

**(A)** Three types of flowers were collected according to developmental stages. Whole flower buds were retained at stage 1 and stage 2, while flowers at stage 3 to 9 were separated into carpel and anther tissues. The MA/MO_S3 and MA/MO_S4 samples were combined into MA/MO_S3-S4 during RNA-seq. **(B)** PCA analysis of transcriptome data of litchi flowers.

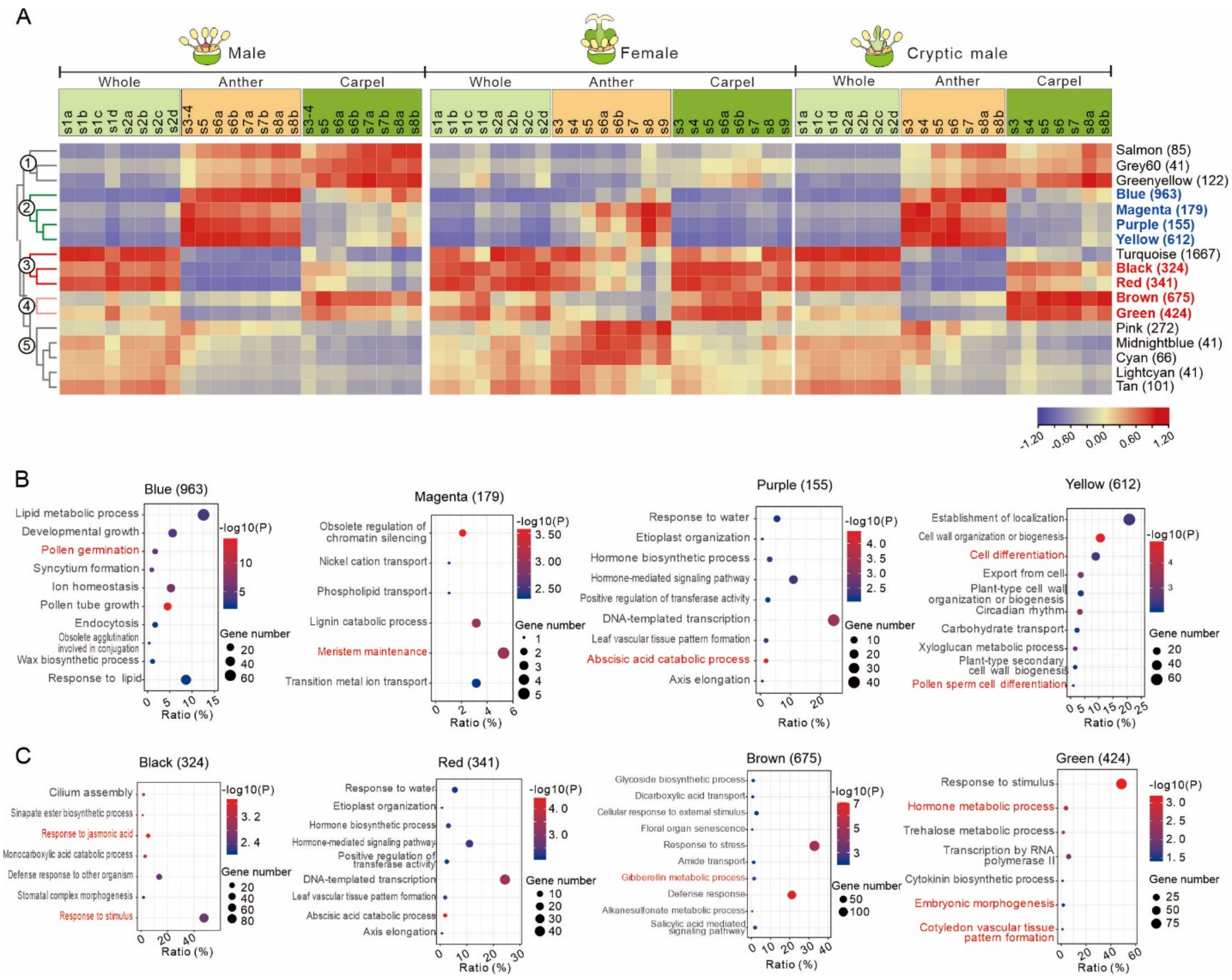


**Fig. S3. Weighted gene correlation network analysis (WGCNA) of flower development in litchi.**

**(A)** Pearson correlation coefficient between gene expression of each module for different flower tissue samples. The modules related to the development of ovary and anther were highlighted in red and blue, respectively. Gene Ontology (GO) enrichment analysis for genes in the modules associated with carpel development **(B)** and anther development **(C)**. The GO term highlighted in red may be involved in the regulation of ovary and anther development.

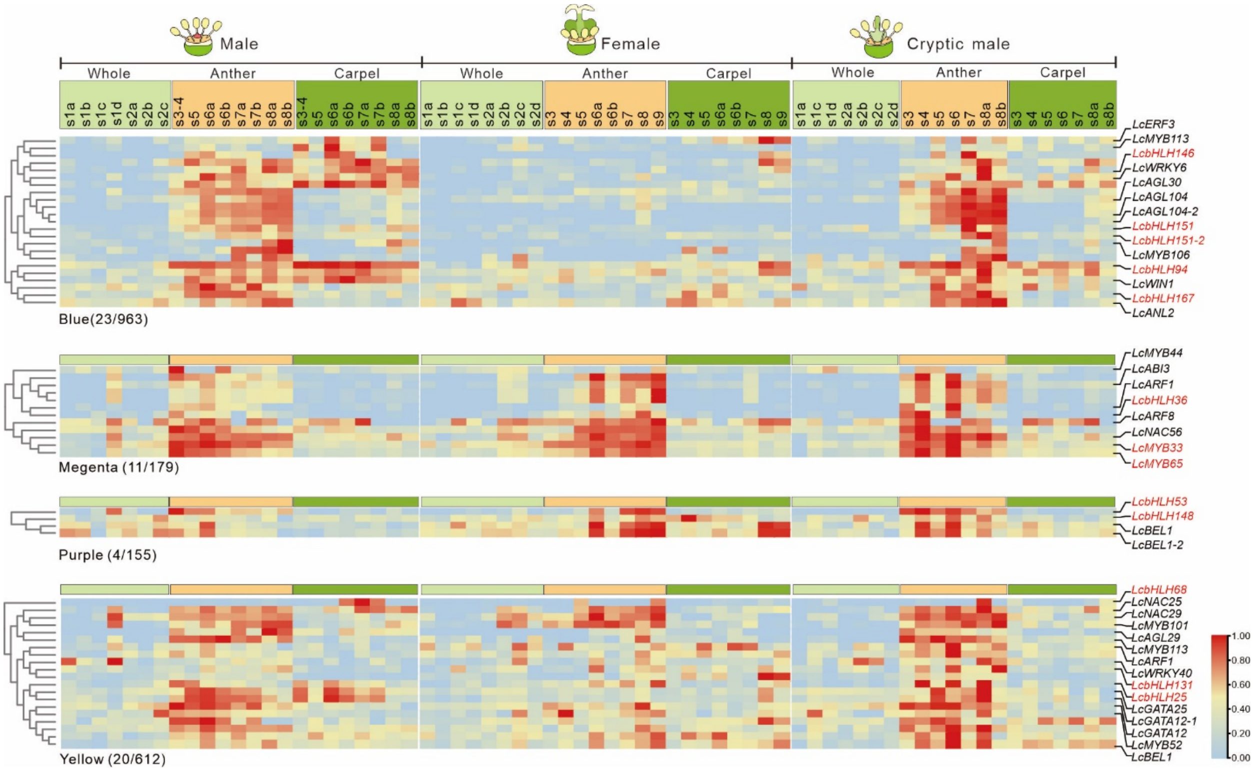


**Fig. S4. Expression pattern of transcription factors from modules associated with anther development.** Genes including *GAMYB* (*LcMYB33* and *LcMYB65*), and bHLH factors are highlighted in red.

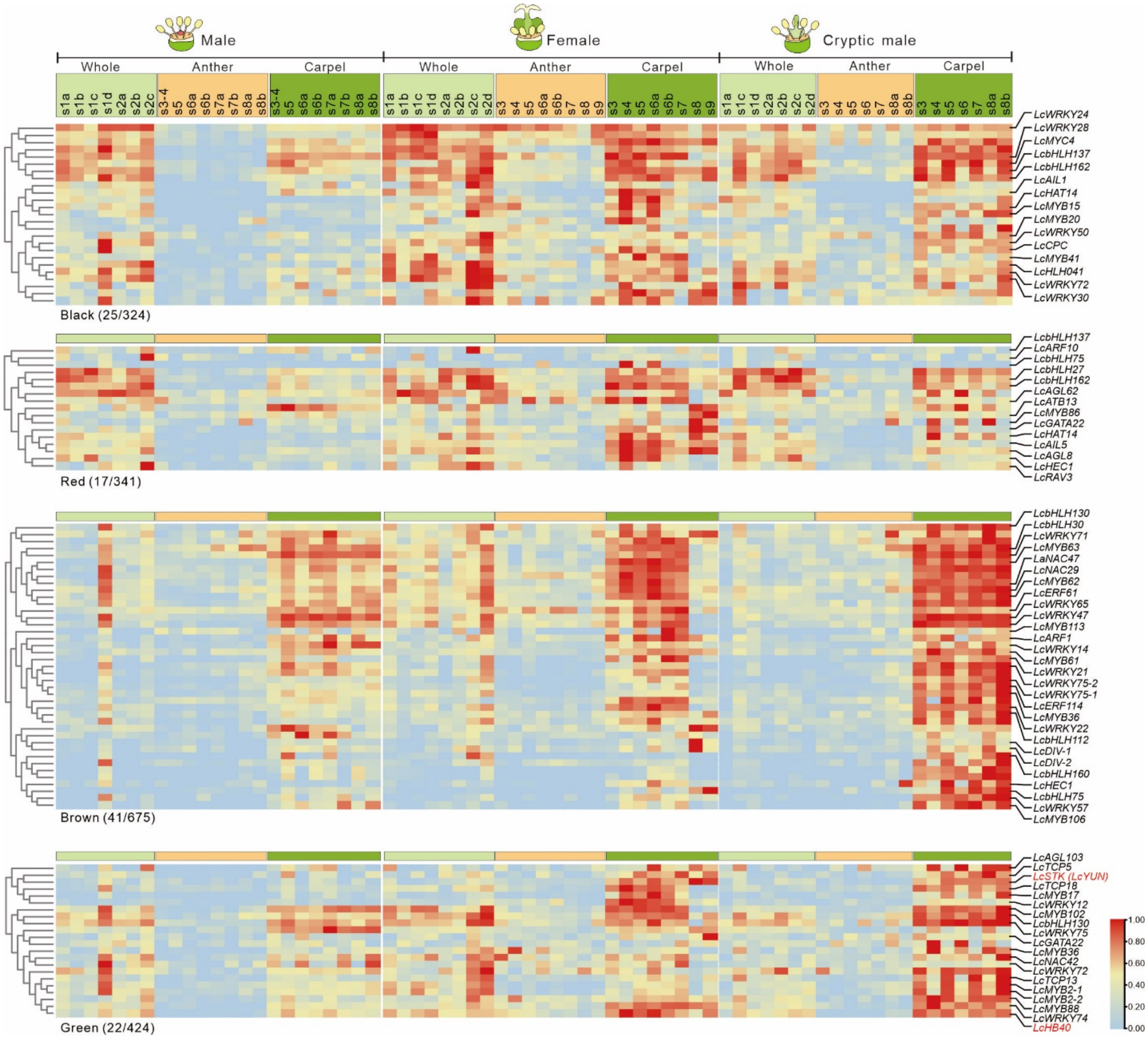


**Fig. S5 Expression pattern of transcription factors from modules associated with carpel development.** Genes including *LcYUN* and *LcHB40* (*LcKUN*) are highlighted in red.

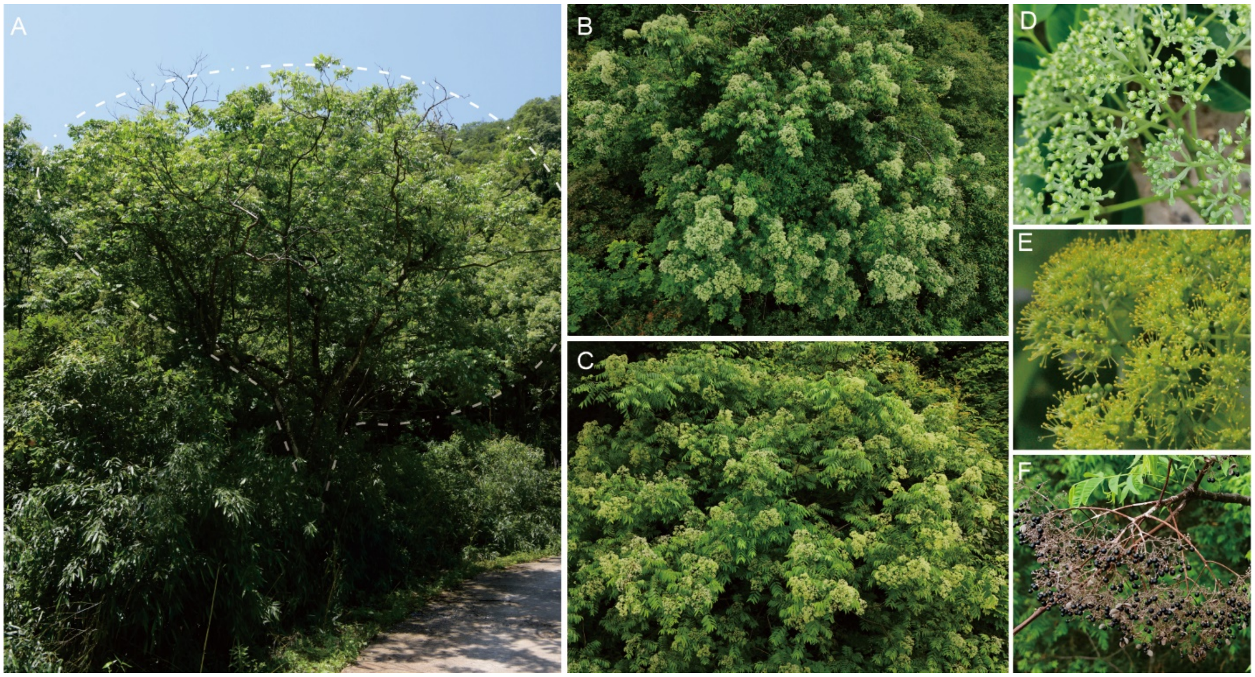


**Fig. S6. Tree, inflorescence and fruit of *E. cavaleriei***

**(A)** Mature *E. cavaleriei* tree can grow up to 3 to 5 meters tall (dashed circle indicates representative individual). **(B–C)** Female plant (B) and male plant (C) trees in full bloom. **(D–E)** Inflorescences of female (D) and male (E). **(F)** Fruiting branch of *E. cavaleriei*, showing dehisced pericarp and releasing glossy black seeds.

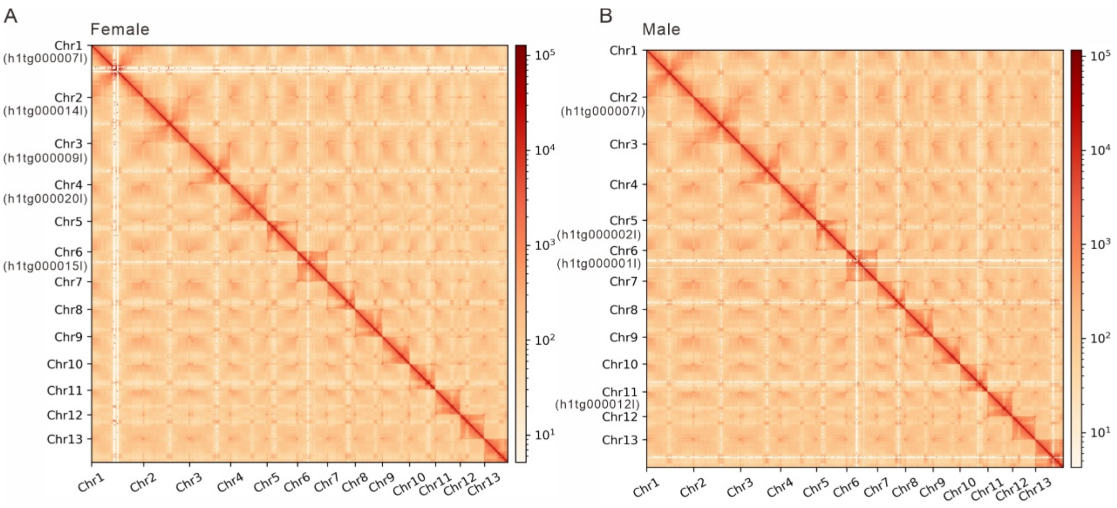


**Fig. S7 Hi-C contact heatmap of *E. cavaleriei*.**

Thirteen pseudochromosomes anchored with Hi-C data in female genome **(A)** and male genome **(B)**. Five chromosomes in female genome and four chromosomes in male genome consist of only one contig.

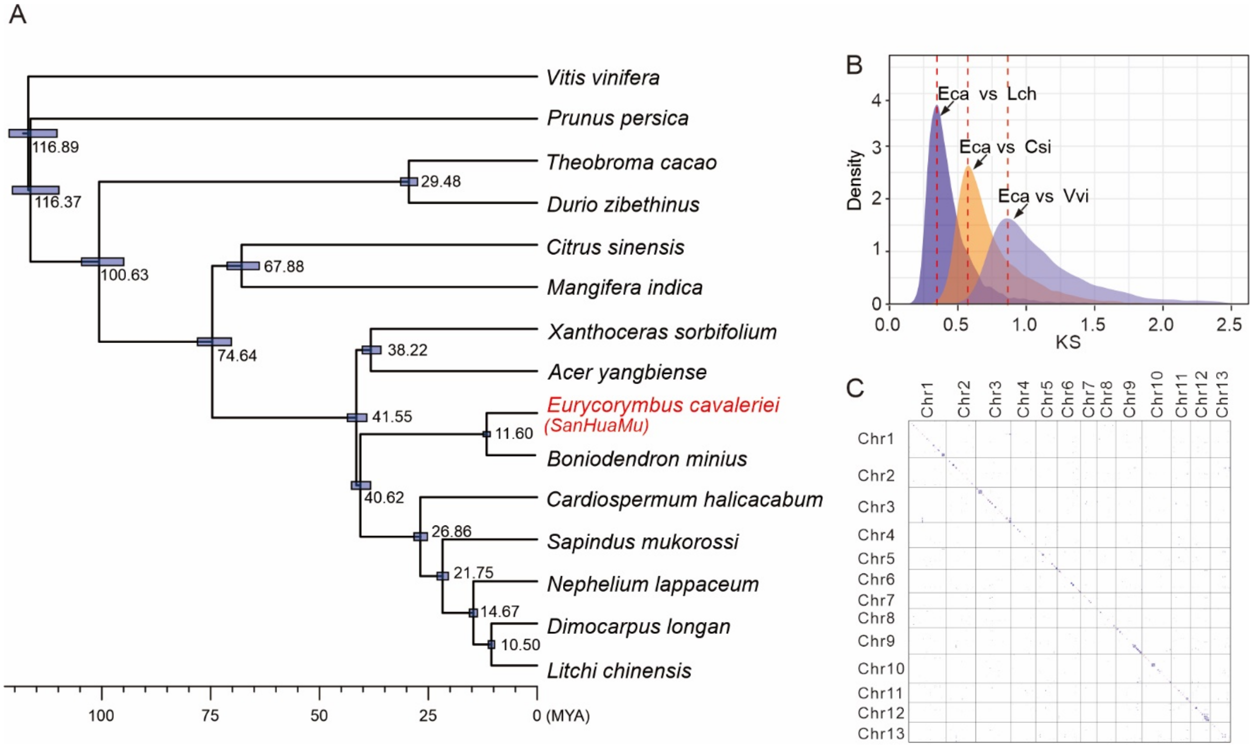


**Fig. S8 Phylogenetic relationship between *E. cavaleriei* and Sapindaceae species.**

**(A)** Phylogenetic tree including the divergence time estimation. (**B**) The density distribution of synonymous nucleotide substitutions per synonymous site (Ks) between orthologs. Eca: *E. cavaleriei*; Lch: *L. chinensis*; Csi: *Citrus* × *sinensis*; Vvi: *V. vinifera* (**C**) The dot plot illustrating the collinearity of paralogous genes in *E. cavaleriei* indicates that there was no independent whole-genome duplication event.

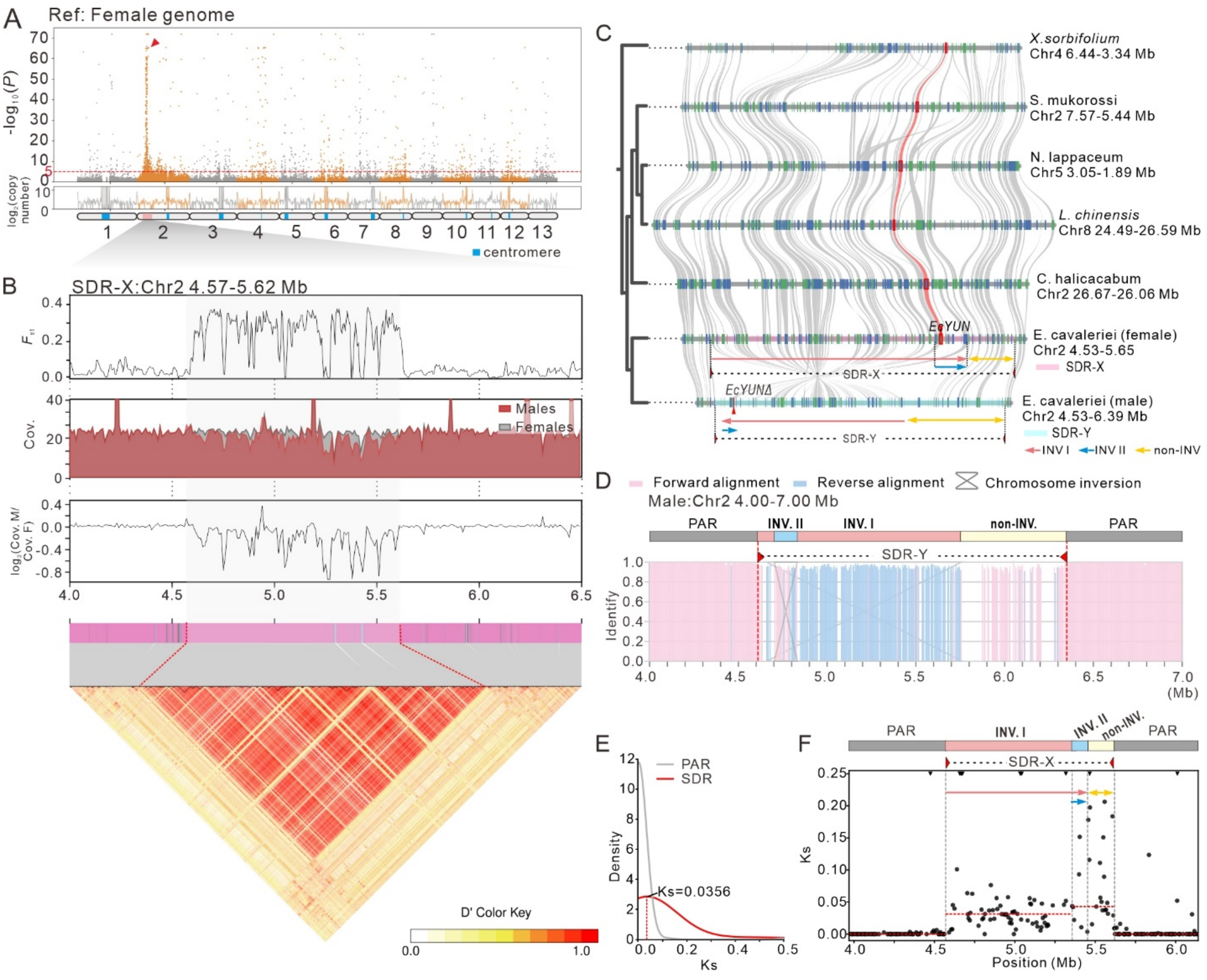


**Fig. S9 Analysis of the sex-determining region.**

**(A)** Using the female genome as a reference, 147 samples were subjected to a genome-wide association study (GWAS) analysis. The red dashed line indicates a *p-value* of $10^{-5}$. An association peak significantly associated with sex type is shown on the chromosome 2 of female genome. **(B)** The analysis of fixation index (*F*st), coverage distribution, and linkage disequilibrium (LD) along the SDR-X. **(C)** Collinearity analysis of the homologous region of SDR with other Sapindaceae species. **(D)** Structure of the SDR. The SDR-X, divided into fragment sequences (length: 500 bp), were mapped to the SDR-Y. And their mapped position and direction (pink for forward; blue for reverse) of these fragments are presented. A large inversion (INV I) occurs in SDR-Y, with most fragments showing large-scale reverse alignment (blue areas). Another inversion (INV II) can be detected, as a number of fragments show forward alignment within INV I (pink areas within blue areas). **(E)** The density distribution of Ks between the homologous gene pairs in SDR and PAR regions. **(F)** Synonymous site divergence values (Ks) between homologous gene pairs in the SDR-X and PAR regions.

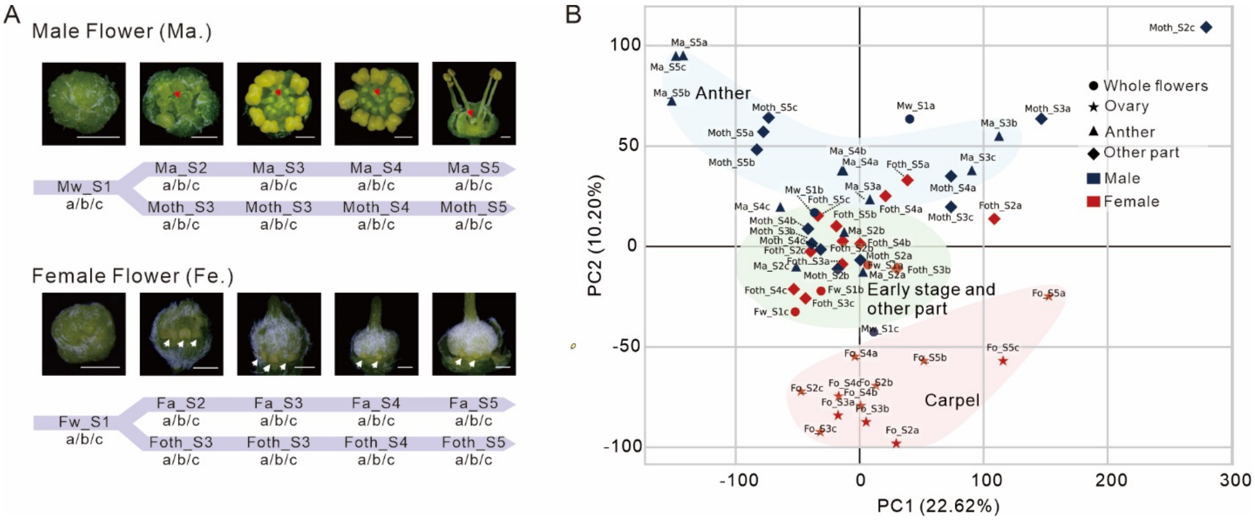


**Fig. S10 Transcriptome analysis of flower in *E. cavaleriei*.**

**(A)** Two types of flowers were collected at six different developmental stages. Whole flower buds were retained stage 1 and 2, while female flowers in stage 3 to 6 were divided into ovary and other tissues, and male flowers in stage 3 to 6 were divided into anther and other tissues. The red arrows in male flowers indicate abortive carpels, while the white arrows in female flowers denote abortive anthers. **(B)** PCA analysis of transcriptome data of *E. cavaleriei* flowers.

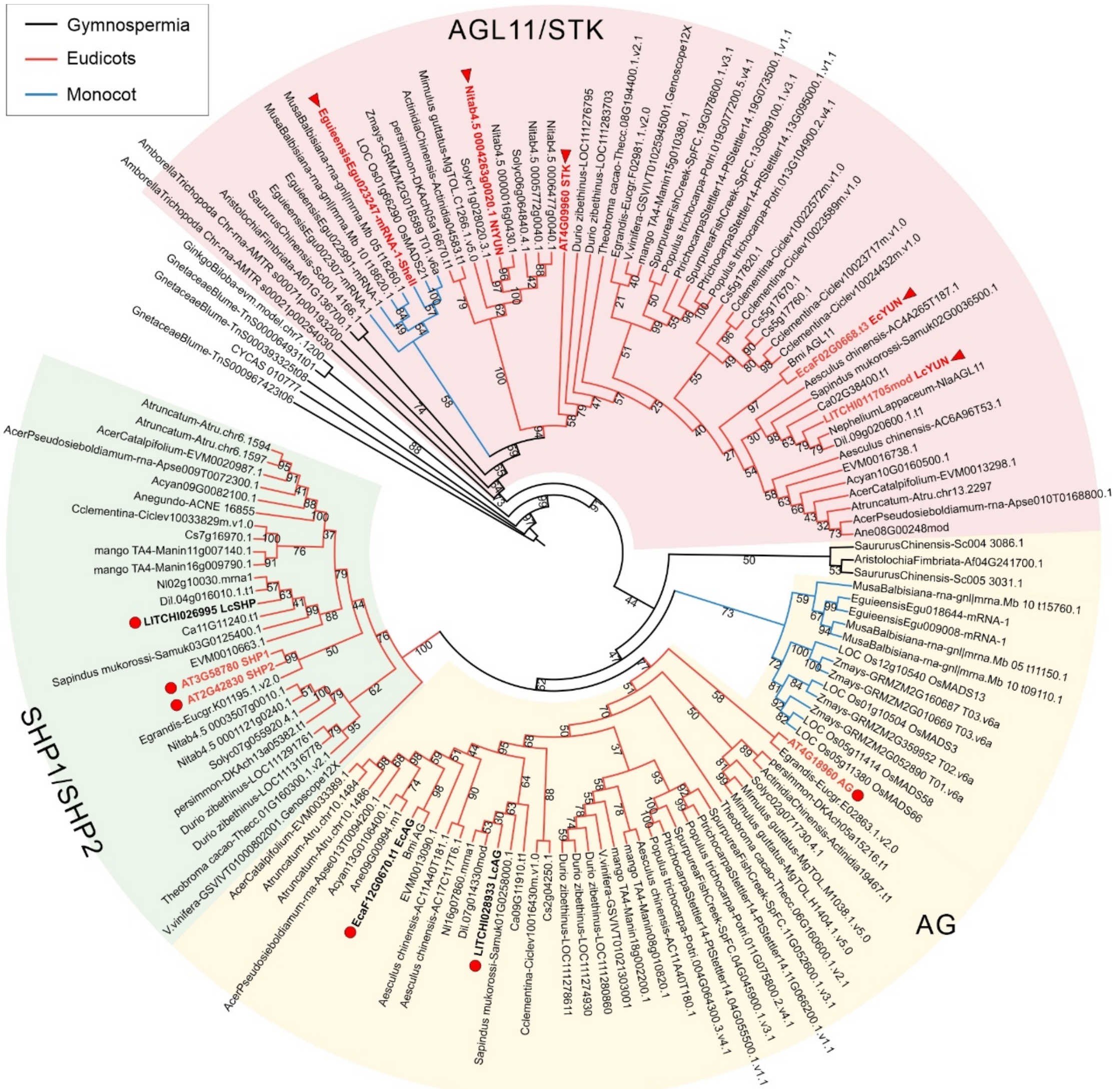


**Fig. S11 Phylogenetic tree of the *AG*-subfamily (consisting of *AG, SHP1/SHP2,* and *AGL11* in *Arabidopsis*).**

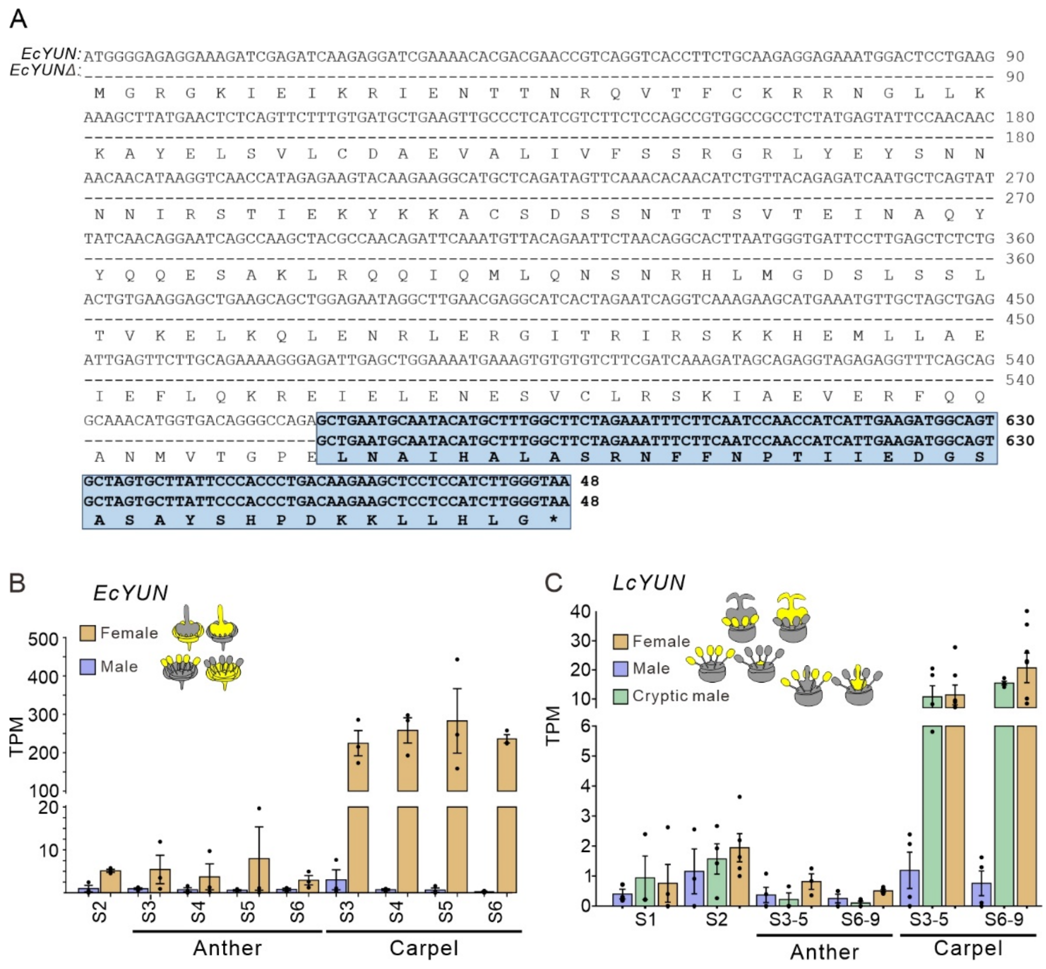


**Fig. S12 Comparison of *EcYUN* in SDR-X and SDR-Y of *E. cavaleriei*.**

**(A)** Sequence comparison between the X-linked *EcYUN* and the Y-linked *EcYUNΔ* in *E. cavaleriei*.

(**B-C)** The expression levels (TPM; transcripts per million) of *EcYUN* (B) and *LcYUN* **(**C**)** in the floral tissues at different developmental stages.

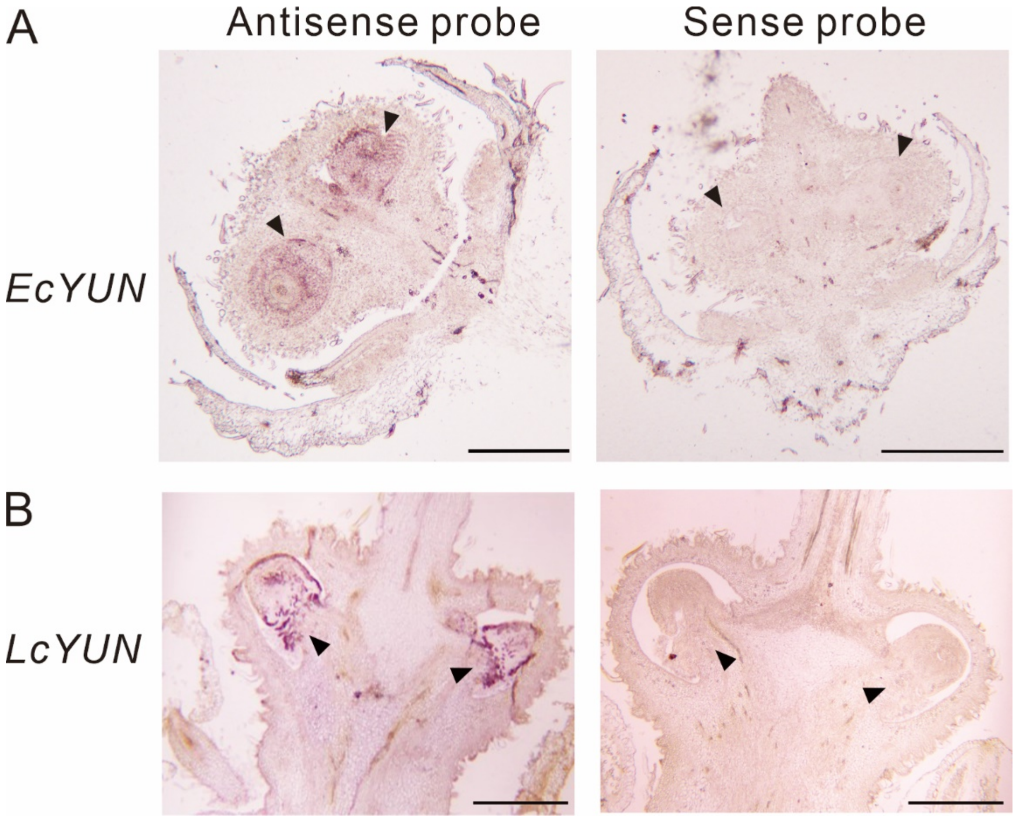


**Fig. S13 *In situ* hybridization of *YUN* in carpel of female flowers**

**(A)** *In situ* hybridization of *EcYUN* in carpel of female flowers at stage 5 in *E. cavaleriei*. (**B)** *In situ* hybridization of *LcYUN* in carpel of female flowers at stage 8. Position of hybridization signals in ovaries are indicated by dark arrows. The sense probe was hybridized as a control (shown on the right). Scale bar: 1 mm.

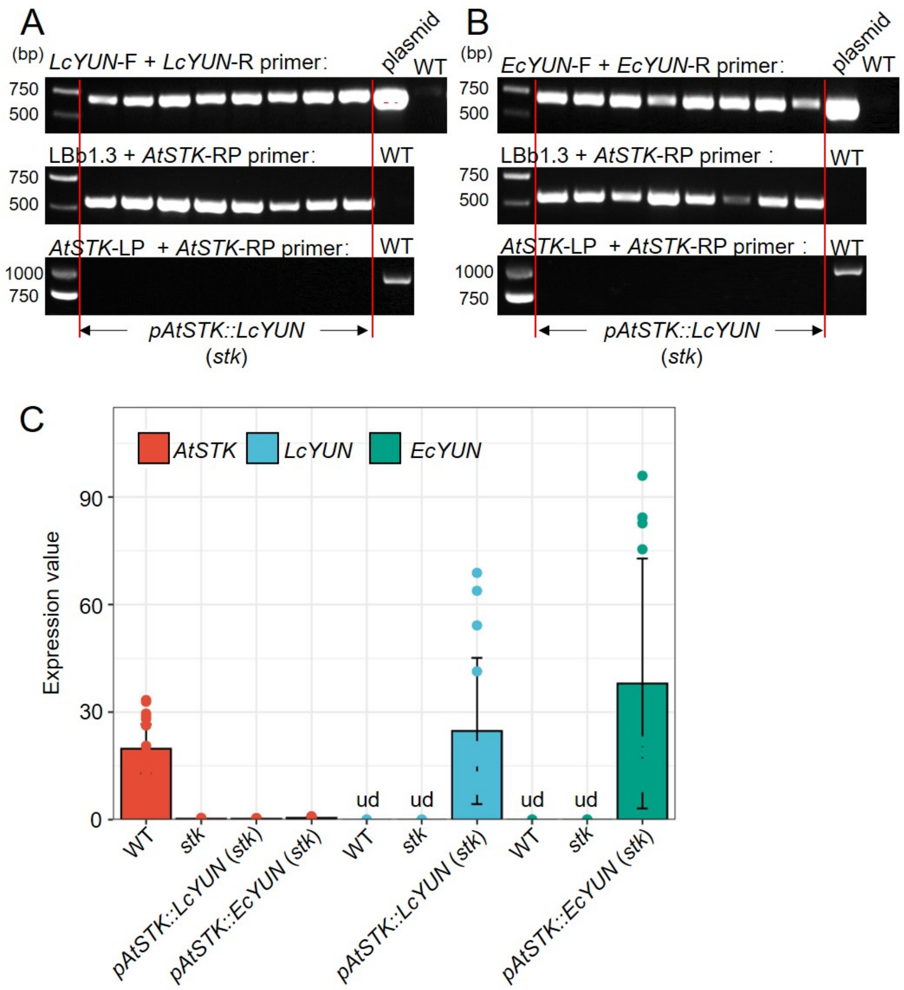


**Fig. S14 Genotyping and expression analysis of *LcYUN* and *EcYUN* in *Arabidopsis* transgenic lines.**

**(A)** PCR analysis of the lithci *LcYUN* and *AtSTK* genes in transgenic plants overexpressing *LcYUN* driven by the *AtSTK* promoter in the *stk* background, with wild-type (WT) plants serving as a negative control. The plasmid containing *pAtSTK::LcYUN* served as a positive control for detecting plants using the *LcYUN*-F and *LcYUN*-R primers. **(B)** PCR analysis of *E. cavaleriei EcYUN* and *AtSTK* genes in transgenic plants overexpressing *EcYUN* driven by the *AtSTK* promoter in the *stk* background, with wild-type (WT) plants serving as a negative control. The plasmid containing *pAtSTK::EcYUN* served as a positive control for detecting plants using the *EcYUN*-F and *EcYUN*-R primers. **(C)** Relative expression of *AtSTK*, *LcYUN* and *EcYUN* in developing siliques across the indicated genotypes: wild-type (WT), *stk* mutant, and *pAtSTK::LcYUN* or *pAtSTK::EcYUN* transgenic lines. Results shown are means ± standard error (SE, n = 4). “ud” denotes that *LcYUN* and *EcYUN* were undetected in the wild-type and *stk* mutant.

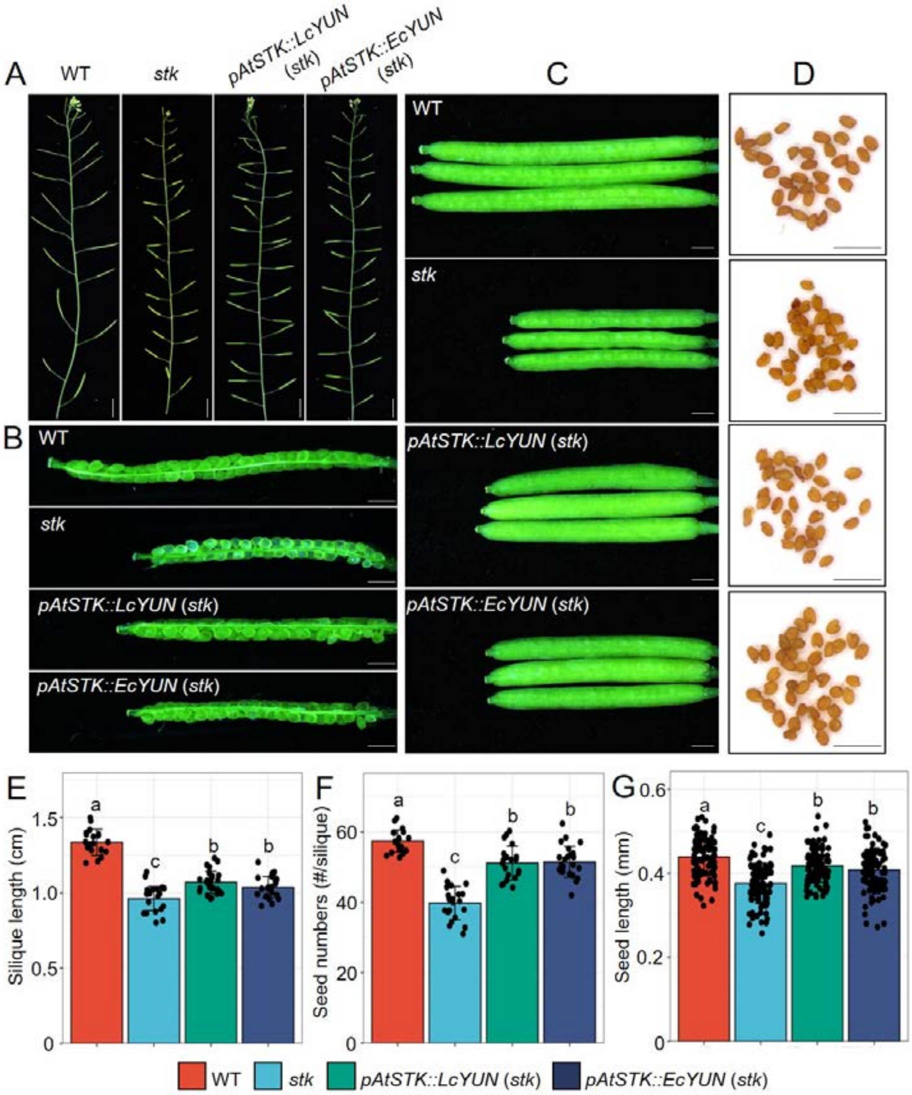


**Fig. S15 Functional complementation of *stk* mutant by overexpressing *LcYUN* and *EcYUN*.**

**(A)** Photographs showing siliques on the main branch of wild type (WT), *stk*, *pAtSTK::LcYUN* and *pAtSTK::EcYUN* plants in the *stk* background. Bars = 1 cm. **(B)** A representative open silique from WT, *stk*, *pAtSTK::LcYUN* and *pAtSTK::EcYUN* plants in the *stk* background, respectively. Bars = 1 mm. **(C)** Fully elongated siliques from WT, *stk*, *pAtSTK::LcYUN* and *pAtSTK::EcYUN* plants in the *stk* background, respectively. Bars = 1 mm. **(D)** Seeds from WT, *stk*, *pAtSTK::LcYUN* and *pAtSTK::EcYUN* plants in the *stk* background, respectively. Bars = 1 mm. **(E)** Quantification of silique length among the genotypes indicated in A-D. Results shown are means ± standard error (SE, n > 20). **(F)** Quantification of seed numbers among the genotypes indicated in A-D. Results shown are means ± standard error (SE, n > 20). (**G)** Quantification of seed length among the genotypes indicated in A-D. Results shown are means ± standard error (SE, n = 100). Different letters in E-G indicate significant differences among different materials for the same trait in a two-way ANOVA test ($P < 0.05$).

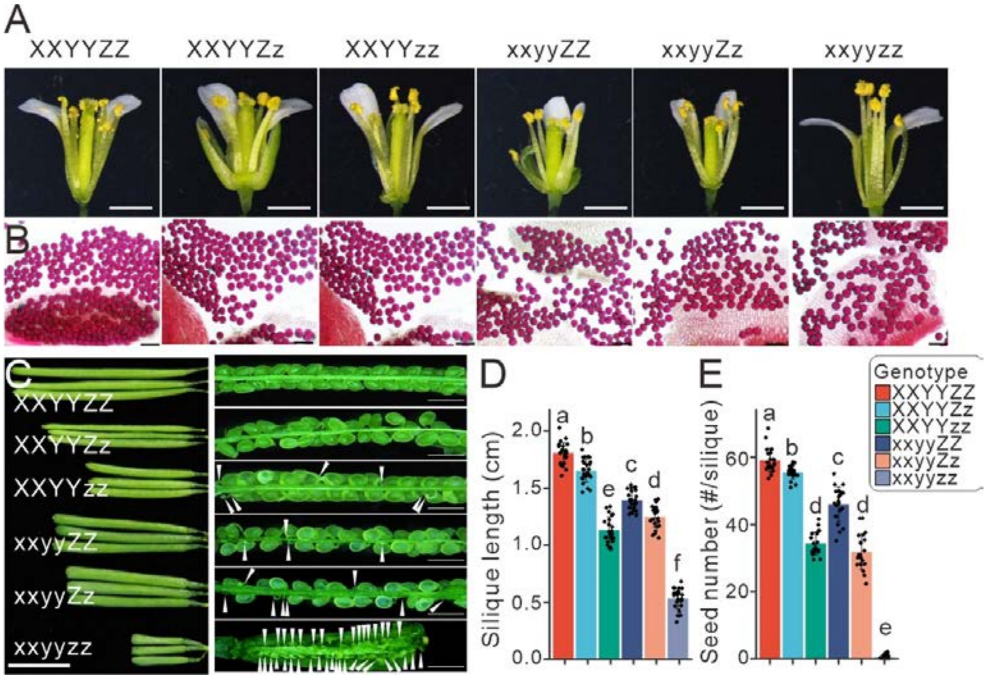


**Fig. S16 Phenotypic characterization of flowers and seeds in mutants of *SHP1*, *SHP2*, and *STK* genes in *Arabidopsis*.**

**(A)** Photographs showing the opened flowers among different combinations of *SHP1*, *SHP2*, *STK* mutants. For clarity, in the figure, X denotes the *SHP1* gene, Y represents the *SHP2* gene, and Z indicates the *STK* gene. These mutants were F2 progeny derived from the hybridization of the *shp1 shp2* double mutant with the *stk* mutant. Bars = 1 mm. **(B)** Alexander staining of pollen grains from different mutants indicated in A. Bars = 50 µm. **(C)** Fully elongated siliques and a representative open silique of different mutants indicated in (A). Abnormal seeds are indicated by white arrowheads. Bars = 0.5 cm. **(D)** Quantification of silique length from different mutants. Results shown are means ± standard error (SE, n = 27). **(E)** Quantification of seed numbers from different mutants. Results shown are means ± standard error (SE, n = 22). Different letters indicate significant differences among different mutants for the same trait in a two-way ANOVA test ($P < 0.05$).

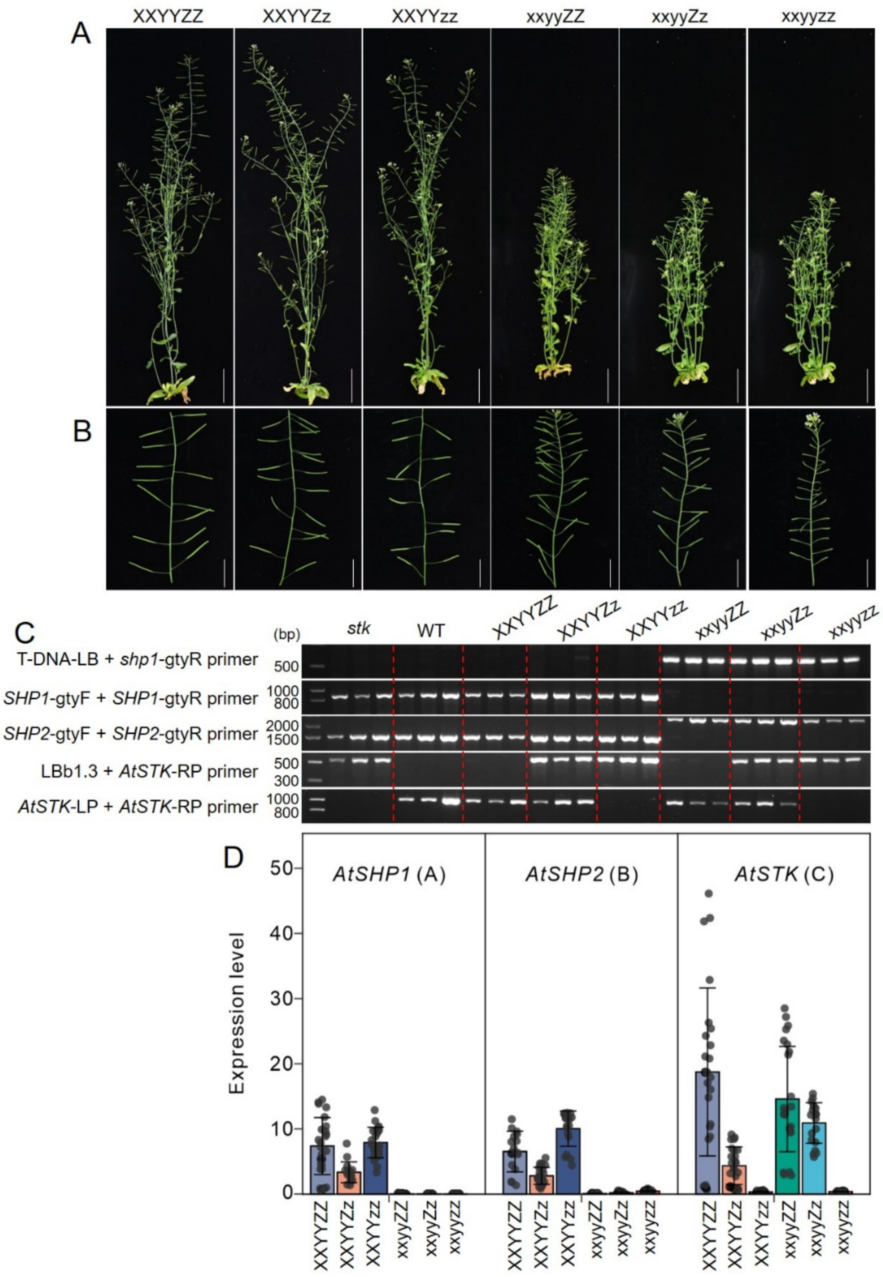


**Fig. S17 Phenotypic and molecular identification of mutants of *SHP1, SHP2,* and *STK* genes in *Arabidopsis*.**

**(A)** Photographs showing *Arabidopsis* plant growth among different mutants of *SHP1*, *SHP2*, and *STK* genes at 6 WAG (weeks after germination). For clarity, in the figure, X denotes the *SHP1* gene, Y represents the *SHP2* gene, and Z indicates the *STK* gene. These mutants were F2 progeny derived from the hybridization of the *shp1 shp2* double mutant with the *stk* mutant. Bars = 5 cm. **(B)** Photographs displaying partial siliques on the main branch of different mutants. Bars = 2 cm. **(C)** PCR analysis of the *AtSHP1*, *AtSHP2*, and *AtSTK* genes in *stk*, wild type (WT) and all mutants as indicated in (A). **(D)** Relative expression of *AtSHP1*, *AtSHP2*, and *AtSTK* in developing siliques of different mutants, Results shown are means ± standard error (SE, n > 6).

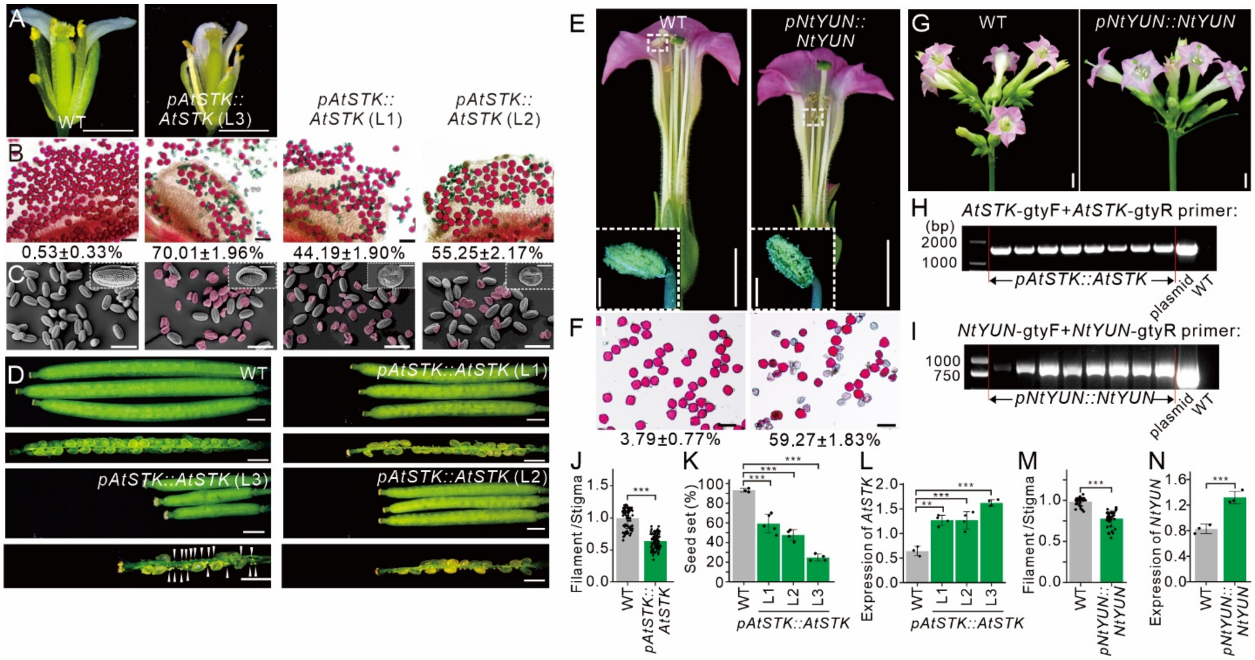


**Fig. S18 Overexpression of *YUN* results in defective anthers and seeds in Arabidopsis and tobacco.**

**(A)** The opened flowers from the wild type (WT) and the *pAtSTK::AtSTK* transgenic *Arabidopsis* line 3. Bars = 1 mm. **(B)** Alexander staining of pollen grains from WT and three independent *pAtSTK::AtSTK* transgenic lines. Bars = 50 μm. The percentage of abnormal pollen grains is shown at the bottom of each transgenic lines. Results are means ± standard error (SE, n > 400). **(C)** SEM images of mature pollen grains from WT and *pAtSTK::AtSTK* transgenic lines. Abnormal pollen grains are highlighted in red. Bars = 50 μm. **(D)** Fully elongated siliques from WT and *pAtSTK::AtSTK* transgenic line. Bars = 1 mm. **(E)** The opened flowers from the WT and the *pNtYUN::NtYUN* transgenic tobacco plants. The bottom left corner shows a magnified image of the pollen-releasing anther. Bars = 1 cm. **(F)** Alexander staining of pollen grains from the wild type (left) and the *pNtYUN::NtYUN* transgenic tobacco plants (right). Bars = 50 μm. **(G)** Photographs showing tabacco flowers of WT and *pNtYUN::NtYUN* transgenic plants. Bars = 1 cm. **(H-I)** PCR analysis of the *AtSTK* gene in transgenic *Arabidopsis* plants overexpessing *AtSTK* driven by the *AtSTK* promoter (H) and the *NtYUN* gene in transgenic tobacco plants overexpressing *NtYUN* driven by the *NtYUN* promoter (I), respectively. The plasmid harboring *pAtSTK::LcYUN* and *pNtYUN::NtYUN* served as a positive control when detecting the plants using the *AtSTK*-gtyF/*AtSTK*-gtyR and *NtYUN*-gtyF/*NtYUN*-gtyR primers, respectively. WT of *Arabidopsis* and tobacco served as a negative control. **(J)** The ratio of filament length to stigma length in the wild type and the *pAtSTK::AtSTK* transgenic plant. **(K)** Quantification of seed set from WT and *pAtSTK::AtSTK* transgenic lines. **(L)** Expression levels of *AtSTK* in developing siliques of WT and *pAtSTK::AtSTK* transgenic lines. Results shown are means ± standard error (SE, $n \geq 3$). (M) The ratio of filament length to stigma length in the wild type and the *pNtSTK::NtSTK* transgenic tobacco plants. Results shown are means ± standard error (SE, $n > 200$). **(N)** The relative expression levels of *NtYUN* in ovary of wild type and *pNtYUN::NtYUN* transgenic tobacco plants. Results shown are means ± standard error (SE, $n = 3$). Asterisks of J-N represent significant differences in Student's *t* tests (***, $P \leq 0.001$).

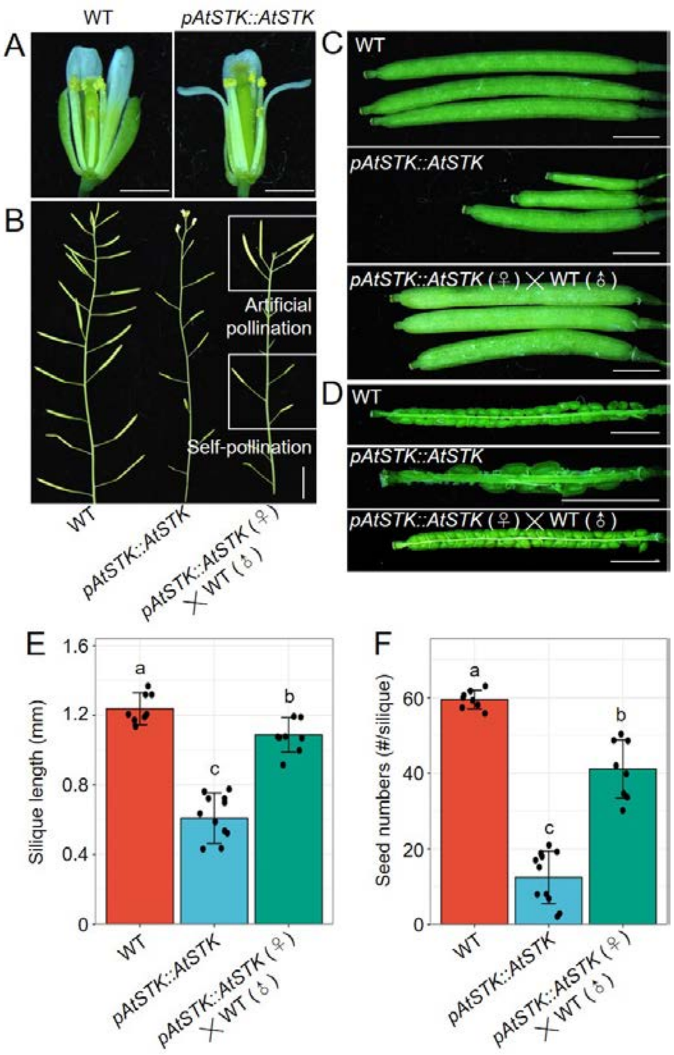


**Fig. S19 The silique and seed defects observed in the *pAtSTK::AtSTK* transgenic line can be partially rescued by artificial pollination with wild-type pollen.**

**(A)** The opened flowers from wild type and *pAtSTK::AtSTK* transgenic line used for hybridization. Bars = 1 mm. **(B)** Photographs showing the siliques on branches of wild type (WT) (left), *pAtSTK::AtSTK* transgenic plant (*pAtSTK::AtSTK*) (middle), *pAtSTK::AtSTK* transgenic plant that underwent artificial pollination of wild-type pollen (*pAtSTK::AtSTK* (♀) × WT (♂)) (right). Bars = 1 cm. **(C)** Fully elongated siliques from wild type (WT), *pAtSTK::AtSTK* transgenic plant, and *pAtSTK::AtSTK* (♀) × WT (♂), respectively. Bars = 2 mm. **(D)** A representative open silique from WT, *pAtSTK::AtSTK* and *pAtSTK::AtSTK* (♀) × WT (♂), respectively. Bars = 2 mm. **(E)** Quantification of silique length among the genotypes indicated in C. Results shown are means ± standard error (SE, n ≥ 7). **(F)** Quantification of seed numbers among the genotypes indicated in C. Results shown are means ± standard error (SE, n ≥ 8). Different letters indicate significant differences among different materials indicated for the same trait in a two-way ANOVA test ($P < 0.05$).

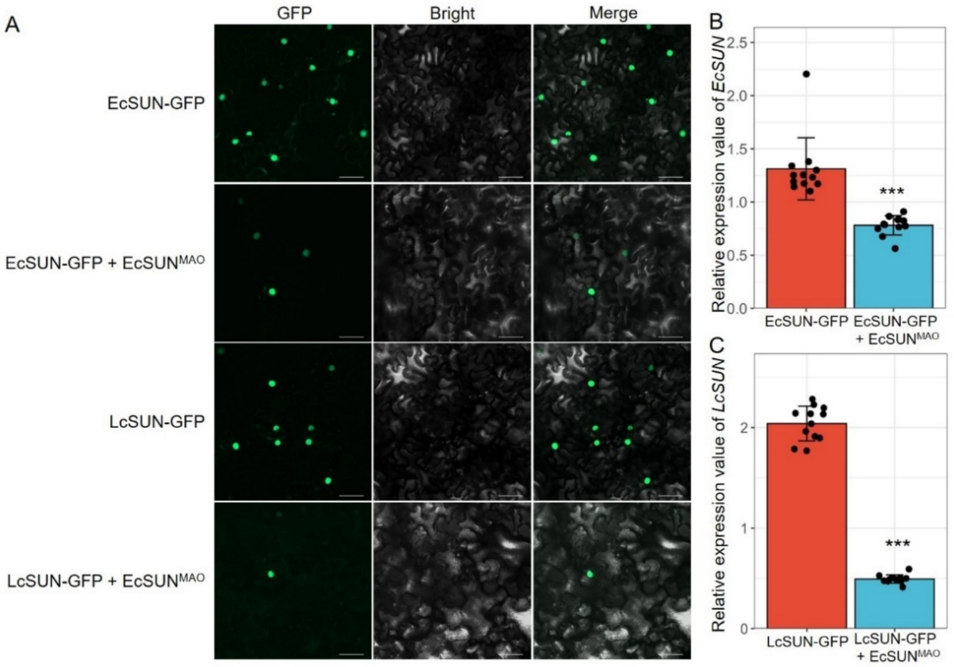


**Fig. S20. The silencing effect of sRNAs generated by the *SUN^MAO^* on the *SUN* gene.**

**(A)** The GFP signal of SUN fused to GFP in tobacco cells. Tobacco cells were observed by green GFP fluorescence of SUN protein. When both *SUN*-GFP and *SUN$^{MAO}$* are introduced simultaneously, the signal of *SUN*-GFP is significantly weakened. Scale bars = 50 μm. **(B)** Relative expression value of *EcSUN* in *EcSUN*-GFP tobacco cells and *EcSUN*-GFP + *EcSUN$^{MAO}$* tobacco cells, respectively. **(C)** Relative expression value of *LcSUN* in *LcSUN*-GFP tobacco cells and *LcSUN*-GFP + *EcSUN$^{MAO}$* tobacco cells, respectively. Results shown are means ± standard error (SE, n = 12). Significant differences (*P*) between the different treatments were assessed using Student's *t*-tests (***, $P \leq 0.001$).

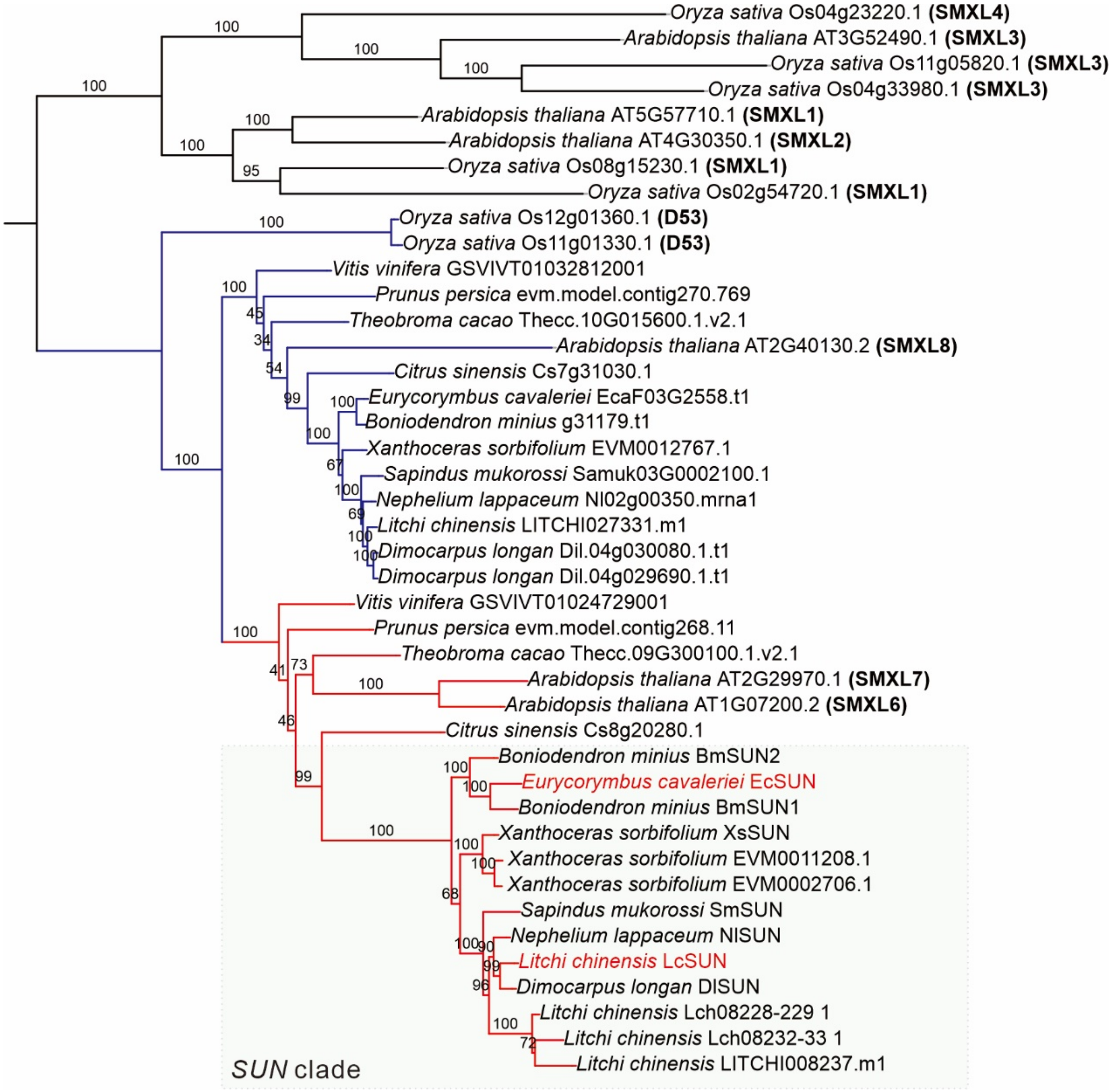


**Fig. S21 Phylogenetic tree of *SMXL* homologous genes.** The colored branches in the phylogenetic tree represent *SMXL6/7/8* orthologs from 13 species. The clade with *SUN* homologous genes were labeled by red.

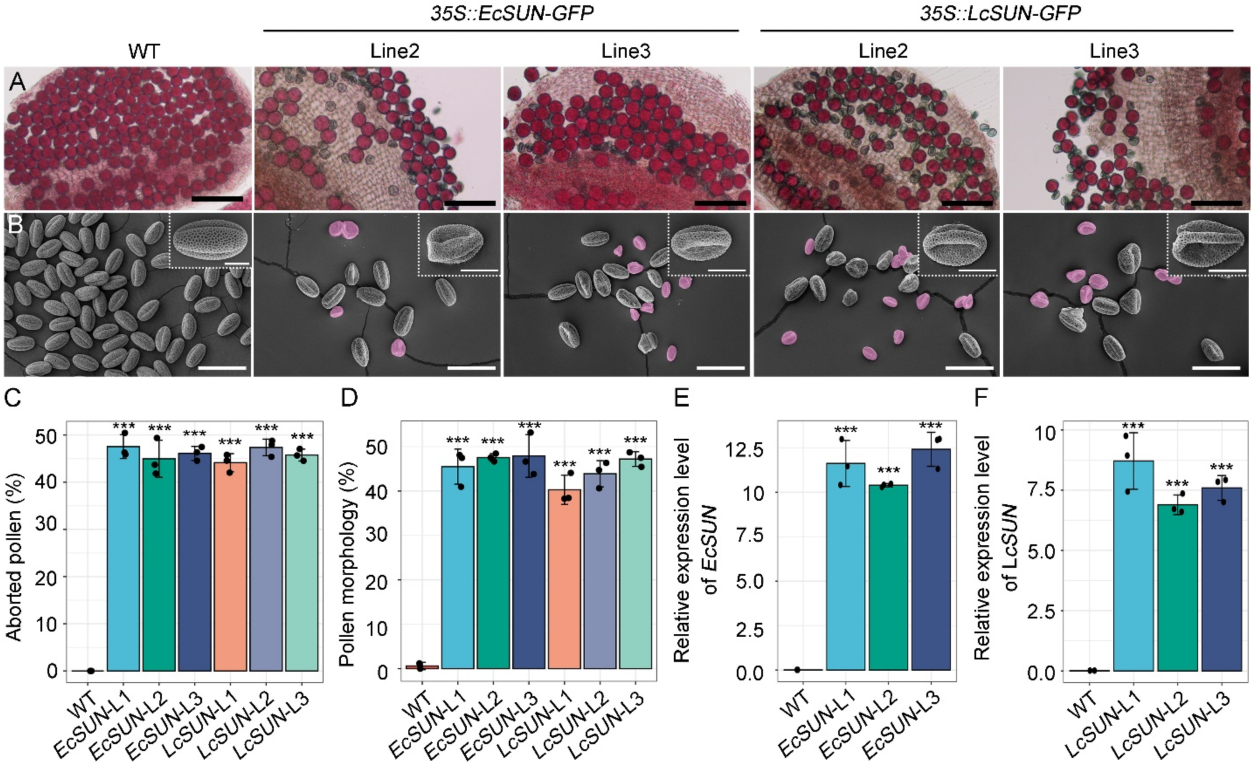


**Fig. S22 Overexpression of either *EcSUN* and *LcSUN* leads to male sterility in *Arabidopsis*.**

**(A)** Alexander staining of mature pollen grains from wild type, two lines of *35S::EcSUN*-GFP and *35S::LcSUN*-GFP transgenic plants. **(B)** Scanning electron micrographs (SEMs) of pollen grains, along with close-up images of a single pollen grain from wild-type plants and two lines of 35S::*EcSUN*-GFP and 35S::*LcSUN*-GFP transgenic plants. Abnormal pollen grains are highlighted in purple. Bars = 50 µm for (A-B); Bars = 10 µm for enlarged view of a pollen grain in (B). **(C)** Percentage of aborted pollen determined by Alexander staining (A). **(D)** Percentage of defective pollen morphology based on SEM (B). **(E-F)** Relative expression level of *EcSUN* **(E)** and LcSUN **(F)** in wild type, three lines of *35S::EcSUN*-GFP and *35S::LcSUN*-GFP transgenic plants. Asterisks represent significant differences between WT and transgenic plants in Student's t tests (***, $P \leq 0.001$).

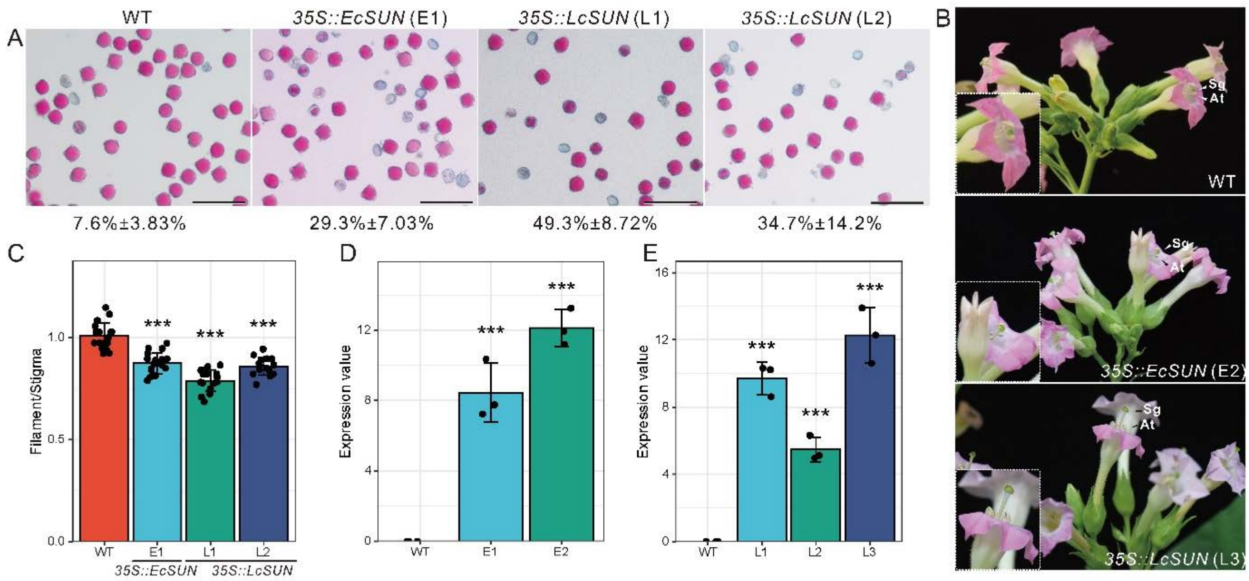


**Fig. S23 Overexpression of either *EcSUN* or *LcSUN* leads to male sterility in tobacco.**

**(A)** Alexander staining of mature pollen grains from wild type and independent lines of *35S::EcSUN* (E1) and *35S::LcSUN* (L1, L2). Bars = 100 μm. Percentages of abnormal pollen grains are shown at the bottom. Results are means ± standard errors (SE, n > 600). (**B)** Photographs showing tobacco flower phenotype in wild type, 35S::*LcSUN* (L3) and 35S::*EcSUN* (E2) transgenic tobacco plants. (**C)** The ratio of filament length to stigma length in the WT, and in the transgenic tobacco plants, *35S::EcSUN* (E1), and *35S::LcSUN* (L1, L2); means ± standard errors are shown. Asterisks represent significant differences between WT, 35S::*LcSUN* and 35S::*EcSUN* transgenic plants in Student's *t* tests (***, $P \leq 0.001$, n > 16). (**D)** The relative expression levels of *EcSUN* in wild type and 35S::*EcSUN* transgenic tobacco plants (E1, E2). Results shown are means ± standard error (SE, n = 3). Asterisks represent significant differences between WT and transgenic plants in Student's *t* tests (***, $P \leq 0.001$,). (**E)** The relative expression levels of *LcSUN* in wild type and 35S::*LcSUN* transgenic tobacco plants (L1, L2, L3). Results shown are means ±

standard error (SE, n = 3). Asterisks represent significant differences between WT and transgenic plants in Student's t tests (***, $P \leq 0.001$).

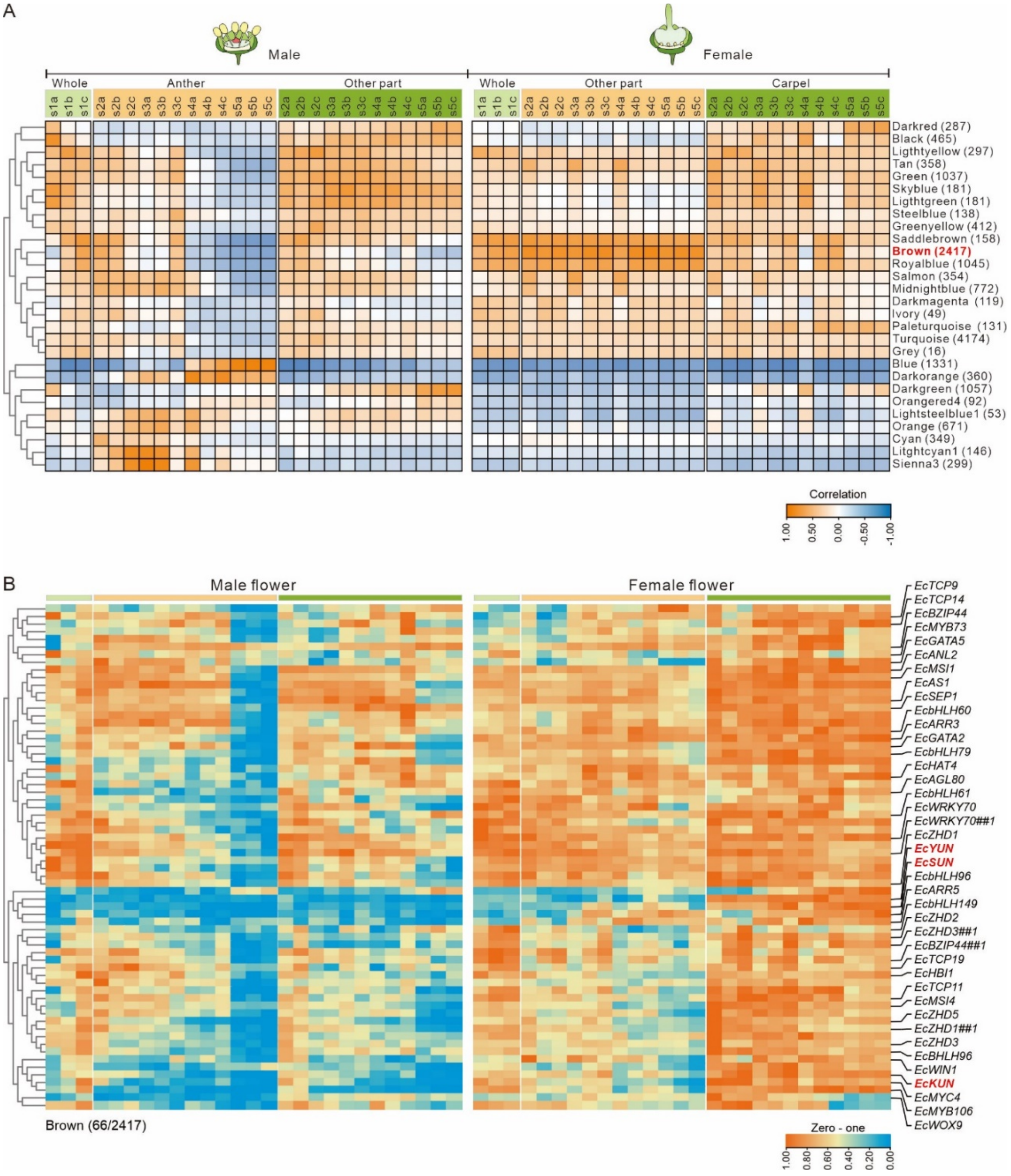


**Fig. S24 Weighted gene correlation network analysis (WGCNA) of flower development in *E. cavaleriei*.**

**(A)** Pearson correlation coefficient between gene expression of each module and different flower samples. (**B)** Expression pattern of transcription factors from modules associated with carpel development. *EcSUN*, *EcKUN*, and *EcYUN* were highlighted in red.

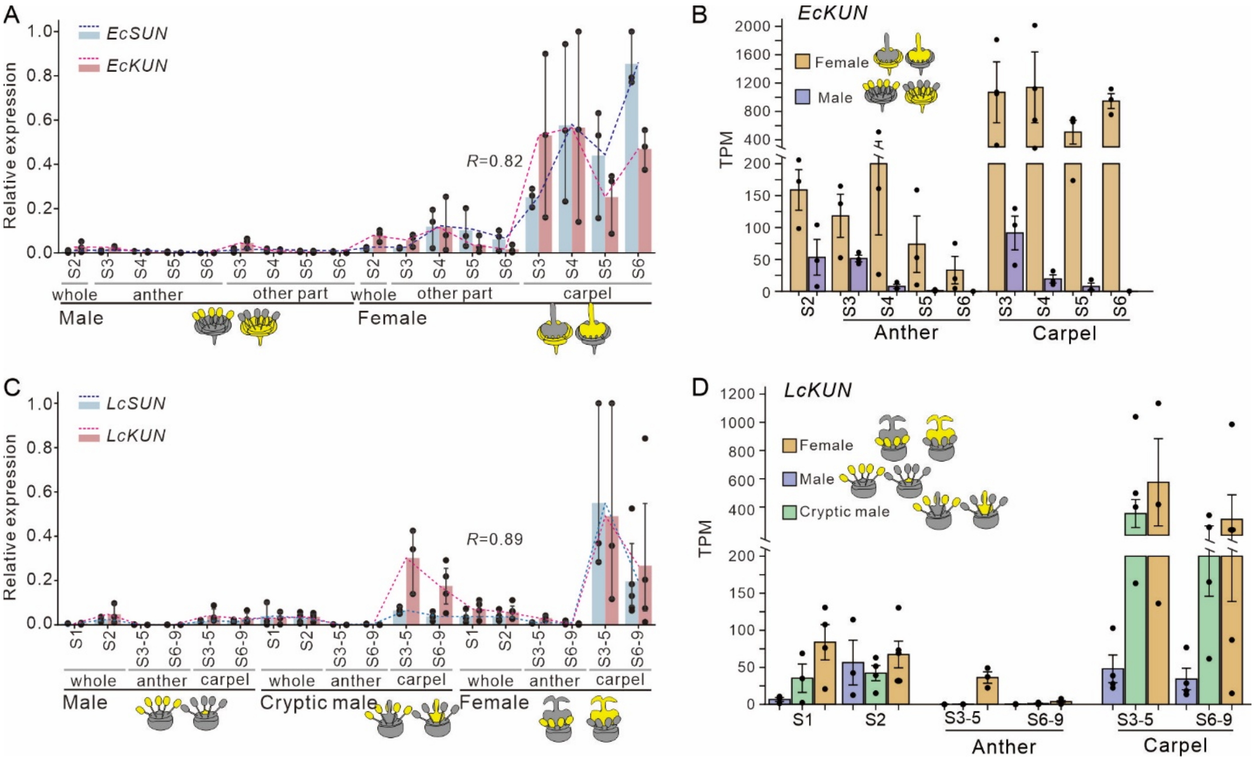


**Fig. S25 Expression of *KUN* across floral development in *E. cavaleriei* and litchi**

**(A, C)** Expression dynamics of *SUN* and *KUN* during floral development in *E. cavaleriei* (A) and litchi (C). Gene expression levels (TPM) were normalized using zero-one scaling (min-max normalization). Pearson correlation coefficients (*R*) were calculated to assess the correlation between *SUN* and *KUN* expression. Dashed lines indicate their expression trends. **(B, D)** Expression level of *KUN* in flower buds of *E. cavaleriei* (B) and litchi (D).

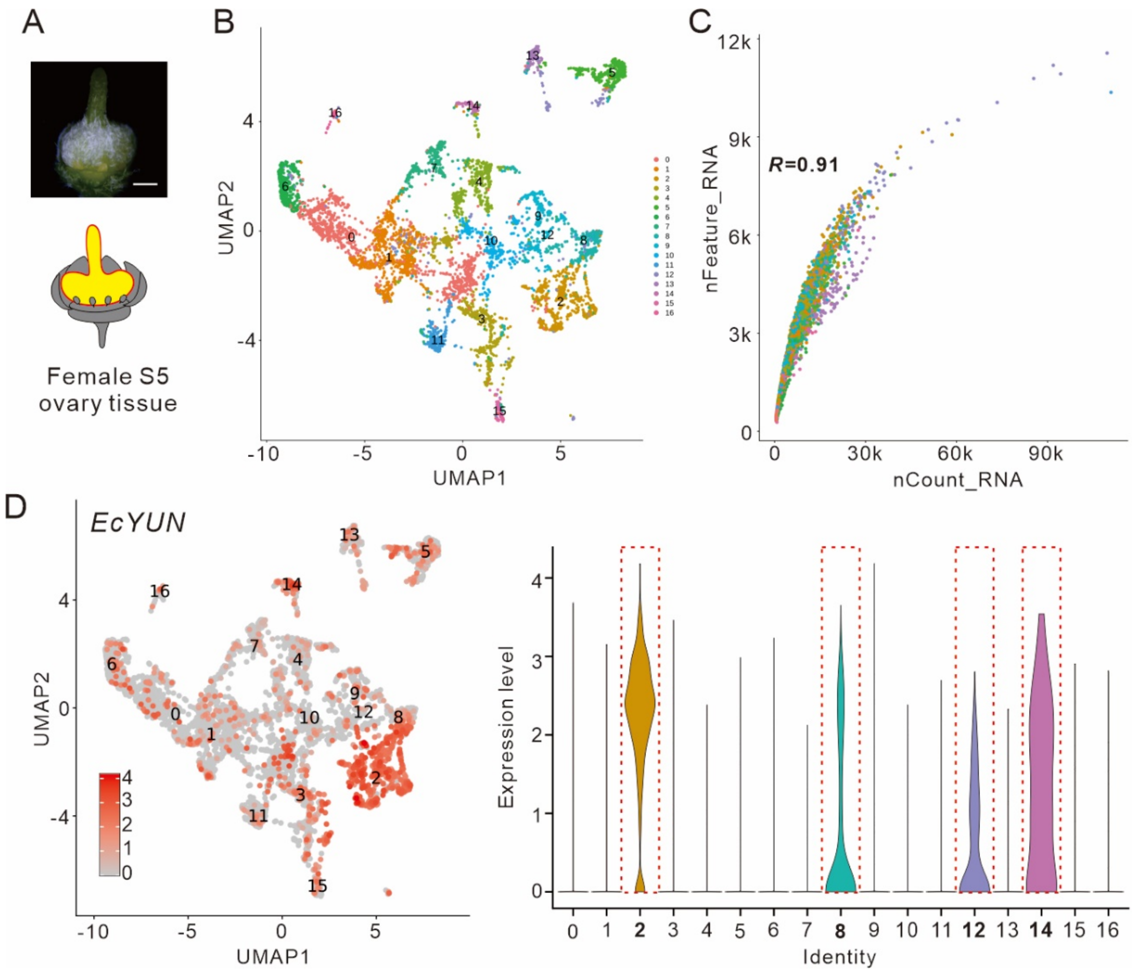


**Fig. S26 snRNA-seq analysis of ovary tissues of *E. cavaleriei* at stage 5.**

**(A)** Nuclei were isolated from stage 5 floral ovaries of *E. cavaleriei*, and subjected to snRNA-seq. **(B)** UMAP analysis of nuclei isolated from the ovary sample by snRNA-seq, colored by 17 cell clusters. **(C)** Positive correlation between gene detection efficiency (nFeature_RNA) and sequencing depth (nCount_RNA) across single nucleus. **(D)** Expression level of *EcYUN* in different cell identities. *EcYUN* showed higher expression in the clusters 2, 8, 12,14 (dotted red rectangle).

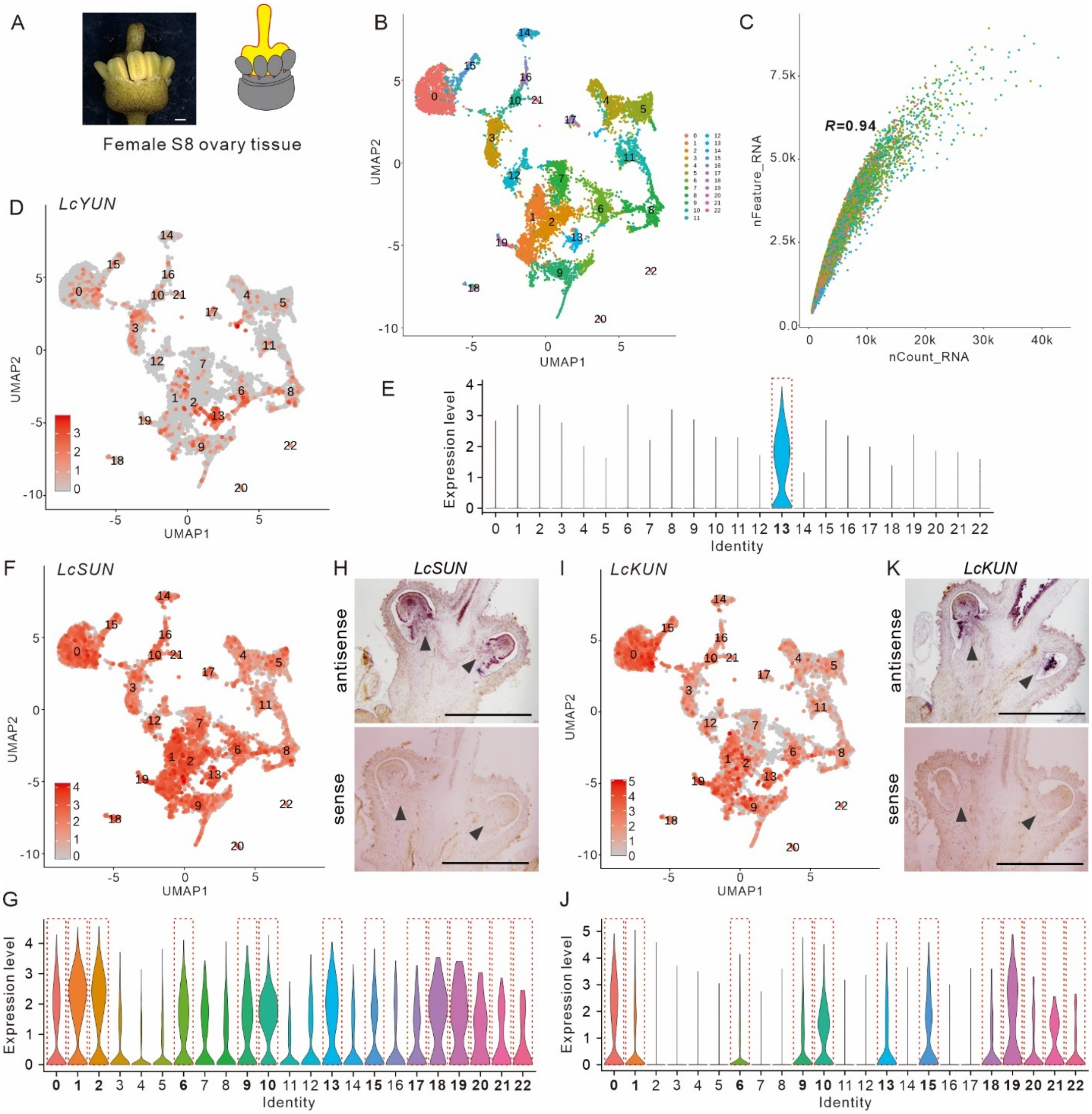


**Fig. S27 snRNA-seq analysis and in situ hybridization of *LcSUN* and *LcKUN* in ovary tissues of litchi at stage 8.**

**(A)** Nuclei were isolated from stage 8 floral ovaries of litchi, and subjected to snRNA-seq. **(B)** UMAP analysis of nuclei isolated from the ovary sample by snRNA-seq, colored by 23 cell clusters. **(C)** Positive correlation between gene detection efficiency (nFeature_RNA) and sequencing depth (nCount_RNA) across single nucleus. **(D-E)** Expression level of *LcYUN* in different cell identities. *LcYUN* showed specific expression in the clusters 13 (dotted red rectangle). **(F-J)** snRNA-seq (F-G, I-J) and in-situ hybridization (H, K) indicates that *LcSUN* and *LcKUN* transcripts are co-localized at the cellular level. Feature plots (F, I) and violin plots (G, J) represent expression of *LcSUN* and *LcKUN* (E-F) in 23 distinct groups. Both *LcSUN* and *LcKUN* are highly expressed in Clusters 0, 1, 6, 9, 10, 13, 15, 18, 19, 20, 21, and 22 (dotted red rectangles). Expression of *LcSUN* (H) and *EcKUN* (K) detected by RNA in situ hybridization assays on S8 female flower buds. The ovule positions are indicated by arrows. Bars = 0.5 mm.

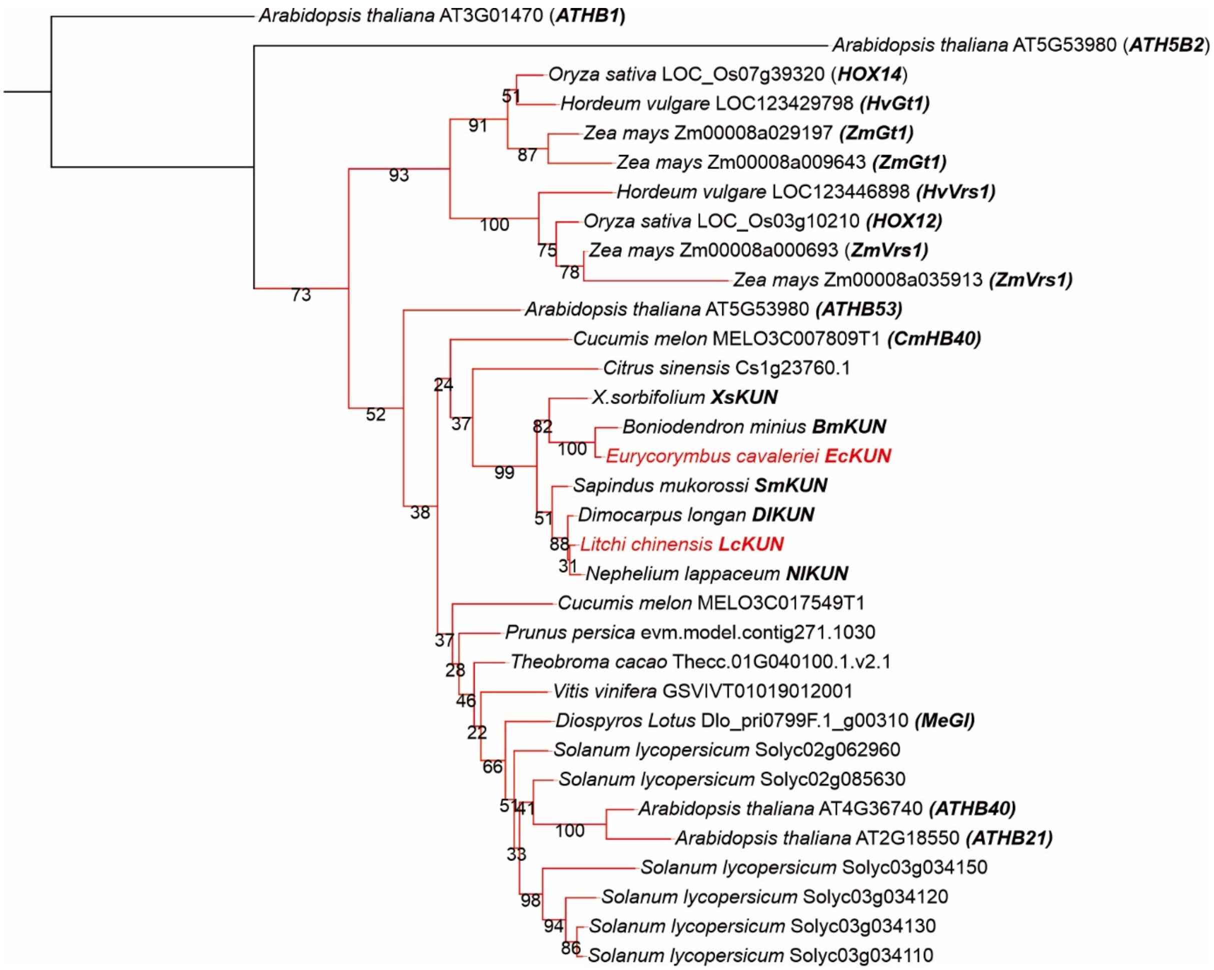


**Fig. S28 Phylogenetic tree of *KUN* with *HD-ZIP I* family genes.**

The colored branches in the phylogenetic tree represent *ATHB53*, *ATHB40*, *ATHB21* in *Arabidopsis* and *KUN* orthologs from 17 species.

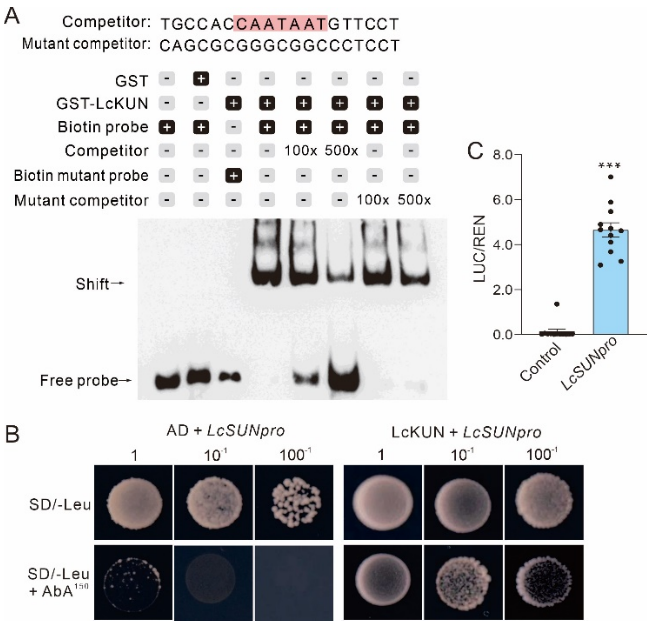


**Fig. S29 LcKUN could activate the expression of *LcSUN* by binding to the promoter region.** **(A)** Electrophoretic mobility shift assay (EMSA) shows that LcKUN directly binds to motifs in the promoter of *LcSUN*. Recombinant purified truncated LcKUN (containing the HD-ZIP I DNA-binding domain) protein (1 µg) was incubated with biotin-labeled probes or with an unlabeled DNA probe containing intact (competitor) or mutated (mutant competitor) binding motifs. **(B)**. The Y1H assay reveals the binding of LcKUN to specific fragments in the promoter of *LcSUN*. pGADT7-*LcKUN* served as the prey, while pAbAi-*LcSUN* was used as the bait. **(C)** LcKUN activated the expression of *LcSUN* in vivo, as demonstrated by transient dual-luciferase reporter assays. Results shown are means ± standard error (n > 10; *** indicates $P \leq 0.001$, Student's *t* tests).

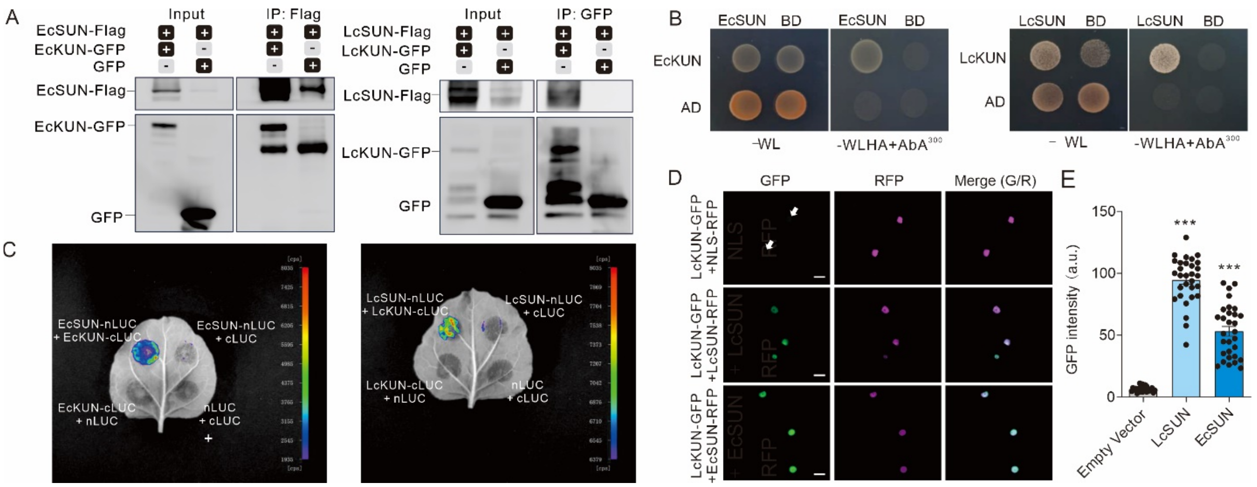


**Fig. S31 Overexpression of *LcKUN* leads to defective anther development in both *Arabidopsis* and tobacco.**

**(A)** The opened flowers from the wild type (WT) (left) and the *35S::LcKUN* transgenic plant (right). Bars = 1 mm. **(B)** Alexander staining of pollen grains from the wild type (left) and the *35S::LcKUN* transgenic Arabidopsis plant (right). Bars = 50 µm. The percentage of abnormal pollen grains is shown at the bottom. Results are means ± standard error (SE, n > 200). **(C)** SEM images of mature pollen grains from the wild type and the *pAtSTK::AtSTK* transgenic line 3. Abnormal pollen grains are indicated by red. Bars = 50 µm. Abnormal pollen grains are highlighted in purple. **(D)** Representative open siliques of the wild type and the *35S::LcKUN* transgenic plant, respectively. Abnormal seeds are indicated by white arrowheads. Bars = 1 mm. **(E)** The ratio of the filament length to the stigma length. WT, wild-type, 35S*::LcKUN*, *Arabidopsis* stable transgenic line overexpressing *LcKUN*. Results shown are means ± standard error (SE, n >6). Asterisks represent significant differences between WT and 35S*::LcKUN* transgenic plants in Student's *t* tests (***, $P \le 0.001$). **(F)** Quantification of seed set from WT and *35S::LcKUN* transgenic plants in *Arabidopsis*. Results shown are means ± standard error (SE, n = 4). Asterisks represent significant differences between WT and *35S::LcKUN* transgenic plants in Student's *t* tests (**, $0.001 < P \le 0.01$). **(G)** The opened flowers from the wild type (WT) and the *35S::LcKUN* transgenic tobacco plant. The middle image shows an enlarged view of the pollen-releasing anthers. Bars = 1 cm. **(H)** Alexander staining of pollen grains from the wild type and the *35S::LcKUN* transgenic tobacco plant. Bars = 50 µm. The percentage of abnormal pollen grains is shown at the bottom. Results are means ± standard error (SE, n > 200). **(I)** The relative expression levels of

*LcKUN* in developing siliques of WT and *35S::LcKUN* transgenic plants in *Arabidopsis*. Results shown are means ± standard error (SE, n >3). Asterisks represent significant differences between WT and *35S::LcKUN* transgenic plants in Student's *t* tests (*, $0.01 < P \leq 0.05$). **(J)** The relative expression levels of *LcKUN* in ovary of wild type and *35S::LcKUN* transgenic tobacco plants. Results shown are means ± standard error (SE, n = 3). Asterisks represent significant differences between WT and *35S::LcKUN* transgenic plants in Student's *t* tests (*, $0.01 < P \leq 0.05$).

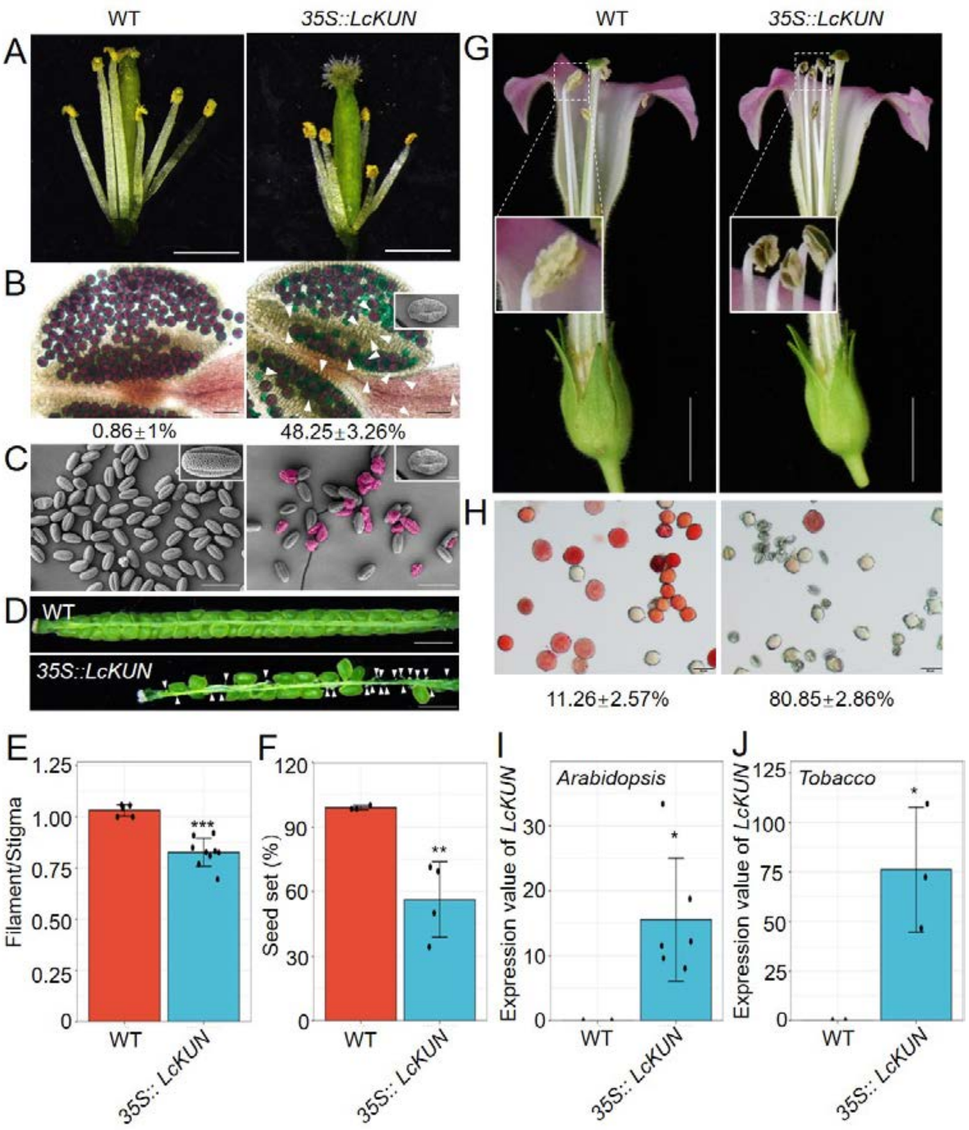


**Fig. S31 Overexpression of *LcKUN* leads to defective anther development in both *Arabidopsis* and tobacco.**

**(A)** The opened flowers from the wild type (WT) (left) and the *35S::LcKUN* transgenic plant (right). Bars = 1 mm. **(B)** Alexander staining of pollen grains from the wild type (left) and the *35S::LcKUN* transgenic Arabidopsis plant (right). Bars = 50 µm. The percentage of abnormal pollen grains is shown at the bottom. Results are means ± standard error (SE, n > 200). **(C)** SEM images of mature pollen grains from the wild type and the *pAtSTK::AtSTK* transgenic line 3.

Abnormal pollen grains are indicated by red. Bars = 50 µm. Abnormal pollen grains are highlighted in purple. **(D)** Representative open siliques of the wild type and the *35S::LcKUN* transgenic plant, respectively. Abnormal seeds are indicated by white arrowheads. Bars = 1 mm. **(E)** The ratio of the filament length to the stigma length. WT, wild-type, 35S*::LcKUN*, *Arabidopsis* stable transgenic line overexpressing *LcKUN*. Results shown are means ± standard error (SE, n >6). Asterisks represent significant differences between WT and 35S*::LcKUN* transgenic plants in Student's *t* tests (***, $P \leq 0.001$). **(F)** Quantification of seed set from WT and *35S::LcKUN* transgenic plants in *Arabidopsis*. Results shown are means ± standard error (SE, n = 4). Asterisks represent significant differences between WT and *35S::LcKUN* transgenic plants in Student's *t* tests (**, $0.001 < P \leq 0.01$). **(G)** The opened flowers from the wild type (WT) and the *35S::LcKUN* transgenic tobacco plant. The middle image shows an enlarged view of the pollen-releasing anthers. Bars = 1 cm. **(H)** Alexander staining of pollen grains from the wild type and the *35S::LcKUN* transgenic tobacco plant. Bars = 50 µm. The percentage of abnormal pollen grains is shown at the bottom. Results are means ± standard error (SE, n > 200). **(I)** The relative expression levels of *LcKUN* in developing siliques of WT and *35S::LcKUN* transgenic plants in *Arabidopsis*. Results shown are means ± standard error (SE, n >3). Asterisks represent significant differences between WT and *35S::LcKUN* transgenic plants in Student's *t* tests (*, $0.01 < P \leq 0.05$). **(J)** The relative expression levels of *LcKUN* in ovary of wild type and *35S::LcKUN* transgenic tobacco plants. Results shown are means ± standard error (SE, n = 3). Asterisks represent significant differences between WT and *35S::LcKUN* transgenic plants in Student's *t* tests (*, $0.01 < P \leq 0.05$).

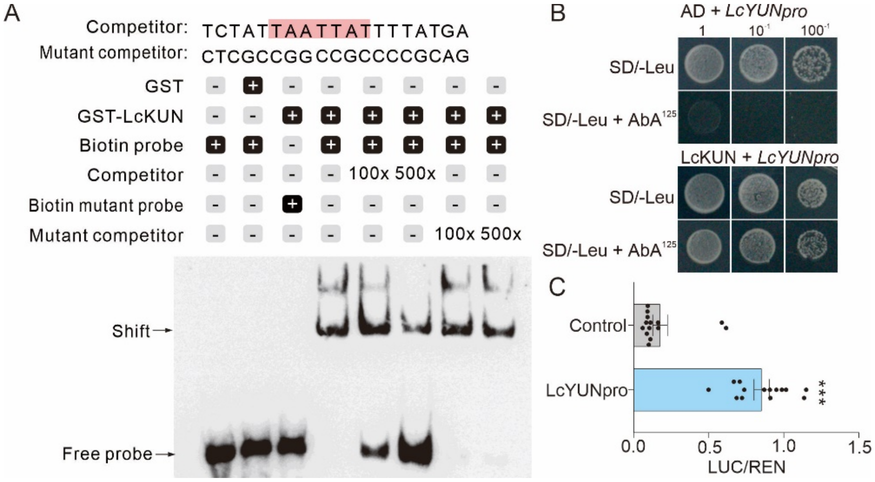


**Fig. S32 LcKUN could activate the expression of *LcYUN* by binding to the promoter region.** **(A)** Electrophoretic mobility shift assay (EMSA) shows that LcKUN directly binds to motifs in the promoter of *LcYUN*. Recombinant purified truncated LcKUN (containing the HD-ZIP I DNA-binding domain) protein (1 µg) was incubated with biotin-labeled probes or with an unlabeled DNA probe containing intact (competitor) or mutated (mutant competitor) binding motifs. **(B)** The Y1H assay reveals the binding of LcKUN to specific fragments in the promoter of *LcSUN*. pGADT7-*LcKUN* served as the prey, while pAbAi-*LcYUN* was used as the bait. **(C)** LcKUN activated the expression of *LcYUN* in vivo, as demonstrated by transient dual-luciferase reporter assays. Results shown are means ± standard error (n > 10; *** indicates $P \leq 0.001$, Student's *t* tests).

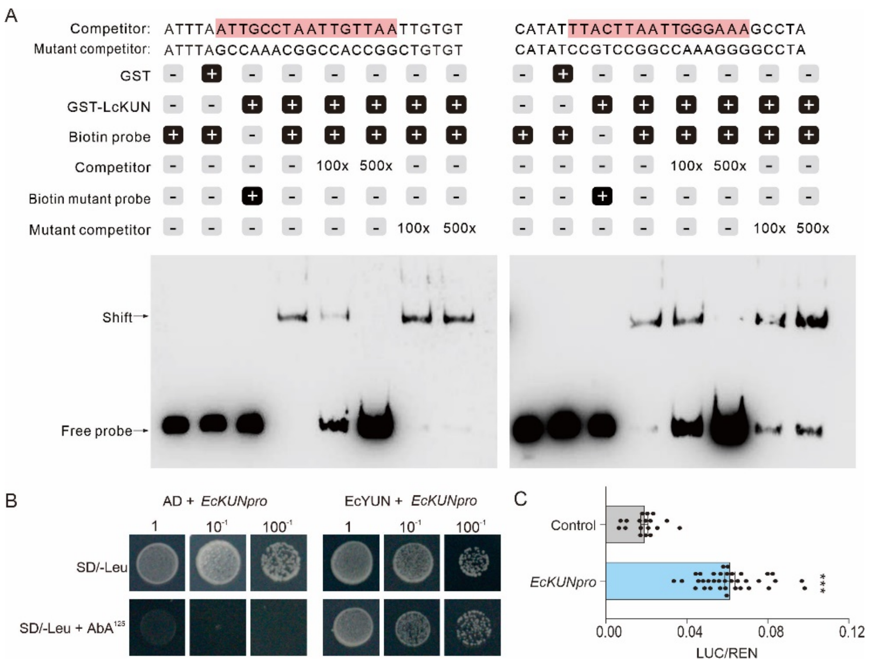


**Fig. S33 EcYUN could activate the expression of *EcKUN* by binding to the promoter region.** **(A)** Electrophoretic mobility shift assay (EMSA) shows direct binding of EcYUN to two distinct motifs in the *EcKUN* promoter. Recombinant purified truncated EcYUN protein (containing the MADS-box DNA-binding domain) (1 µg) was incubated with biotin-labeled probes or with an unlabeled DNA probe containing intact (competitor) or mutated (mutant probe) binding motifs. **(B)** The Y1H assay reveals the binding of EcYUN to specific fragments in the promoter of *EcKUN*. pGADT7-*EcYUN* served as the prey, while pAbAi-*EcKUN* was used as the bait. **(C)** EcYUN activated the expression of *EcKUN* in vivo, as demonstrated by transient dual-luciferase reporter assays. Results shown are means ± standard error (n > 10; *** indicates $P \leq 0.001$, Student's *t* tests).

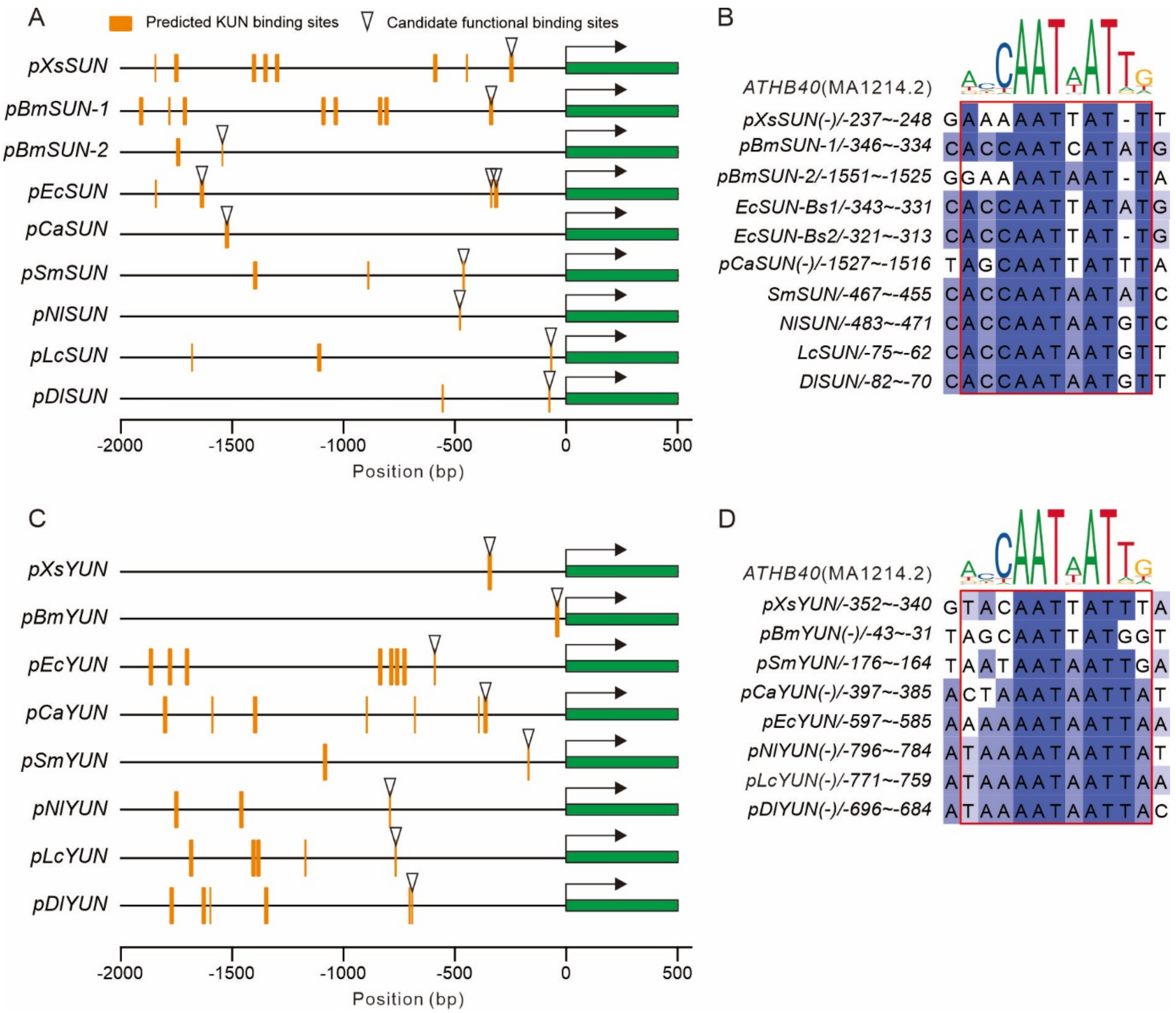


**Fig. S34 Binding sites of KUN in the promoter regions of *SUN* and *YUN* exhibited high conservation across Sapindaceae species.**

**(A)** Predicted binding sites of KUN in the 2000 bp upstream region of *SUN*. **(B)** Sequence alignment of putative functional binding sites of KUN in *SUN*. **(C)** Predicted binding sites of KUN in the 2000 bp upstream region of *YUN*. **(D)** Sequence alignment of putative functional binding sites of KUN in *YUN*.

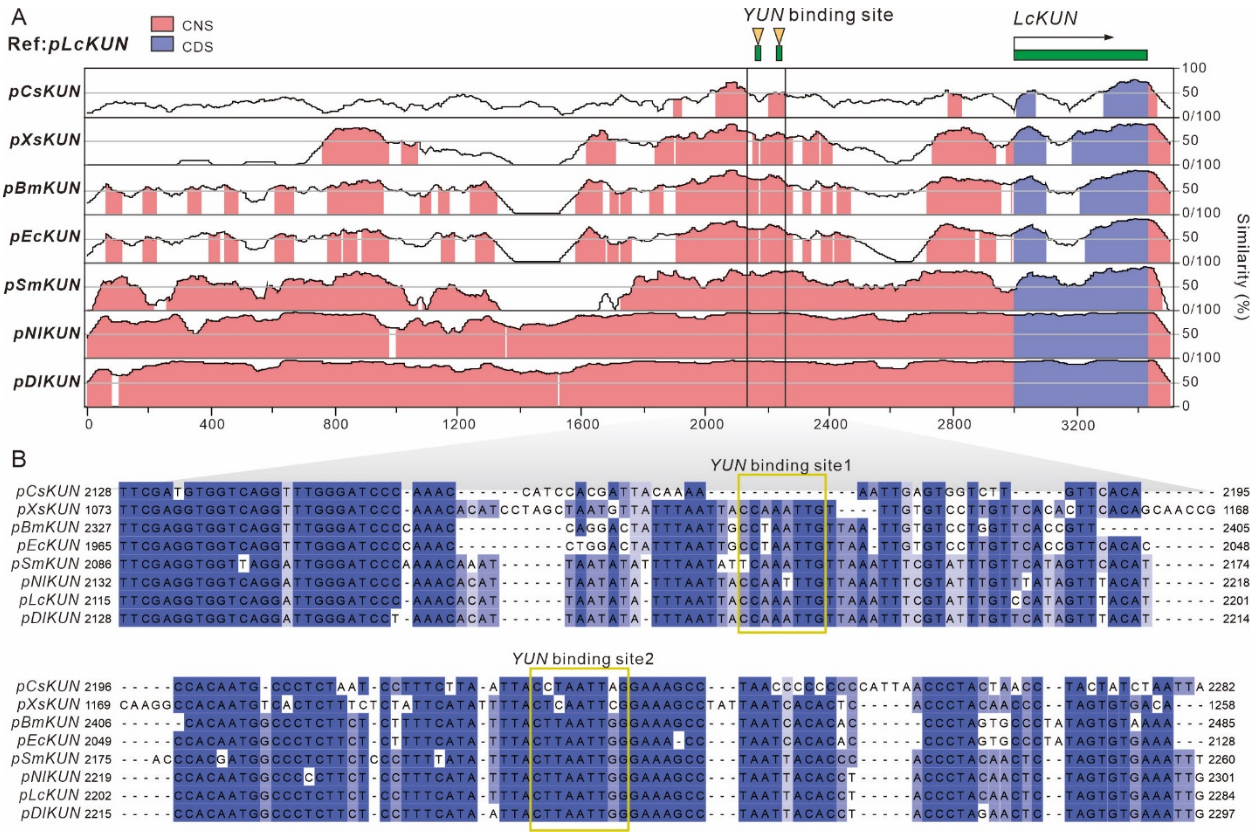


**Fig. S35 Binding sites of YUN in the promoter regions of *KUN* exhibited high conservation across Sapindaceae species.**

**(A)** Conservative non-coding sequence **(**CNS) in the upstream regulatory region of *KUN* across the *Citrus* and seven Sapindaceae species. (**B)** Sequence alignment of the CNS with the putative functional binding sites of YUN.

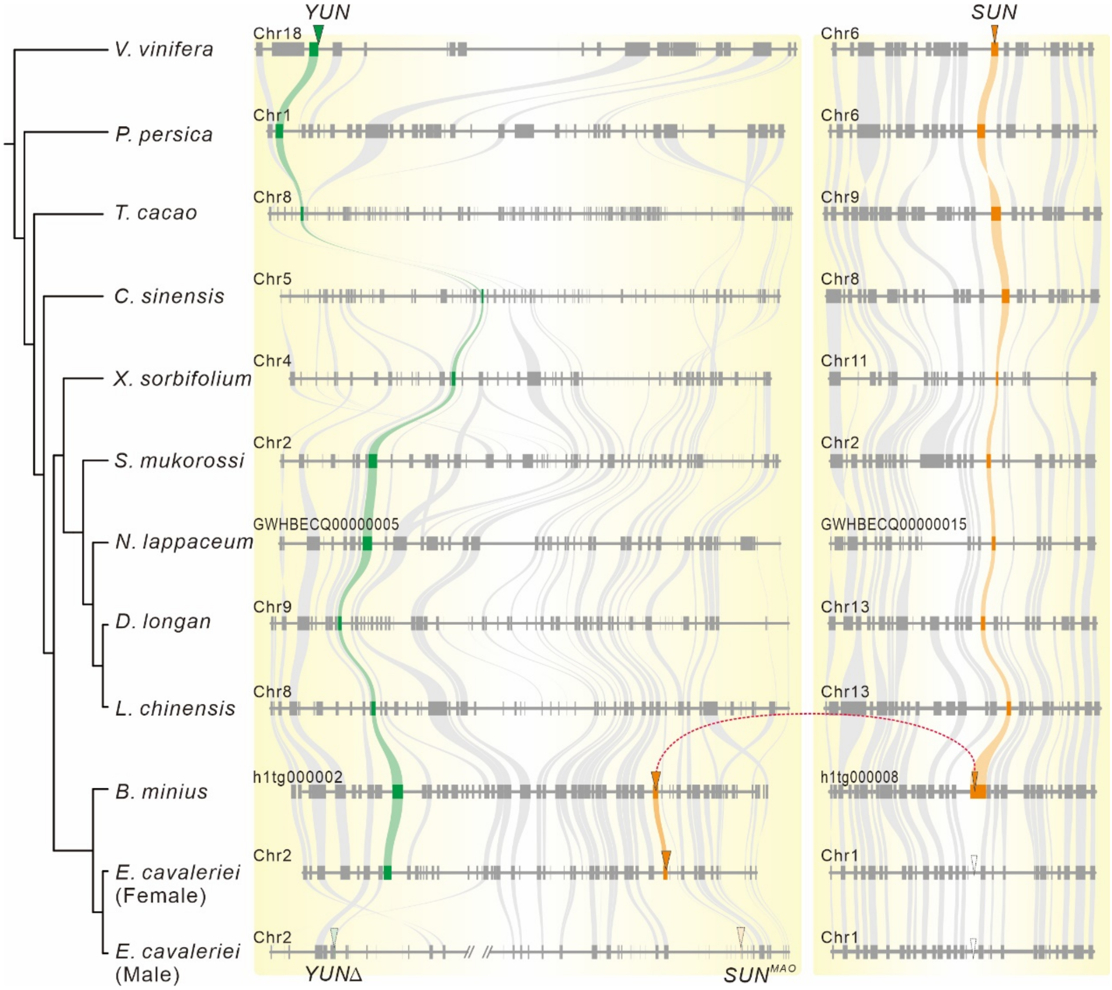


**Fig. S36 Collinearity analysis of *YUN* and *SUN* genes among 12 angiosperm species.**

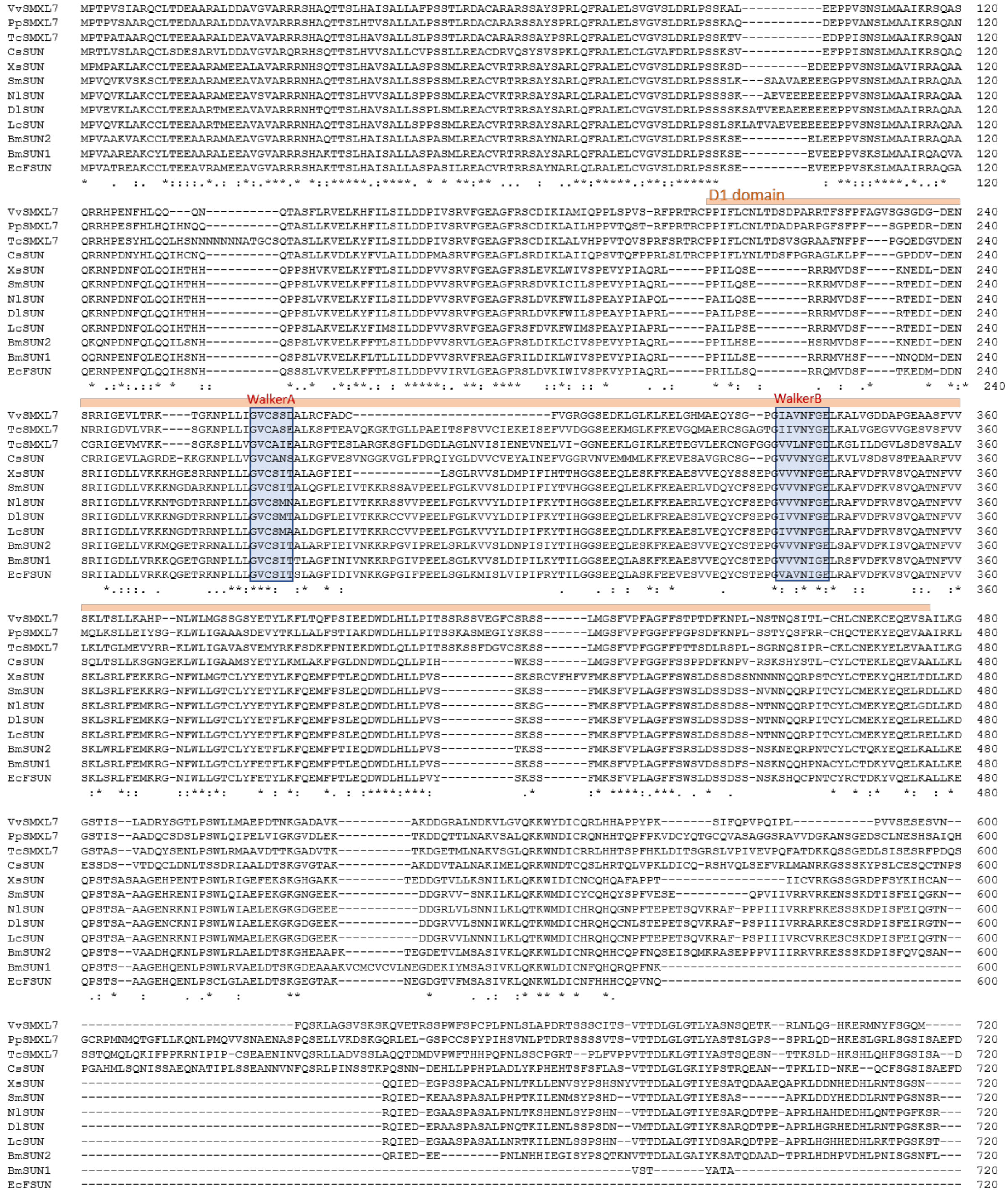
D1 domain
WalkerA
WalkerB

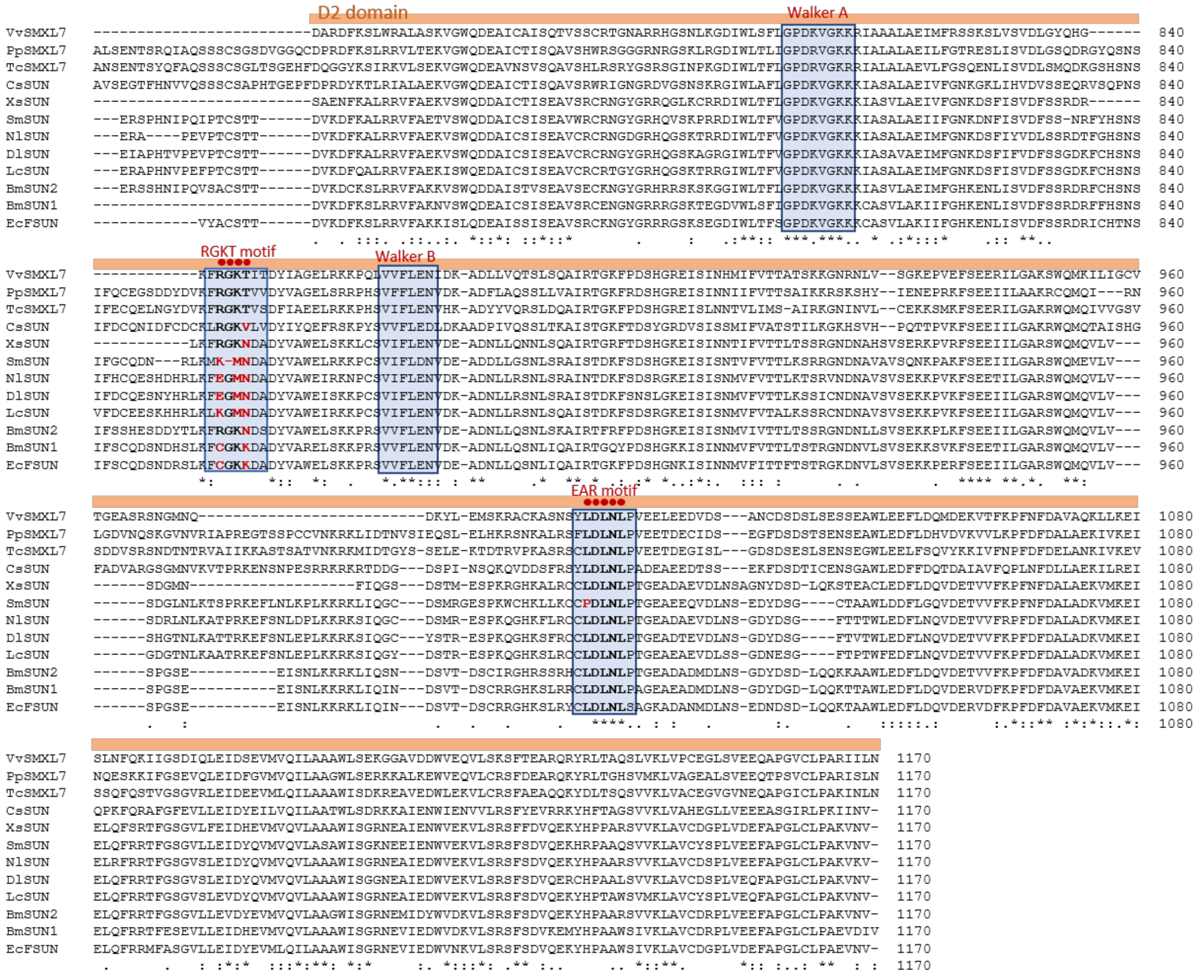


**Fig. S37 Multiple sequence alignment of *SUN* and *SMXL7*.**

SMXL7 proteins contain a D1 ATPase domain, and a D2 ATPase domain. The predicted Walker A (P-loop), Walker B, and EAR motifs are highlighted by light blue box.
